# CD33 and clusterin interact biophysically and genetically to modulate Alzheimer risk

**DOI:** 10.1101/2025.07.29.667318

**Authors:** Roger B. Dodd, Masahiro Enomoto, Ye Zhou, Kanayo Satoh, Yalun Zhang, Fusheng Chen, Beatrice Acheson, Deniz Ghaffari, Elizabeth Sayn-Wittgenstein, Jamie J. Manning, George V. Dukas, Ronak Patel, Alondra Schweizer Burguete, Mason J. Kralovec, Mamunur Rashid, Jennifer L. Hall, Kirstin A. Tamucci, Zena K. Chatila, Minghua Liu, Annie J. Lee, Badri N. Vardarajan, Mariko F. Taga, Sirkku Pollari, Alon Rabinovitch, Cory D. Rillahan, Andrey A. Bobkov, Eduard Sergienko, William Meadows, Seema Qamar, Suzanne J. Randle, Christopher M. Johnson, Jean Sevalle, Jennifer Griffin, Christopher Bohm, Mitsuhiko Ikura, Xunde Xian, Joachim Herz, Megan A. Kelly, Jennifer West, Sandeep Satapathy, Mark R. Wilson, Jonathan A. Javitch, Paul E. Fraser, David A. Bennett, Philip L. De Jager, Zvi Fishelson, Dan Frenkel, Wesley B. Asher, Elizabeth M. Bradshaw, Peter St George-Hyslop

## Abstract

We report the results of structural, functional and genetic studies on the CD33 sialic acid- binding receptor that reveal how non-coding variants in CD33 alter risk for Alzheimer’s disease (AD). The full-length CD33^M^ isoform, whose expression is upregulated by non-coding AD-risk alleles, preferentially forms dimers at the cell surface, where they interact with AD-related proteins (clusterin and Aβ). This interaction induces CD33^M^ inhibitory signalling and downregulates protective microglial functions including phagocytic removal of amyloid plaques. Human brain expression quantitative trait loci (eQTL) and causal mediation analyses confirm that quantitative interactions between CLU and CD33 genotypes modulate AD phenotypes and suggest that genotypes at these loci might be used to personalise future therapeutic approaches. Our work also highlights several other unexpected aspects of CD33 biology, including a soluble shed extracellular fragment of CD33^M^ and a similar soluble secreted product arising from a truncating mutation in the CD33 extracellular domain (CD33^MΔ4bp^).

## INTRODUCTION

The sialic-acid binding immunoglobulin-like lectin 3 receptor (Siglec-3 / CD33) gene encodes a 67 kDa Type I transmembrane protein that is expressed in monocytes and microglia^1,2^. It contains an immunoreceptor tyrosine-based inhibition motif (ITIMs) and an ITIM-like motif in its intracellular cytoplasmic tail. As is the case with many other Siglec receptors, sialylated ligands for CD33 can be offered from the same cell (cis) or from other cells or in solution (trans). Ligands binding to CD33 are thought to induce SRC kinase-mediated tyrosine phosphorylation of the CD33 ITIM domains; recruitment of SH2-contaning phosphatases (SHP-1/SHP-2), and dephosphorylation of downstream targets (e.g. spleen tyrosine kinase - SYK), which suppresses SYK signalling activity. This cascade serves as a negative regulator of protective innate immune and microglial responses^3–9^. CD33 is expressed as two isoforms arising from alternative splicing of Exon 2 (Uniprot ID P20138; Figure 1a)^10,11^. The full-length CD33^M^ isoform contains: i) a putative sialic acid binding site within the V-set domain encoded by Exon 2, and ii) a C2-set immunoglobulin-like (Ig- like) domain (Figure 1a)^12–16^. The truncated CD33^m^ isoform (also known as CD33^ΔE2^) contains only the C2-set domain, with no ligand binding site (Figure 1a).

**FIGURE 1.**
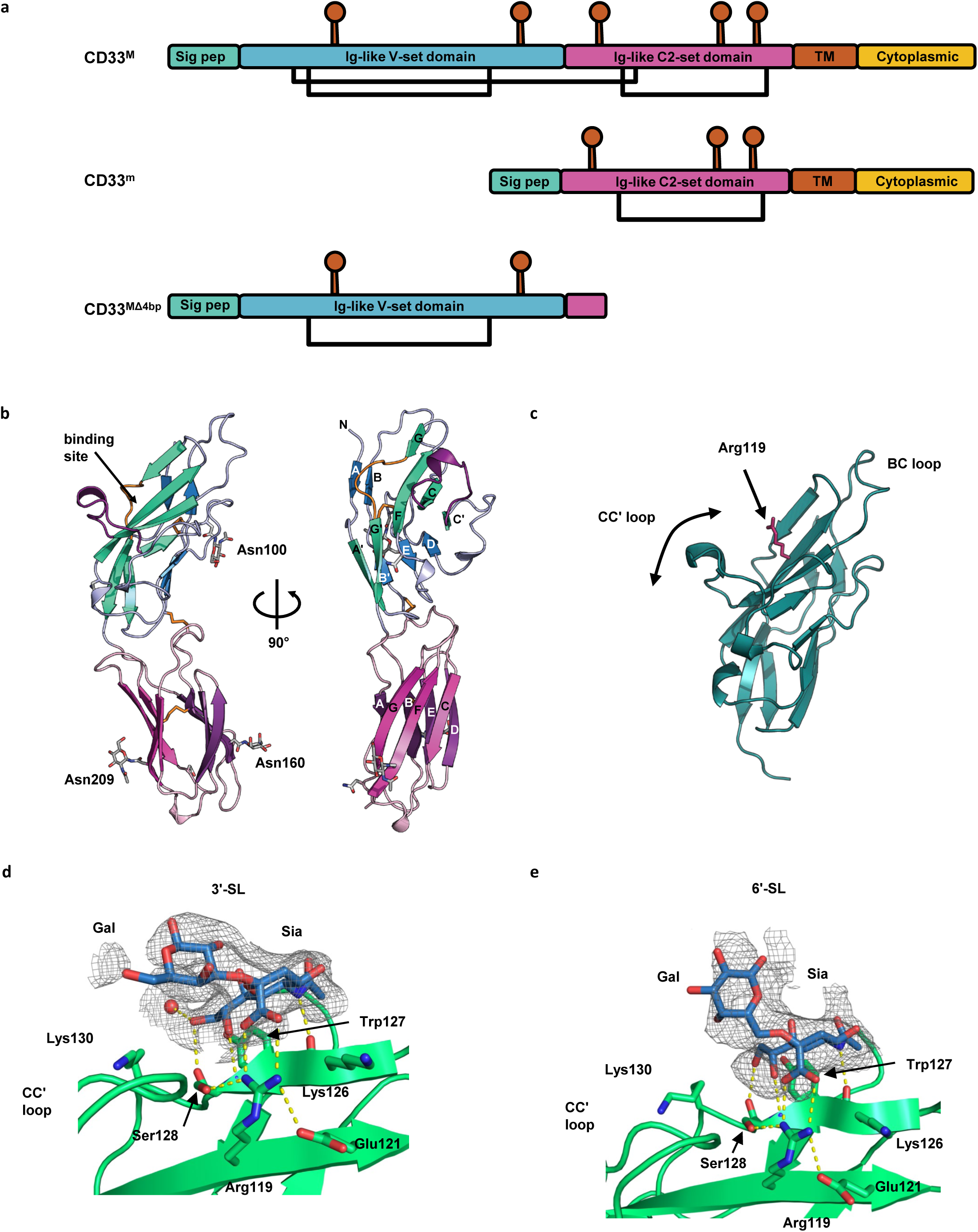
Structure of the CD33 extracellular domain. a. A schematic of CD33/Siglec-3 long (CD33^M^)(*Top*), short (CD33^m^) (*Middle*) and CD33^MΔ4bp^ (*Bottom*) isoforms indicating signal peptide (sig pep, green), two Ig-like domains (blue, purple), transmembrane helix (red) and cytoplasmic tail (yellow). Glycosylation sites are indicated by red lollipops and disulphide bonds by black lines connecting Cys residues. b. Cartoon of the CD33^M^ ECD structure in two views rotated 90° about the vertical axis. The sheets in the V-set domain are green (C′CFGG′A) and blue (ABED), and the C2-set domain pink (CFG) and purple (ABED). The variable CC′ and GG′ loops are coloured purple and orange, respectively. The V-set domain is stabilised by a disulphide bond between the BBʹ loop (Cys 41) and the E strand (Cys 101). The three *N*-linked GlcNAc residues attached to the V-set domain (Asn100) and C2-set domain (Asn160 and 209) are shown in stick form and labelled by the amino acid to which they are linked c. A cartoon of the CD33 V-set domain structure highlighting important features for binding, and regions of difference to other siglec structures. Arg119, required for binding sialic acid, is shown in stick form (magenta). Regions of significant deviation to other siglecs are limited to the BC loop and the CC’ loop. The CC‘ loop occupies conformations varying from more open (CD33^M^) to closed (Siglec-5) over the binding site. d. Images of the ligand binding sites with bound 3′-sialyllactose. The polypeptide chain is shown in cartoon representation with the residues constituting the binding site in stick form. The sialyllactoses are shown in ball-and-stick format (carbon atoms – light blue, oxygen – red, nitrogen – dark blue) with omit *mFo-DFc* electron density contoured at 2 σ. The electron density maps were supportive of inclusion of sialic acid and Gal residues, but insufficient to model Glc. The CC′ loop is indicated and, in contrast to other Siglecs, makes no direct interactions with the ligands. e. Images of the ligand binding sites with bound 6′-sialyllactose.

Multiple genetic association studies have implicated CD33 as a risk modifying factor for late-onset Alzheimer’s Disease (AD) (meta-analysis odds ratio = 0.94; 95% confidence interval 0.91– 0.96; p = 3.0 × 10^−6^) ^17–27^. The sequence variants conferring altered risk are non-coding, single nucleotide polymorphism (SNP) alleles (rs3865444) that affect both the absolute protein level of expression of CD33 and the relative splicing of exon 2. The major allele (“reference“ or “C”) is associated with increased total levels of CD33 expression and increased relative expression of the long CD33^M^ splice form^10,28–30^. The minor (“A”) allele is associated with lower levels of CD33 expression and increased relative expression of the short CD33^m^ splice form^10,28–30^. *In vitro*, *ex vivo* and *in vivo* functional studies reveal that human major/risk allele carriers have: i) impaired uptake of dextran beads and amyloid β-peptide 1-42 (Aβ) by cultured *ex vivo* peripheral blood mononuclear macrophage cells (monocytes), and ii) increased *in vivo* brain Aβ levels as measured by amyloid positron emission tomography imaging^6–10,28–31^. Intriguingly, the functional effects of CD33^M^ expression (as measured by in vivo brain Aβ levels) were found to be indistinguishable in heterozygous and homozygous risk allele carriers^10,31^. However, there has been some debate as to whether the molecular outcome of the genetic variance that drives the disease association is due to increased CD33^M^, reduced CD33^m^, or both. Some studies suggest that the longer CD33^M^ isoform has a dominant immune-inhibitory phenotypic effect, while others suggest that the CD33^m^ isoform has a dominant protective effect^10,28,30^. These various viewpoints are all supported by findings in murine BV2 cells, primary murine microglia, human U937, human THP-1 and human CHME3 microglial-like cells with recombinant expression models^6–9^. Therefore, more work is required to understand the molecular basis of the dominant immuno-inhibitory phenotypic effect of CD33^M^ seen in primary human cells and in vivo, as well as the identity of potential ligands that might activate immuno-inhibitory CD33^M^ signalling are unknown.

To explore these questions, we solved the structure of the CD33^M^ extracellular domain (ECD). We applied a hypothesis-driven approach to identify potential ligands by investigating how they affect CD33 ITIM inhibitory signalling. We show here the 2.24 Å structure of the extracellular domain of the CD33^M^ isoform. We demonstrate that CD33^M^ forms dimers that are selectively trafficked to the cell surface. We identify clusterin (CLU) and CLU plus Aβ 1-42 oligomers, but not ApoE, as candidate ligands. These ligands do not bind either CD33^M^ monomers or CD33^m/m^ dimers. We demonstrate that binding of CLU with or without Aβ 1-42 oligomers (hereafter CLU ± Aβ oligomers) to CD33^M^ activates CD33 ITIM signalling. We show in *ex vivo* cortical slices from APP transgenic mice that activation of CD33^M^ immuno-inhibitory signalling impairs clearance of amyloid plaques by human monocyte-like cells endogenously expressing CD33^M^ isoforms. Finally, we report our discovery of a previously unrecognised soluble ectodomain fragment of CD33^M^ analogous to the sTREM2 fragment of TREM2. We also demonstrate that a rare CD33 frame shift mutant (CD33^MΔ4BP^), which was previously thought to be a null mutant, produces a stable soluble extra-cellular domain protein, similar to sCD33^M^. Further work is required to determine if sCD33^M^ and CD33^MΔ4BP^ have biological activity. Taken together these observations suggest a molecular basis for the dominant effect of the major “C” CD33 risk allele that favours CD33^M^ dimer formation. They also raise the possibility that therapeutic down-regulation of CD33^M^ signalling could be achieved by several mechanisms, including selective disruption of the CD33 dimer interface.

## RESULTS

### Overall structure of the CD33^M^ ectodomain

To solve their structures, we expressed the CD33^M^ (residues 21-232) and CD33^m^ ectodomains in HEK293F cells grown in the presence of kifunensine, a mannosidase inhibitor, to render glycans Endoglycosidase H (Endo H)-sensitive. The ectodomains were purified by affinity and gel filtration chromatography, treated with Endo H, and crystallised. We were unable to generate high quality crystals for the protective CD33^m^ isoform lacking the V-set domain. However, its structure can be inferred from that of the CD33^M^ isoform.

The CD33^M^ ECD crystallises in an orthorhombic space group with four chains (A, B, C, D) in the asymmetric unit that differ only very slightly in the conformation of two flexible loops (average RMSD of 1.3 Å; Table 1). Unless stated otherwise, the following text refers to chain B. The CD33^M^ ECD adopts a two-domain fold (Figure 1b, Supplementary Data Figure 1A-C) composed of an N- terminal V-set domain (residues 21-139) and a C2-set domain (residues 143-232). The two domains are joined by a short linker region (residues 140-142) and are related by a hinge angle of 135°. An intramolecular disulphide bond between Cys36 and Cys169 stabilises the interdomain interface (Supplementary Data Figure 1A).

**TABLE 1.**
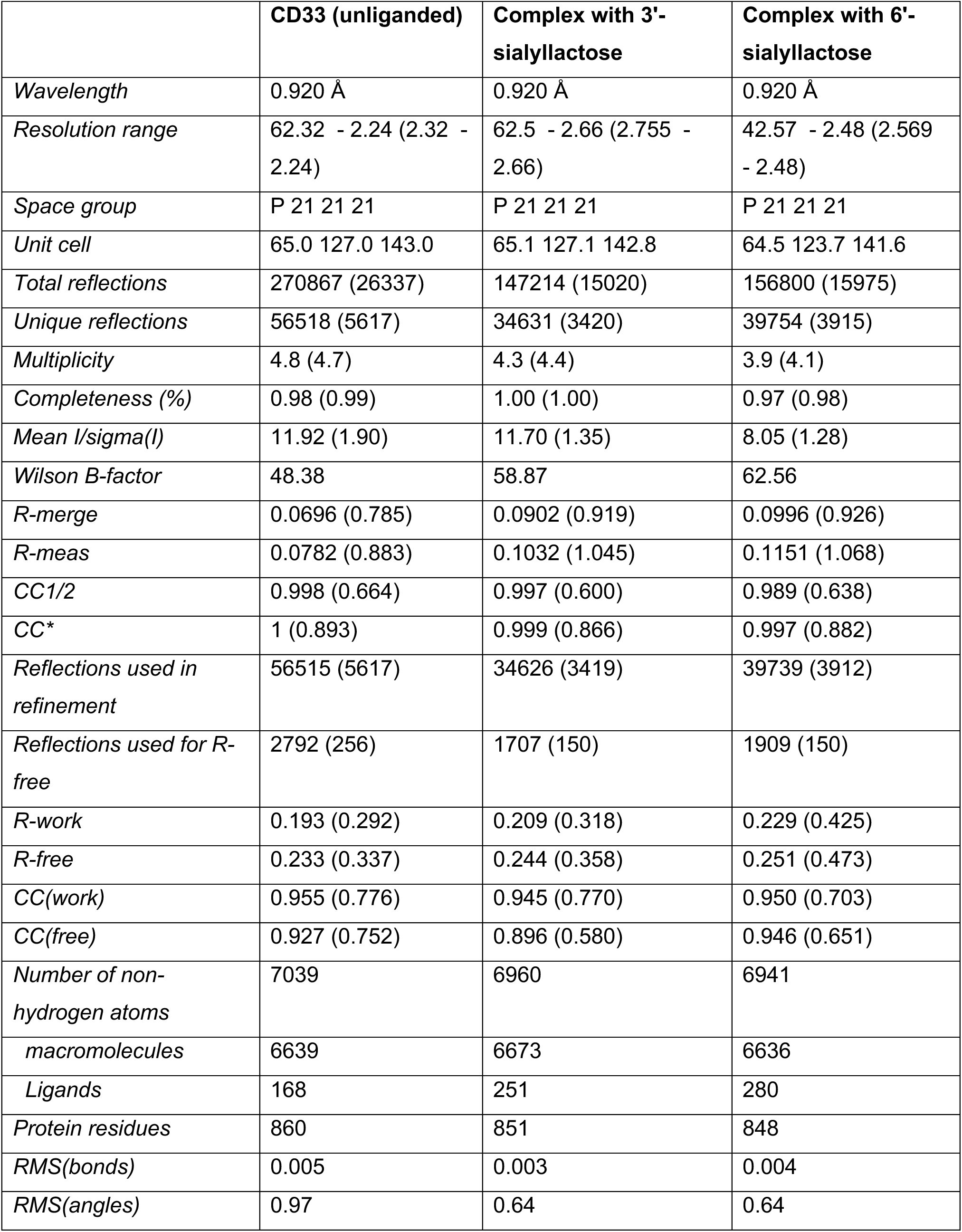
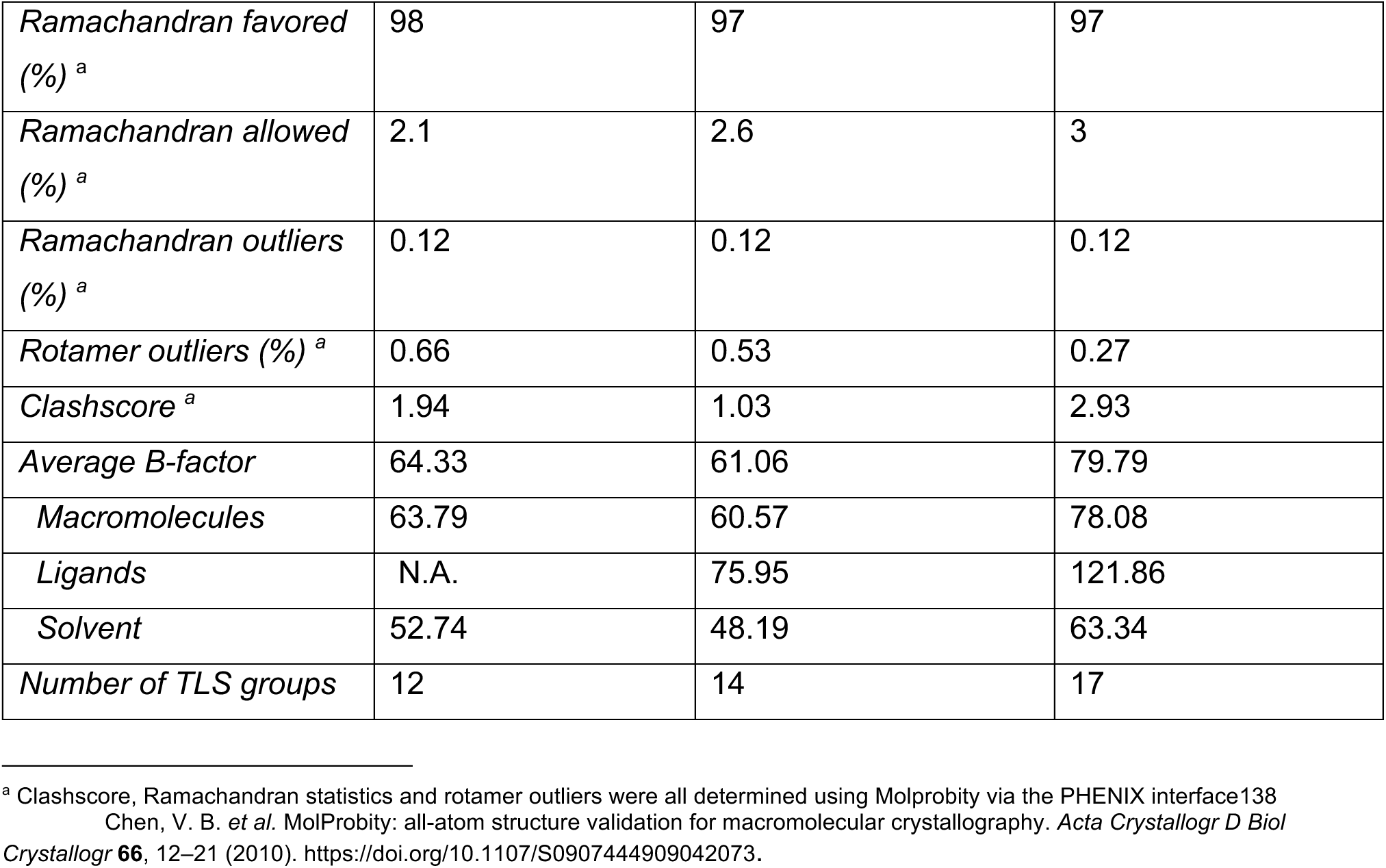

|  | **CD33 (unliganded)** | **Complex with 3'-sialyllactose** | **Complex with 6'-sialyllactose** |
| --- | --- | --- | --- |
| *Wavelength* | 0.920 Å | 0.920 Å | 0.920 Å |
| *Resolution range* | 62.32 - 2.24 (2.32 - 2.24) | 62.5 - 2.66 (2.755 - 2.66) | 42.57 - 2.48 (2.569 - 2.48) |
| *Space group* | P 21 21 21 | P 21 21 21 | P 21 21 21 |
| *Unit cell* | 65.0 127.0 143.0 | 65.1 127.1 142.8 | 64.5 123.7 141.6 |
| *Total reflections* | 270867 (26337) | 147214 (15020) | 156800 (15975) |
| *Unique reflections* | 56518 (5617) | 34631 (3420) | 39754 (3915) |
| *Multiplicity* | 4.8 (4.7) | 4.3 (4.4) | 3.9 (4.1) |
| *Completeness (%)* | 0.98 (0.99) | 1.00 (1.00) | 0.97 (0.98) |
| *Mean I/sigma(I)* | 11.92 (1.90) | 11.70 (1.35) | 8.05 (1.28) |
| *Wilson B-factor* | 48.38 | 58.87 | 62.56 |
| *R-merge* | 0.0696 (0.785) | 0.0902 (0.919) | 0.0996 (0.926) |
| *R-meas* | 0.0782 (0.883) | 0.1032 (1.045) | 0.1151 (1.068) |
| *CC1/2* | 0.998 (0.664) | 0.997 (0.600) | 0.989 (0.638) |
| *CC\** | 1 (0.893) | 0.999 (0.866) | 0.997 (0.882) |
| *Reflections used in refinement* | 56515 (5617) | 34626 (3419) | 39739 (3912) |
| *Reflections used for R-free* | 2792 (256) | 1707 (150) | 1909 (150) |
| *R-work* | 0.193 (0.292) | 0.209 (0.318) | 0.229 (0.425) |
| *R-free* | 0.233 (0.337) | 0.244 (0.358) | 0.251 (0.473) |
| *CC(work)* | 0.955 (0.776) | 0.945 (0.770) | 0.950 (0.703) |
| *CC(free)* | 0.927 (0.752) | 0.896 (0.580) | 0.946 (0.651) |
| *Number of non-hydrogen atoms* | 7039 | 6960 | 6941 |
| *macromolecules* | 6639 | 6673 | 6636 |
| *Ligands* | 168 | 251 | 280 |
| *Protein residues* | 860 | 851 | 848 |
| *RMS(bonds)* | 0.005 | 0.003 | 0.004 |
| *RMS(angles)* | 0.97 | 0.64 | 0.64 |
| *Ramachandran favored (%)*<sup>a</sup> | 98 | 97 | 97 |
| *Ramachandran allowed (%)*<sup>a</sup> | 2.1 | 2.6 | 3 |
| *Ramachandran outliers (%)*<sup>a</sup> | 0.12 | 0.12 | 0.12 |
| *Rotamer outliers (%)*<sup>a</sup> | 0.66 | 0.53 | 0.27 |
| *Clashscore*<sup>a</sup> | 1.94 | 1.03 | 2.93 |
| *Average B-factor* | 64.33 | 61.06 | 79.79 |
| *Macromolecules* | 63.79 | 60.57 | 78.08 |
| *Ligands* | N.A. | 75.95 | 121.86 |
| *Solvent* | 52.74 | 48.19 | 63.34 |
| *Number of TLS groups* | 12 | 14 | 17 |

<sup>a</sup> Clashscore, Ramachandran statistics and rotamer outliers were all determined using Molprobity via the PHENIX interface Chen, V. B. *et al.* MolProbity: all-atom structure validation for macromolecular crystallography. *Acta Crystallogr D Biol Crystallogr* **66**, 12–21 (2010). https://doi.org/10.1107/S0907444909042073.

The V-set domain contains a β-sandwich formed by the juxtaposition of two β-sheets stabilized by the Cys41-Cys101 disulphide bond (Figure 1a-c) and has a similar overall conformation to those of sialoadhesin^5^, Siglec-5^8^ and Siglec-7^4^ (Supplementary Data Figure 1B). The C2-set domain is comprised of two β-sheets forming a sandwich stabilised by the Cys163- Cys212 disulphide bond (Figure 1a,b). The structure of the C2-set domain differs from that of Siglec-5 by absence of a Cʹ strand and by formation of an additional D-strand abutting the E-strand to yield a 4-stranded ABED β-sheet, rather than loops with two single-turn helices (Supplementary Data Figure 1C).

Electron densities reflecting *N*-linked GlcNAc residues were observed at Asn100, Asn160 and Asn209 (Supplementary Data Figure 1D), with weaker densities suggestive of partial glycosylation at Asn113 and Asn230. The Asn100 residue, which has been previously implicated in auto-inhibition of ligand binding^32^, is located on the opposite site of the CD33 chain to the ligand binding site on the same chain.

The structure of the sialic acid ligand binding site of CD33^M^ in both unbound apo-form, and ligand-bound forms complexed with 3ʹ-sialyllactose (3ʹ-SL) or with 6ʹ-sialyllactose (6ʹ-SL), were solved at 2.66 Å and 2.48 Å, respectively (Figures 1c-e; Supplementary Data Figure 1D). The ligand binding site contains the principal elements of canonical Siglec ligand-binding sites^12,15,16^.

These canonical features include: i) an invariant arginine in the F strand (Arg119, the guanidino group of which engages with the sialic acid); and ii) elements contributed from the G strand, GGʹ and the more heterogeneous CCʹ loops (Figure 1c-e, Supplementary Data Figures 1D,E). The structure of the ligand binding site reported here is in good agreement with three other CD33 V-set domain structures in the Protein Data Bank (PDB 6D48, 6D49, 6D4A ^33^), an unpublished CD33^M^ structure (7AW6), and an AlphaFold2 model (RMSD in the range of 0.3-0.4 Å, Supplementary Data Figure 1B, panels i-iii). We were interested to determine if the sialyllactose complex structures could explain contradictory published observations regarding CD33 substrate specificity^34,35^. However, we were surprised to find that no direct interactions were made with the sub-terminal sugar residues, with both α2-3 and α2-6 linked glycans equally accommodated (Figure 1d,e, Supplementary Data Figure 1D). The lack of glycan preference is in agreement with the observed promiscuity in binding various sialylated glycans. The data suggest substrate preference is likely specified by the precise chemical nature, density and spatial disposition of glycans on CD33- binding proteins. An analogous situation has been suggested previously for Siglec-5^16^.

A comparison of the CD33^M^ structure to those of other Siglec family members indicates that the CD33^M^ ligand binding site differs in three ways (Figure 1c, Supplementary Data Figure 1B, panels i-iii). First, the guanidino group of Arg119 of CD33 is pre-positioned to interact with the carboxylate group of the sialic acid ligand (Figure 1c, Supplementary Data Figure 1E,F). In other Siglecs (e.g. Siglec-5), the guanidino group of the canonical central arginine must undergo a 180° rotation to engage substrates. Second, in CD33^M^, two lysine residues (Lys126 in the G strand, and Lys130 in the GG’ loop) form an electrostatically favourable surface for binding of negatively charged sugar residues (Figure 1d, Supplementary Data Figure 1B-F). In contrast, in other Siglecs, the lysines equivalent to CD33 Lys130 block access to the ligand binding site. Third, the CCʹ loop of CD33 does not directly interact with ligand, whereas the equivalent loops in other Siglecs directly bind ligand (e.g. via Tyr68 in Siglec-5) (Figure 1c-e, Supplementary Data 1B-F). However, the CCʹ loop may be able to adopt multiple conformations to allow binding to a range of sialic acid-containing ligands of difference sizes and configurations – including, as described below, much larger sialylated molecules.

Other Siglecs for which structures have been solved do not show explicit evidence of dimerization. However, unexpectedly, the crystal structure reveals that CD33^M^ ECD forms asymmetric dimeric quaternary associations (Figure 2a; dimer pairs are A:D and B:C). There is no apparent mechanism revealed by the crystal structure for higher order oligomerisation (e.g. nanoclusters) beyond the dimeric state. However, it is possible that such nanoclusters could be driven indirectly by highly glycosylated ligands. The following text describes the A:D dimer in detail. This interaction occurs via a series of residues that are conserved (>60% identity) in the C2-set domains of 105 diverse Siglecs from multiple species (Supplementary Data Figure 1G), and is predicted by the Evolutionary Protein Protein Interface Classifier (EPICC)^36^ algorithm as being an authentic biological interface (P_bio_ = 1.0). The dimer interface is comprised of an 843 Å^2^ surface generated predominantly by the C and D strands (Figures 2a,b). The dimer is stabilised by: i) main chain inter-strand hydrogen bonds between residues 176-181 (Supplementary Data Figure 1H); ii) hydrophobic side chain interactions between residues Ile176, Phe177, Trp179, Leu180 and Leu187 (Supplementary Data Figure 1I); and iii) polar interactions from residues surrounding the hydrophobic core, including: Arg190A to Ser178D; Gln213D to Gly188A; and Lys215D to Arg190A (Supplementary Data Figure 1J). We note that the dimeric interaction described here is also observed in the unpublished 7AW6 CD33^M^ structure, crystallised in a different crystal lattice, supporting our observations (Supplementary Data Figure 1B, panel iii). We cannot fully discount the possibility that the observed asymmetry, which is unusual, was imposed by the crystalline packing arrangement, and that the dimer is actually symmetric. Nevertheless, *bona fide* asymmetric dimers are known (e.g. α-catenin^37^ and EGFR^38^).

**FIGURE 2.**
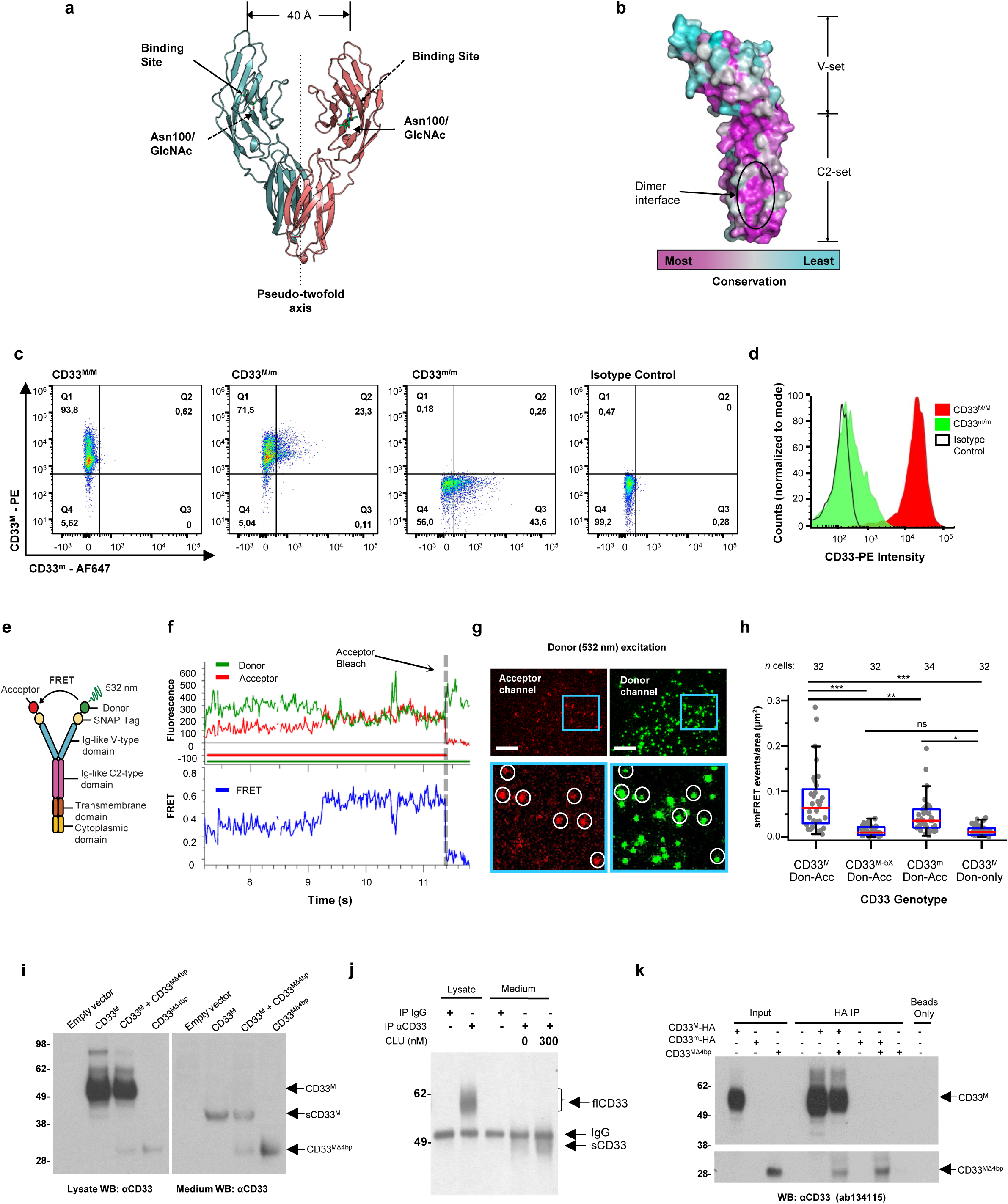
CD33 forms dimers. a. **Cartoon showing the dimerization of two CD33^M^ ectodomain chains** (teal and pink) mediated by their C2-set domains. The two C strands hydrogen bond in a parallel fashion to create a continuous β-sheet (GFCCFG) and the remainder of the interface is formed by residues from the CD33-specific D strand. The C2-set dimeric interface and the rigid nature of the V-set/C2-set interface places the two ligand binding sites 40 Å apart, facing outwards from the central axis of the dimer and pointing away from one another. b. **Surface representation of CD33^M^ ectodomain colour-coded according to conservation** (purple – conserved, cyan – non-conserved). The conserved patch constituting the dimer interface is indicated. c. **Dimeric CD33^M^ is robustly targeted to the cell surface in both CD33^M^/CD33^M^ and CD33^M^/CD33^m^ expressing cells**: representative flow cytometry analyses showing CD33^M^ and CD33^m^ surface labelling in cells expressing CD33^M^ only (CD33^M^/^M^), CD33^m^ only (CD33^m^/^m^) or CD33^M^ and CD33^m^ (CD33^M^/^m^), reveals that the majority of both CD33^M^/^M^ and CD33^M^/^m^ cells have similarly robust surface exposure of CD33^M^. We were not able to include CD33^M-5X^ cells in these experiments because the 5X dimer site mutant disrupts the recognition motif for the anti-CD33 antibody that recognizes CD33^M^ in fluorescence flow cytometry. Cells were double stained using antibodies specific to CD33^M^ (WM53) and CD33^m^ (A16121H) respectively shows, in the displayed experiment (see Figure 2c), that most CD33^M^/^M^ cells (upper left, Q1+Q2: 93.8% + 0.62%) or CD33^M^/^m^ cells (upper right, Q1+Q2: 71.5% + 23.3%) are surface CD33^M^ positive. Only a small portion of cells in CD33^M^/^m^ cells (upper right, Q2+Q3: 23.3% + 0.11%) are CD33^m^ positive. In the CD33^m^/^m^ (lower left, Q2+Q3: 0.25% + 43.6%) cells, CD33^m^ positive cells are less than a half of the total cells. Figures show one experiment out of three independent experiments (each with 6 technical replicates) with similar results. The MFI statistics of CD33M and CD33m from three independent experiments are shown in Extended Figure 2J d. Single staining using the HIM3-4 antibody, which recognises both CD33^M^ and CD33^m^, shows much greater surface exposure of CD33^M^ in CD33^M^/^M^ cells (red) than there is surface exposure of CD33^m^ in CD33^m^/^m^ (green) cells. e. **Schematic of single-molecule TIRF imaging of CHO cells expressing SNAP_f_-CD33^M^ labeled with donor and acceptor fluorophores**. The colours on the dimer cartoon reflect the same CD33 domains as in Figure 1a. f. **Representative smFRET trajectory and its corresponding fluorescence and FRET time traces for the CD33^M^ receptor (bottom panel).** The smFRET trajectory for the individual molecule is shown to the left of its fluorescence (donor and acceptor trajectories and intensities are shown in green and red, respectively) and FRET trace (in blue). The green and red bars along the time axis in the fluorescence time trace plot indicate that the signal was derived during tracking. g. Top panels: **Representative image from a movie (frame 2, 0.03 s) of labeled SNAP_f_- CD33^M^ e**xcited by the donor laser (532 nm), with the enlarged view (bottom panels) showing sensitized acceptor signals that have colocalizing trajectories with their corresponding donors, both delineated by white circles. (See cartoon in Supplementary Figure 2K for methodology) Scale bar, 5 μm; enlarged view, 9.1 μm × 7.3 μm. Donors without colocalized acceptor are consistent with the stochastic labeling approach employed in which a population of dimers labeled with two donors or only one donor and no acceptor is possible. h. **Distributions of smFRET events per cell area** for SNAP_f_-CD33^M^, -CD33^M-5X^, and -CD33^m^, all labeled with donor and acceptor (Don-Acc) as well as SNAP_f_-CD33^M^ labeled with only donor (Don-only). The smFRET events represent the total number of freely diffusing smFRET trajectories (see cartoon in Supplementary Figure 2K for methodology). Dots represent smFRET events per area for each cell. Box plots indicate the median (value shown as the central line) and interquartile range (lower and upper lines represent the 25th and 75th percentiles, respectively), while the whiskers represent the points that fall within 1.5 × interquartile range. The indicated total number of cells (*n* cells) were collected over 4 independent experiments for the CD33^M^ and CD33^m^ samples, and 3 independent experiments for the CD33^M-5X^ and CD33^M^ donor only samples. ***, p = 1.3 × 10^-8^; ***, p = 1.6 × 10^-8^; **, p = 0.0056; *, p = 0.011; ns = 0.99; ordinary one-way ANOVA with Šídák’s post- hoc multiple comparisons test. i. **CD33^MΔ4bp^ variant forms a stable soluble protein in cell lysates and is secreted into the media**, where it may bind with the extracellular domains of CD33^M^ and CD33^m^. There may be a similar physiological soluble CD33 protein that is either shed into the medium by CD33^M^ ectodomain proteolysis or is produced as an alternative splicing product. A cartoon showing the domain structures and epitope tags of CD33^M^, CD33^m^, CD33^Δ4bp^ and the putative soluble CD33^M^ ectodomain product (sCD33^M^) is included in Supplementary Data Figure 2M. j. **Western blot analysis of CD33 immunoprecipitated by the CD33 Ig-like V-set domain antibody (ab134115) from lysates and conditioned medium of iMGs**. Immunoprecipitation products were probed for CD33 using the CD33 Ig-like C2-set domain antibody (PWS44). The data demonstrate the presence of fuzzy, likely glycosylated soluble CD33- immunoreactvive (sCD33) cleavage product in the medium, but not in iMG cell lysates. Notably, sCD33 increases upon stimulation of the iMG cells with 300 nM CLU (lane 5). k. **Western blot of lysates and HA-immunoprecipitation products from HEK293 cells expressing CD33^M^-HA, CD33^m^-HA and CD33^MΔ4bp.^** HA-immunoprecipitation products were probed with the anti-CD33 antibody (ab134115) which recognises only epitopes in the V-set domain. Arrowheads denote CD33 derivative protein. Representative blots for n = 3 independent biological replicates. The result reveals that CD33^MΔ4bp^ co-precipitates with CD33^M^ and CD33^m^. These studies do not distinguish whether this interaction occurs in the intracellular compartment or whether they occur at the cell surface.

Dimerization of CD33 positions the ligand binding sites 40 Å apart, pointing outwards from the two-fold axis (Figure 2a). The spatially defined double binding site is likely to enhance substrate selection for larger moieties (see below), and to enhance binding of weak interaction partners via avidity effects.

To directly confirm the existence of CD33 dimers, we undertook four orthogonal biological and biophysical experiments.

First, Blue Native gel electrophoresis of total membrane lysates solubilised in 1% digitonin from the human THP-1 monocytic cell line, and from human blood monocytes, reveals that the endogenous CD33 protein migrates as a ∼240 kDa species (Supplementary Data Figure 2A). This closely approximates the combined mass (220 kDa) expected for two monomeric full-length glycosylated CD33 proteins (∼75 kDa each) together with the digitonin micelle (70 kDa). Due to the poor resolution of Blue Native Gels, we were not able to resolve size differences between CD33^M^/CD33^M^; CD33^M^/CD33^m^; and CD33^m^/CD33^m^ complexes. However, in agreement with prior results^29,30^, the abundance of the CD33^M^ dimers in monocyte lysates from homozygous minor allele carriers (AA genotype) was significantly lower than the abundance of CD33^M^ dimers in homozygous and heterozygous carriers of the common (risk) allele (CC and AC genotypes). This likely reflects the lower abundance of CD33^M^/CD33^M^ and CD33^M^/CD33^m^ dimers in minor allele carriers.

Second, reciprocal co-immuno-precipitation assays in transfected cells (Supplemental Figure 2B) confirm that combinations of differently tagged (-Myc or -HA) CD33^M^ and CD33^m^ co- precipitate with each other as CD33^M^/CD33^M^, CD33^M^/CD33^m^ and CD33^m^/CD33^m^ dimers.

Third, mutagenesis of selected key residues designed to disrupt the hydrophobic and polar interactions (Phe177Ala + Trp179Ala + Leu180Ala + Arg190Ala + Gln213Ala – CD33^M-5X^) form stable glycosylated CD33^M^ and CD33^m^ monomers that do not co-precipitate other CD33 peptides (Supplementary Data Figure 2B). The dimer was not disrupted by ≤4 mutations, reflecting the high affinity and complex structure of the dimer site. We cannot exclude that there may be some degree of misfolding of the CD33^M-5X^ protein, the presence of mature glycosylation of the CD33^M-5X^ mutant protein suggest that it is predominantly properly folded and processed in the ER and Golgi (Supplementary Data Figures 2B and 3C). The loss of dimerization is therefore unlikely to be solely due to misfolding of the mutant CD33^M-5X^ protein.

**FIGURE 3.**
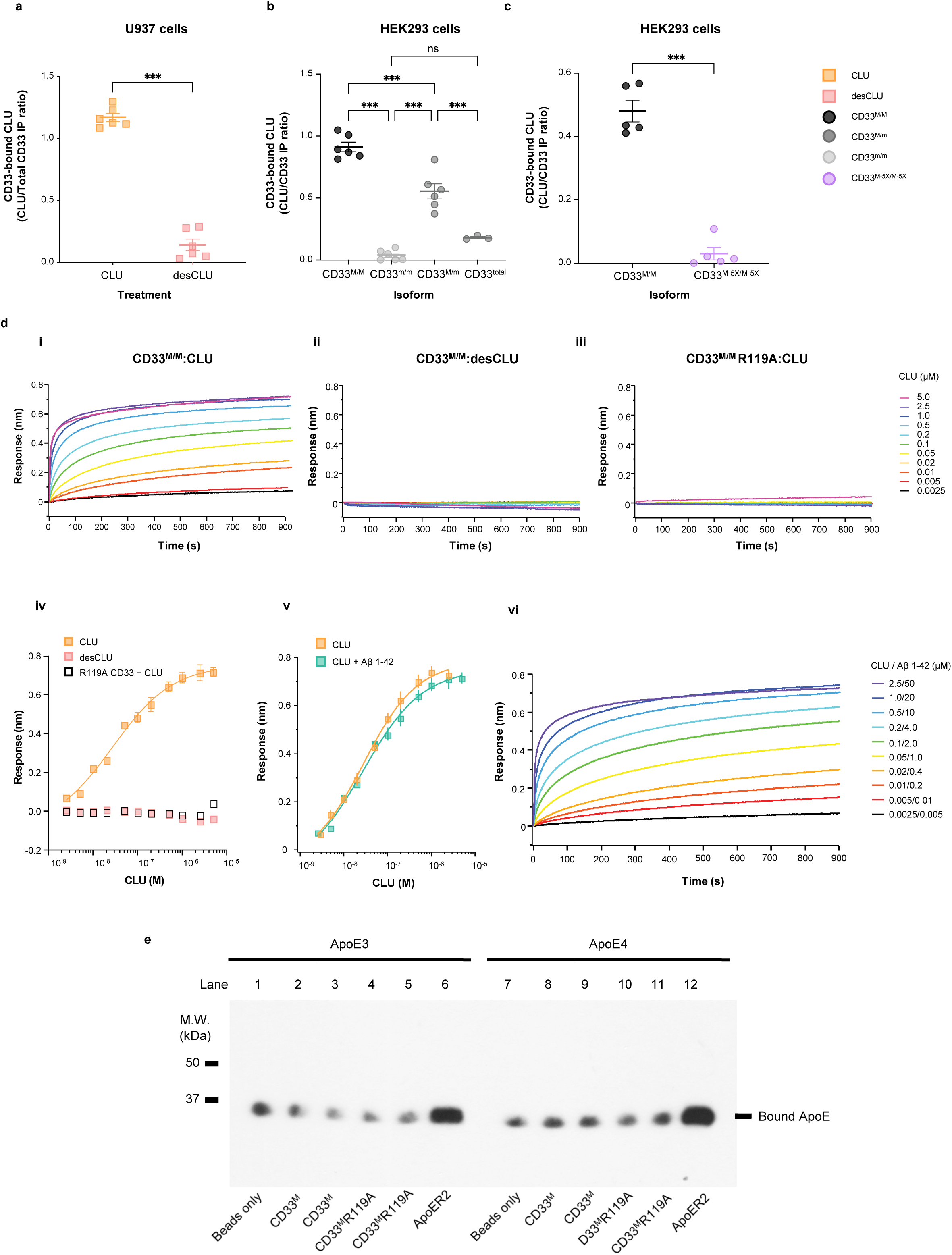
CLU and CD33 interact in vivo and in cells. a. **CD33^M^ co-immunoprecipitates sialylated CLU, but not desialylated CLU.** Endogenous human CD33 expressed on human monocytic U937 cells interact with native human plasma CLU, but not with desialylated CLU applied to the culture supernatant. The graph shows quantification of western blots of anti-CD33 co-immunoprecipitation products from cell lysates, blotted with anti-CLU antibodies. Quantification is performed by measuring the band intensities on autoradiograms of co-IP Western blots using anti-CD33 and anti-CLU antibodies to detect CD33 and coprecipitating CLU respectively. The result (CD33-bound CLU) is expressed as the ratio of CLU to CD33 band intensities as arbitrary units (a.u.). Representative western blot is shown in Supplementary Data Figure 3C left panel. *** = p<0.0001; Student t-test; error bars = mean ± SEM CLU: 1.23±0.04; desCLU: 0.23 ± 0.051; n = 5 independent replications. b. **CD33^M^ but not CD33^m^ co-immunoprecipitates with CLU.** The graph shows quantification of CLU co-precipitating with CD33^M^ in HEK293 cells as measured by the band intensities on western blots of the co-immunoprecipitating products. The results are expressed as CD33-bound CLU using arbitrary units to depict the ratio of CLU band intensity relative to the CD33 band intensity. HEK293 cells expressed: CD33^M^ only (CD33^M/M^ dimers), both CD33^M^ and CD33^m^ (CD33^M/M^; CD33^M*/*m^ or CD33^m/m^ dimers) (CLU does not co-precipitate with CD33 in cells expressing only CD33^m/m^ dimers). Graph columns 3 and 4 depict the results for cells expressing both CD33^M^ and CD33^m^. The third column shows the CLU: CD33^M^ ratio. The fourth column shows the CD33 bound-CLU ratio computed for total cellular CD33. The HA epitope tag on both CD33^M^ and CD33m is used to measure both CD33^M^ and CD33^m^ immunoreactivities (CD33^Total^). *A representative blot is shown in* Supplementary Data Figure 3C, right panel *** = p<0.00001; two-tailed Student t- tests; error bars = mean ± SEM CD33^M^: 0.85 ± 0.023; CD33m: 0.004 ± 0.00043; CD33^M^/^m^ : 0.45±0.041; n = 5 biological replications. c. **Dimerization of CD33 is required for CLU binding** Co-IP studies on lysates from HEK293 cells expressing similar quantities of CD33^M^ reveal that much less CLU is co- precipitated with the CD33^M-5X^ dimer interface mutant than with wild type CD33^M^. The data is quantified as in panel b. A representative western blot is available in Supplementary Data Figure 3D. *** p<0.00001, error bars = mean ± SEM CD33^M^: 0.425 ± 0.0073; CD33^M- 5X^: 0.012 ± 0.006; n = 5 replications. Each lane represents one independent biological replication of the respective condition. d. **CLU binds to CD33^M^ ECD with nanomolar avidity via sialic acids: i.** BLI curves of CLU binding to CD33^M^ ECD WT; **ii.** Desialylated-CLU showed no detectable binding to CD33^M^ ECD WT; **iii**. CD33^M^ ECD R119A mutant showed no detectable CLU binding. CLU concentrations are indicated. Each binding curve is the average of n = 4 independent measurements; **iv**. R119A mutation of the canonical sialic acid binding site disrupts CLU binding. Steady-state BLI analysis of the CLU binding to CD33^M^ ECD WT (yellow square) or CD33^M^ ECD R119A (open square) and desialylated-human CLU to CD33^M^ ECD WT (pink square). Curve fitting was performed using Hill equation. Equilibrium dissociation constant is 27.9 ± 7.2 nM. This result is not due to misfolding of the CD33^M^ R119A protein because the Far UV CD spectra of wild type and R119A proteins are similar (Supplementary Data Figure 3F). Error bars = SD. n = 4 independent replications; **v.** CLU+Aβ 1-42 showed similar binding avidity to CD33^M^ ectodomain with CLU alone. Steady-state BLI analysis of the CLU/Aβ 1-42 mixture binding to CD33^M^ ECD WT (teal square). yellow squares indicate binding responses of CLU alone to CD33^M^ ECD WT. Curve fitting was performed using Hill equation. Equilibrium dissociation constant of CLU/Aβ42 is 28.2 ± 4.9 nM; **vi.** BLI curves of CLU/Aβ 1-42 mixture binding to CD33^M^ ECD WT. Binding curves obtained from the same concentration of Aβ 1-42 alone was subtracted from the curve from CLU/Aβ 1-42 mixture. Each curve is the average of four independent measurements. CLU/Aβ 1-42 concentrations are indicated. Error bars = SD, n = 4 independent biological replications. e. **ApoE does not bind to CD33**. Representative western blots for co-immunoprecipitation of ApoE isoforms with His-tagged CD33 and ApoER2. 20µg wild-type (WT) and mutant (R119A) CD33 proteins with or without glycosylation (G) fused to His-tag were incubated with 50µl dynabeads, followed by incubation with 500µl of ApoE3 or ApoE4 (3µg/ml). ApoE bound to CD33 and ApoER2 was detected by immunoblots using anti-ApoE antibody. Beads only and ApoER2 ectodomain with His-tag were used for negative and positive controls, respectively. N = 4 replications.

Fourth, size exclusion chromatography-multi angle light scattering (SEC-MALS) analysis of the molar mass of both fully glycosylated and Endo H-trimmed samples of CD33 ECD in solution (Supplementary Data Figure 2C) revealed a mass of 50 kDa by UV (47 kDa, RI) for the trimmed sample at 10 µM, corresponding to an essentially fully dimeric form of the 25 kDa monomer, and a mass of 44 kDa by UV (41 kDa, RI) at 2 µM concentration. In parallel analyses, the glycosylated CD33 ECD display a mass of 43 kDa for the protein component, and 18 kDa for the carbohydrate modifier at 10 µM (Supplementary Data Figure 2C). This result is in good agreement with the mass expected for GlcNAc_2_Man_9_ glycans synthesized in the presence of kifunensine. Based on these analyses, the Kd of the ECD dimer is approximately 10^-7^ M, and indicates that this interaction will occur at physiologic concentrations, and is unlikely to be a biologically irrelevant crystal artefact.

These dimerization affinities were confirmed by analytical ultra-centrifugation sedimentation velocity (AUC-SV) and sedimentation equilibrium (AUC-SE) experiments (Kd of 7.1 ± 0.01 x 10^-7^ M for de- glycosylated CD33, and 3.0 ± 0.1 x 10^-7^ M for glycosylated CD33 (n = 3 measurements; Supplementary Data Figures 2D-I).

Taken together, these experiments suggest that the multimerization (and specifically dimerization) of CD33 is likely to occur physiologically in cells.

### CD33^M^ is targeted to the cell surface

Recent work has suggested that CD33^m^ is predominantly targeted to peroxisomes, and that little if any reaches the cell surface^39^. However, the dimerization of CD33 raises questions about the cell surface disposition of CD33^M^ and CD33^m^ species in cells expressing both CD33^M^ and CD33^m^.

To investigate the trafficking of CD33 isoforms to the cell surface, we developed HEK293 cells expressing equivalent total cellular quantities of CD33 as either: CD33^M^ only (CD33^M/M^); CD33^m^ only (CD33^m/m^); or both CD33^M^ and CD33^m^ (CD33^M/m^). The cellular quantity of CD33 was measured by western blots of cell lysates using anti-HA antibody to HA-tags on both CD33^M^ and CD33^m^. Prior work has shown that exogenous CD33^M^ and CD33^m^ are correctly expressed in HEK293 cells^40^. We then used antibodies that recognised CD33^M^ only (WM53); CD33^m^ only (A1612H); or both CD33^M^ and CD33^m^ (HIM3-4) coupled with flow cytometry to assess trafficking of the CD33 isoforms to the surface of these cells.

Our flow cytometry assays investigated 1×10^5^ cells for each genotype and were conducted as 6 replicates in 3 independent experiments. The majority (81.2%±9.3%) of CD33^M/M^ cells express CD33^M^ at the cell surface, whereas only 36.1%±2.9% of CD33^m/m^ cells had detectable cell surface CD33^m^ (Figure 2c). Furthermore, the intensity of surface exposure of CD33^m^ in CD33^m/m^ cells was much lower than the corresponding intensity for CD33^M^ in CD33^M/M^ cells (as measured by the HIM3-4 antibody which recognises the same extracellular HA-epitope in CD33^m^ and CD33^M^) (Figure 2d, Supplementary Data Figure 2J). Intriguingly however, in CD33^M/m^ cells, despite expressing similar overall levels of CD33^M^ and CD33^m^, most CD33^M/m^ cells (83.2%±6.5%) express CD33^M^ at the cell surface. Only a small proportion of CD33^M/m^ cells (17.7%±1.8%) express both CD33^M^ and CD33^m^ at the cell surface (Figure 2c, Supplementary Data Figure 2J).

### Imaging CD33 receptor dimers in mammalian cells by smFRET

To determine whether the surface targeted receptors dimerize in the plasma membrane, we used a recently developed single-molecule fluorescence resonance energy transfer (smFRET) method for imaging transmembrane receptor complexes in a cellular context^41^. This approach has the distinct advantage of offering the molecular-scale resolution needed to directly measure full-length CD33 receptor dimers diffusing within the plasma membrane of living mammalian cells. We prepared Chinese Hamster Ovary (CHO) cell lines stably expressing amino-terminally SNAPfast (SNAP_f_)-tagged CD33^M^, CD33^m^, or CD33^M-5x^ receptors under a tetracycline regulatable promoter. This produces low basal, tetracycline-regulatable receptor densities that are compatible with imaging and tracking single molecules in the plasma membrane. We used the membrane- impermeant, SNAP_f_-tag reactive Lumidyne 555p (LD555p) and 655 (LD655) dyes as the smFRET donor and acceptor fluorophores, respectively. This strategy allows stochastic labelling of the SNAP_f_-CD33 receptors (Figure 2e).

Using total internal reflection fluorescence microscopy (TIRF) for imaging, the CHO cells were visualized briefly by direct and simultaneous donor and acceptor excitation immediately before smFRET imaging, which enables quantification of the surface density of labelled receptors. Note that the amount of tetracycline used to induce expression of each cell line was empirically determined such that the surface expression of the examined CD33 receptors were similar. The median surface density of total labelled receptors as determined by single-particle detection ranged from 0.34 – 0.60 molecules/μm^2^ (Supplementary Data Figure 2K), consistent with those in our previous smFRET study of G protein-coupled receptor (GPCR) interactions^41^.

Homodimerization between CD33 protomers was assessed by measuring the total number of freely diffusing smFRET events per cell area. Because smFRET from immobilized or confined receptors could result from close packing of receptors within membrane microdomains and not from their direct interaction at a dimeric interface, we focused on smFRET trajectories from CD33 receptors freely diffusing within the membrane as previously described (Figure 2f-h; Supplementary Data Figure 2K)^41^. Using this approach, CD33^M^ labelled with both donor and acceptor showed numerous smFRET events (Figure 2f-h, Supplementary Data Figure 2K). However, as expected, the same receptor labelled with only donor (which serves as a negative control and estimates donor bleed through into the acceptor channel) showed little to no smFRET events (Figure 2 f-h, Supplementary Data Figure 2K). These results strongly suggest that full-length CD33^M^ protomers efficiently interact in the plasma membrane, consistent with the dimeric CD33^M^ extracellular domain crystal structure. Conversely, CD33^M-5X^ showed little to no smFRET events, and presented almost identically in distribution to that of the donor only negative control despite being expressed at a similar surface density (Figure 2h, Supplementary Data Figure 2K). This result is consistent with the 5X mutations disrupting dimerization within CD33^M^, supporting the notion that this region of the

extracellular domain serves as a major interface stabilizing the dimer. However, as discussed below, although not seen in the crystal structure, we do not exclude that other motifs within the N-terminal domain may also contribute to dimerization *in vivo*. Interestingly, CD33^m^ showed significantly less smFRET events compared to CD33^M^ even when expressed at comparable levels but showed more smFRET events than the donor-only control (Figure 2g,h, Supplementary Data Figure 2K). The lower number of mobile smFRET events for CD33^m^ relative to CD33^M^ was not due to differences in time spent in other diffusion states (Supplementary Data Figure 2K). This result suggests that CD33^m^ is less prone to dimerization at the cell surface, and raises questions as to whether other motifs within the N-terminal domain that are not shared by CD33^M^ and CD33^m^ may also contribute to dimerization *in vivo*.

### A CD33 frameshift mutant does not form dimers and does not alter AD-risk

A rare four base pair deletion (rs201074739) in exon 3 of CD33 has recently been described^42^. This variant, which has been referred to as a “null allele”, is predicted to cause a frameshift with production of a C-terminally truncated protein comprising residues 1-155 of CD33^M^ plus four additional residues followed by a stop codon. The encoded protein is predicted to contain the signal peptide and the ligand binding site within the V-set domain of CD33^M^. However, it contains only 12 residues within the linker and C2-set domains. It does not include the dimer site, transmembrane domain or ITIM/ITIM-like domains (Figure 1a). As a result, if this truncated protein (CD33^MΔ4bp^) is biochemically stable, it might be secreted and could potentially have biological properties. However, because this allele is rare (0.02) the majority of individuals will be heterozygous for this allele and for the major “C” allele at rs3865444, which promotes production of full-length CD33^M^. Genetic studies of the truncated CD33^MΔ4bp^ variant in a small dataset (n = 21,982 AD cases; 41,944 non-AD cases) showed that it is not associated with reduced risk (p = 0.133; OR = 0.90 with 95% CI 0.79-1.03). This contrasts strongly with the minor “A” at rs3865444 which, in the same study^43^, is associated with increased expression of the CD33^m^ isoform and with reduced risk of AD.

To explore the biological properties of CD33^MΔ4bp^, we transfected HEK293 cells with CD33^MΔ4bp^ alone or with CD33^M^ or CD33^m^ cDNAs (using the same methods as described above). We find that low levels of CD33^MΔ4bp^ protein are present in lysates of cells expressing either CD33^MΔ4bp^ alone or CD33^M^ plus CD33^MΔ4bp^ (Figure 2i, left panel). Intriguingly, we also observed significant amounts of the CD33^MΔ4bp^ protein in the media from cells expressing CD33^MΔ4bp^ alone or CD33^M^ plus CD33^MΔ4bp^ (Figure 2i, right panel) at 48 hours incubation. However, unexpectedly, we also observe a truncated CD33^M^ protein in the media of cells expressing CD33^M^ only (Figure 2i, right panel). This soluble truncated CD33^M^ protein (hereafter sCD33^M^) contains epitopes recognised with antibody clone EPR4423 (to residues 30-80aa) in the N-terminus of CD33^44M^ (Figure 2i, right panel). Crucially, this CD33^M^ ectodomain-containing species is not present in the media of cells expressing CD33^MΔ4bp^ only (Figure 2i, right panel). We interpret this sCD33^M^ to represent a previously unrecognised endoproteolytic cleavage product (possibly similar in origin to those observed for other AD-related Type I membrane proteins such as sAPP and sTREM2)^45–49^. Because this protein is seen in cells expressing the exogenous full-length CD33^M^ cDNA, it is unlikely to reflect a splice variant lacking the transmembrane and ITIM domains. The CD33^M^ soluble fragment (sCD33^M^) is also present as a fuzzy (likely glycosylated) ∼38 kDa species in western blot studies of conditioned medium from cultured human iPSC derived macrophages and microglia (Figure 2j). In an orthogonal set of experiments, we used an anti-CD33 ELISA assay directed at the N-terminus of CD33^M^ (Abcam, ab283542) to confirm the presence of sCD33^M^ species in the media of iPSC cells and the sensitivity of sCD33^M^ species to 5µM batimastat (a potent broad spectrum matrix metalloprotease inhibitor, Sigma-Aldrich, SML0041) (Supplementary Data Figure 2L). Additional work will be required to map the precise C-terminal residues, confirm that they do not conform to an exon splice boundary, and identify the putative sheddases.

Because the CD33^MΔ4bp^ protein contains an intact ligand binding site, we were curious as to whether it could potentially interact with CD33^M^ or CD33^m^ proteins. Our initial expectation was that this would be unlikely because CD33^MΔ4bp^ lacks residues within the C2-set domain that form the dimer interface. We transfected HEK293 cells with HA-tagged CD33^M^ together with CD33^MΔ4bp^, or with HA-tagged CD33^m^ together with CD33^MΔ4bp^ (Figure 2k) To our surprise, CD33^MΔ4bp^ is readily detectable in the HA-immunoprecipitation products from lysates of cells co-expressing CD33^MΔ4bp^ with CD33^M^-HA (Figure 2k). CD33^MΔ4bp^ protein is also present in the HA-immunoprecipitation products from lysates of cells co-expressing CD33^MΔ4bp^ with CD33^m^-HA (Figure 2k). A cartoon depicting these putative interactions is available in Supplementary Data Figure 2M. These experiments raise the possibility that the reason why CD33^MΔ4bp^ is not associated with a reduction in AD risk is that it bears a ligand-binding site that may confer biological properties that replicate some aspects of the deleterious properties of CD33^M^. For instance, CD33^MΔ4bp^ might form duplex chimeric CD33^M/MΔ4bp^ receptors with an intact ligand binding site. Alternatively, CD33^MΔ4bp^ might bind to other cell surface sialylated molecules, potentially altering their signaling activities. If so, sCD33^M^ would likely share the same biological activity. In this context, it is noteworthy that sCD33^M^ is exclusively a product of CD33^M^ but not CD33^m^ – adding an additional difference in the cell biology of these two splice forms of CD33 that may play into how variants that upregulate CD33^M^ expression could increase risk for AD.

### CD33^M^ binds the AD-relevant ligand CLU but not ApoE

CD33 binds to a variety of sialylated moieties (especially di-sulphated glycans^50–52^). Well-documented ligands include hepatitis B surface antigen and a glycosylated isoform of Receptor Protein Tyrosine Kinases ζ (RPTPζ^S53L^)^53^. However, the presence of two ligand binding sites in slightly different orientations, raises the possibility that CD33 might bind larger soluble and/or cell-surface sialylated ligands. In an accompanying manuscript (Vo et al.), we show that CD33 interacts with several large sialylated cell surface molecules, including CD45. However, we were curious as to whether CD33 might also interact with large soluble sialylated ligands. Two such proteins, clusterin (CLU ∼67 kDa^54^) and apolipoprotein E (ApoE) stood out because of their known association with AD pathobiology^10,28,29,55–62^.

To determine whether CLU and/or ApoE might be ligands for CD33, we grew human U937 cells (endogenously expressing CD33) and HEK293 cells expressing exogenous CD33 in media containing physiological levels of CLU (60 nM) or ApoE (3 µg/ml). We then undertook anti-CD33 immunoprecipitation experiments and searched for these ligands in the immunoprecipitates. These co-immunoprecipitation experiments reveal that both sialylated native CLU (from human plasma) and sialylated recombinant human CLU (characterised in Supplementary Data Figures 3A) co-precipitate with endogenous CD33^M^ in U937 cells (Figure 3a, Supplementary Data Figure 3B, left panel). Sialylated human CLU also co-precipitates with CD33^M^ in HEK293 cells expressing either CD33^M^ alone (CD33^M^/CD33^M^) or CD33^M^ plus CD33^m^ (CD33^M^/CD33^m^) (Figure 3b, Supplementary Data Figure 3B right panel). However, CLU does not co-precipitate with CD33 in HEK293 cells expressing either CD33^m^ alone or the dimer-disruptive CD33^M-5X^ mutant (Figure 3b,c, Supplementary Data Figure 3C). Desialylated CLU does not co-immunoprecipitate with endogenous CD33 expressed in human U937 cells (Figure 3a, Supplementary Data Figure 3B left panel) and does not bind to CD33 lacking the canonical R119 ligand binding site residue (Supplementary Data Figure 3D,E). These results indicate that CLU: CD33 binding requires sialylation of CLU, an intact CD33 ligand binding site (absent in CD33^m^) and a dimeric configuration of CD33M.

To quantitatively assess the CD33^M^:CLU interaction, we used bio-layer interferometry (BLI), immobilizing CD33^M^ ECD (0.5 µM) and then exposing it to various concentrations of CLU (2.5 nM- 5.0 µM) (Figure 3d, panel i, Supplementary Data Figures 3E,F). Because CLU forms oligomers in solution, we employed steady-state BLI analysis to calculate a dissociation constant. These quantitative experiments show that CD33^M^ ECD strongly binds CLU with Kd values that are likely to be physiologically relevant (Kd = 28.9 ± 10.3 nM; CLU concentration in CSF = 60 nM^63^). These results were independently confirmed by microscale thermophoresis and by measuring the change in the fluorescence intensity of fluorescently labelled CD33 ECD in response to CLU binding (Kd for CLU binding 55.1 ± 2.6 nM; Supplementary Data Figure 3G). To confirm that the interactions assessed by BLI were dependent upon engagement of sialic acid moieties on the ligand with the cognate Arg119 in the ligand binding site, we also assessed the binding avidity for desialylated CLU, and for CD33 lacking the cognate Arg119 docking residue. There was no binding between CD33^M^ and desialylated CLU (Figure 3d, panel ii). There is also no binding between CLU and mutant CD33^M^ R119A ECD (Figure 3d, panel iii, iv). The absence of CLU binding with CD33 ^M^ R119A is not due to simple misfolding of the CD33^M^ ECD R119A protein because the far-UV circular dichroism spectra of the wild-type and the R119A mutant CD33 ECD proteins are highly similar. Thus, both wild-type and R119A spectra display intense minima at ∼215 nm, typical of a β- sheet-rich protein (Supplementary Data Figure 3H).

We could not undertake similar studies on recombinant ectodomains of CD33^m^ and CD33^M-5X^ because, without their stabilizing TM and intracellular domains, they tend to aggregate during purification, making them unsuitable for BLI. Nevertheless, the results of the cell biological studies described in Figure 3a-c are not due to aggregation of the full length CD33^m^ and CD33^M-5X^ proteins. In cells, both CD33^m^ and CD33^M-5X^ proteins are well expressed (including *in vivo* for CD33^m^) and they are glycosylated – a feature that does not usually occur on misfolded proteins.

We repeated these co-immunoprecipitation studies using physiological concentrations of fully sialylated ApoE secreted from HEK293 cells^64,65^. To our surprise, we found that, under the experimental conditions that we used, neither ApoE ε3 nor ApoE ε4 show significant binding to CD33^M^ and give only background signals similar to that of beads, even when co-precipitated using a case-affinity pulldown approach with high concentrations of His-tagged CD33 CD33^M^ (Figure 3e). In contrast, both ApoE ε3 and ApoE ε4 readily bind the ApoER2 receptor^66^ under similar conditions (Figure 3e). However, ApoE undergoes highly variable and complex O-glycosylation patterns in different cell types and cell activation states^64,67,68^. Our conclusions are therefore naturally limited to our experimental system where we did not detect APOE binding to CD33^M^.

Taken together, these experiments indicate that CLU is a *bone fide* ligand for CD33, while ApoE is probably not. CD33^M^:CLU binding requires both engagement of the sialic acid binding site on CD33 via the cognate Arg119 docking site and dimerization of CD33^M^. The apparent lack of a significant in vitro interaction of ApoE with CD33 underscores the specificity of CLU binding to CD33. It suggests that other properties of CD33 and/or CLU, such as steric requirements (e.g. contributed by the positioning of the CD33 CC’ loop, as noted above) or another epitope are required for their high avidity interaction.

### CD33 interacts with CLU + Aβ complexes in AD brain

Next, to determine whether the CLU:CD33^M^ interaction occurs *in vivo*, and is relevant to AD, we applied two independent assays.

First, we used an *in-situ* proximity ligation assay (PLA) to show that endogenous CLU colocalized with CD33 on Iba1(+)ve microglia in post-mortem human brain with Braak Stage 6 Alzheimer Disease (Figure 4a). Crucially, the density of CD33:CLU colocalization in each microglial cell is a function of the proximity of that specific microglial cell to amyloid plaques (as identified by 6E10 anti-Aβ antibody staining) (Figure 4b). Amyloid plaques are a source of both CLU and Aβ oligomers, and CLU levels are higher in AD brain and in areas rich in Aβ pathology^69,70^. No PLA signal was observed in controls where one of the antibodies was replaced with an isotype control.

**FIGURE 4.**
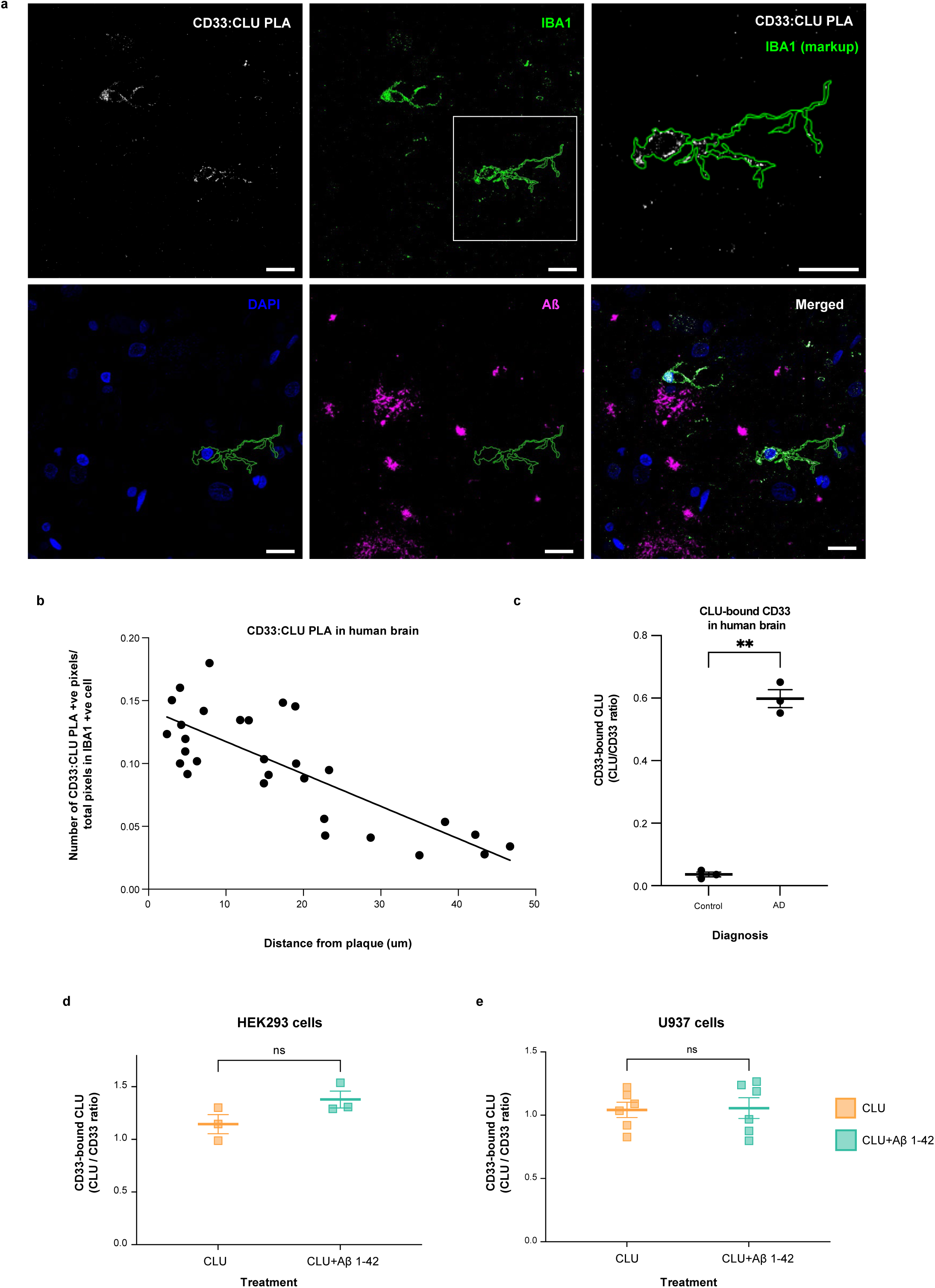
CD33 interacts with CLU and with CLU plus Aβ oligomers in human AD-affected brain, and cultured human cells. a. ***In situ* Proximity ligation assay for CD33-bound CLU on IBA-1 positive microglial cells in post-mortem human AD brain**. Representative confocal microscopy images showing CD33:CLU PLA signals in microglia close to amyloid plaques in human AD brain cortex (63x, scale bar = 50 µm). An enlarged view of the PLA signal is shown in the upper right panel. FFPE sections were stained for 6E10 Aβ = violet; IBA1 microglia = green; PLA = red; and DAPI nucleus = blue. b. **Dot plot showing *In vivo* correlations between microglial CD33:CLU PLA signal intensity and the distance of the microglial cell from Aβ plaque.** The proportion of CD33-CLU positive pixels for each IBA-1 positive microglial cell was plotted relative to the distance of that cell from the edge of Aβ plaque (r^2^ = 0.6235). There was a negative correlation between distance from the plaque edge and the proportion of CD33-CLU PLA positive pixels in microglia in human AD brain (spearman r = -0.739, *p* < 0.0001, n = 29 microglia from n = 2 independent AD affected brains). c. **More endogenous CLU can be co-immunoprecipitated with endogenous CD33 from human AD brain than from control brain.** The graph shows quantification of CLU:CD33 co-immunoprecipitation (CD33-bound CLU) in brain lysates from AD cases (n = 3), but only weak CLU:CD33 co-immunoprecipitation is seen in brain lysates from control cases (n = 3). Representative co-IP western blot of lysates and anti-CD33 co-immunoprecipitation products probed with anti-CLU antibodies is observed in Supplementary Data Figure 4A. *** = p < 0.001; ANOVA; error bars = mean ± SEM; n = 3 independent samples. Because both CLU and CD33 levels are highly variable in post-mortem tissue, brains were selected based on equal expression levels of both CLU and CD33 from a panel of n=53 control and AD cerebral cortices. d. **Native human CLU (either alone or complexed with Aβ1-42 oligomers) also binds exogenous human CD33^M^ on transfected HEK293 cells.** The graph shows quantification of CLU:CD33 co-IP products (CD33-bound CLU). A representative co-IP western blot is shown in Supplementary Data Figure 4B, right panel. There is no significant difference in the amount of CLU bound to CD33M in lysates from HEK293 cells expressing exogenous CD33M and exposed to CLU alone versus CLU plus Aβ oligomers. NS = not significant; Student t-test; error bars = mean ± SEM CLU: 1.145± 0.0904; CLU+Aβ1-42: 1.379± 0.0798; n = 3. Each dot represents one independent biological replication of the respective condition e. **Native human CLU (either alone or complexed with Aβ1-42 oligomers) also binds endogenous human CD33^M^ physiologically expressed on U937 cells**. Representative co-IP western blot is shown in Supplementary Data Figure 4B, left panel. There is no significant difference in the amount of CD33-bound CLU in lysates from U937 cells exposed to CLU alone versus CLU plus Aβ oligomers NS = not significant; Student t-test; error bars = mean ± SEM; CLU: 0.928± 0.06; CLU+Aβ1-42: 0.881± 0.0491; n = 6. Each dot represents one independent biological replication of the respective condition.

An interaction between CLU and CD33 *in vivo* is further supported by orthogonal co-immunoprecipitation experiments from human brain tissue. Anti-CD33 co-immunoprecipitation assays of lysates from AD cerebral cortex (n = 3) and non-AD control cortex (n = 3), revealed direct physical interaction between endogenous CLU and endogenous CD33 in AD brain tissue, and to a lesser extent, in control brain (quantified in Figure 4c, representative co-immunoprecipitation western blot image in Supplementary Data Figure 4A).

The increased CLU:CD33 co-immunoprecipitation observed in AD brain lysates likely results simply from increased abundance of both CLU and of activated microglia expressing CD33 in AD-affected cortex. However, another possible explanation is that CLU:Aβ oligomer complexes, which are enriched in AD brain^71,72^, might have a higher binding avidity than CLU alone. To examine this possibility, we used the same BLI method described above to estimate the avidities of CD33^M^ for Aβ oligomers alone, CLU alone, and CLU + Aβ oligomers. CD33^M^ only weakly binds Aβ oligomers (Supplementary Data Figure 3E,F). However, CD33^M^ binds with high, but similar avidities to both CLU alone and CLU + Aβ oligomers (Kd = 28.2 ± 4.9 nM), especially at physiological CLU concentrations around 100 nM (Figure 3d, panels v-vi; Supplementary Data Figures 3E,F). This result is strongly supported by co-immunoprecipitation studies from HEK293 cells (expressing exogenous CD33^M^) and human U937 monocyte-like cells (expressing endogenous CD33^M^), which also showed no significant difference in CD33^M^ binding to CLU alone and CLU + Aβ oligomers (quantified in Figure 4d,e).

Together, these experiments support the notion that under conditions that would be operative in AD-affected brain tissue, CD33^M^ forms high avidity interactions with both CLU alone and CLU + Aβ oligomers, but not with ApoE. Crucially, this interaction is dependent upon both an intact sialic acid binding site and the formation of CD33^M/M^ dimers.

### CLU induces CD33^M^ ITIM signalling

These experimental results raise the question as to whether binding of CLU ± Aβ oligomers to CD33^M/M^ can modulate CD33^M^ ITIM signalling.

Previous work has used sodium orthovanadate to induce CD33 ITIM phosphorylation or binding of inactivating antibody to investigate CD33 signalling. These studies have implied, but not robustly proven, that ligand binding to the CD33^M^ extracellular domain activates CD33 ITIM signalling via: i) phosphorylation of one or more tyrosine residues in the intracellular ITIM domain (e.g. Tyr340) ^73–75^, and ii) recruitment of the Src-homology 2 domain (SH2)-containing protein tyrosine phosphatases 1 and 2 (SHP-1 and SHP-2)^73,74^. Based upon the idea that binding of ligands like CLU might induce CD33^M/M^ ITIM signalling, we next applied physiological concentrations (60 nM) of CLU ± Aβ oligomers to human monocyte-like U973 cells. We then assessed their impact on phosphorylation of endogenous CD33 and recruitment of endogenous SHP-1 by western blotting.

These experiments reveal the induction of CD33^M/M^ tyrosine phosphorylation and SHP-1 recruitment within 5 - 15 minutes after application of recombinant or human plasma-derived CLU to U937 cells (Figure 5a(i) and 5a(ii), p<0.05 – 0.001; representative western blots in Supplementary Data Figure 5A). We confirmed and extended these conclusions in HEK293 cells expressing CD33^M^, showing that recombinant and human plasma CLU ± Aβ 1-42 equivalently augment CD33 ITIM phosphorylation, while desialylated CLU did not (Figure 5b, p<0.05 – 0.001; representative western blots in Supplementary Data Figure 5B).

**FIGURE 5.**
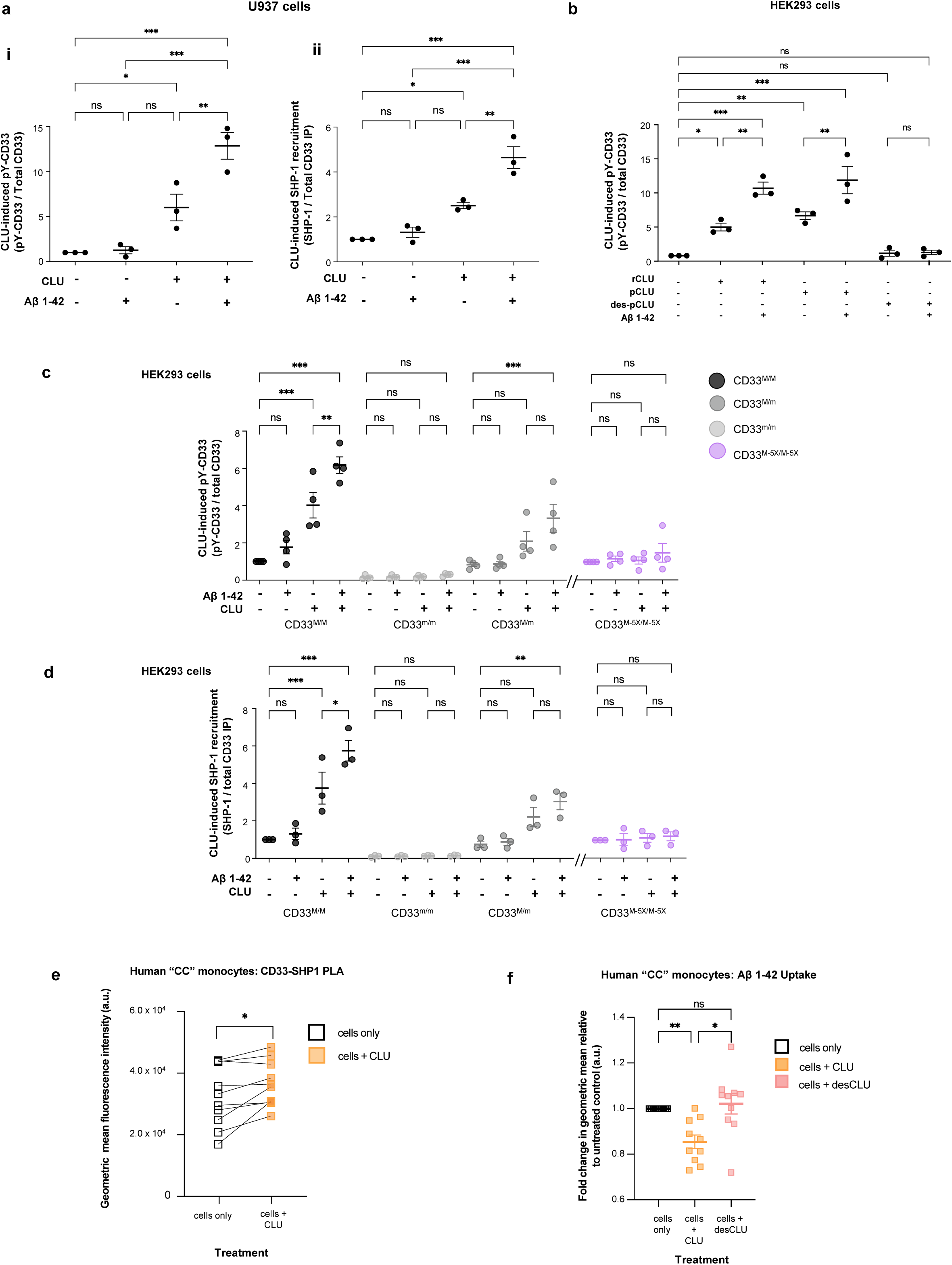
CLU, and especially CLU + Aβ oligomers, induce CD33 ITIM tyrosine phosphorylation of CD33^M^/CD33^M^ and CD33^M^/CD33^m^ dimers, but not CD33^m^/CD33^m^ dimers. a. **Human plasma CLU alone, and especially human CLU + Aβ oligomers induce ITIM tyrosine phosphorylation of endogenous CD33 *(left panel)* and recruitment of endogenous SHP-1** *(right panel)* in human monocytic U937 cells. Quantification of CD33 phosphotyrosine (pY20 antibody immunoreactive anti-CD33 IP products) and of SHP-1 recruitment (anti-SHP-1 immunoreactive anti-CD33 IP products) using band intensity (representative blots are in Supplementary Data Figure 5A. * = p <0.05; *** = p <0.001; NS = not significant; ANOVA with Tukey post-hoc correction; error bars = mean ± SEM; For CD33^M^-pTyr, No treatment: 1.00±0.0; Aβ treatment: 1.367±0.4107; CLU treatment: 6.283±1.667; CLU + Aβ treatment: 11.92±1.116; For SHP-1, No treatment: 1.00±0.0; Aβ treatment: 1.447±0.2726; CLU treatment: 2.589±0.0688; CLU + Aβ treatment: 4.362±0.5971, n ≥ 3 independent biological replications. b. **Recombinant human clusterin (rCLU) and native human plasma clusterin (pCLU), but not desialylated clusterin (desCLU) equivalently activate CD33 ITIM phosphorylation and SHP-1 recruitment in HEK293 cells**. CD33 tyrosine phosphorylation and SHP-1 recruitment were measured by western blots band intensity ratios for pY20-phosphotyrosine immunoreactive CD33 versus total CD33 by co-IP western blots. with different cell lines are available in Supplementary Data Figure 5C. c. **CLU + Aβ induces greater CD33 ITIM phosphorylation than CLU alone, but only in CD33M expressing cells.** Quantification of western blots of anti-CD33 immunoprecipitates probed with pY20 anti-phosphotyrosine or anti-HA antibodies reveals that treatment with CLU alone, and especially treatment with CLU + Aβ oligomers, increases CD33 tyrosine phosphorylation in cells expressing either CD33M only or CD33^M^ plus CD33^m^. No CD33 phosphorylation was observed in cells expressing only CD33^m^, and very little in CD33^M-5X^ cells. n = 3 replications. Representative western blots are in Supplementary Data Figure 5D. d. **CLU + Aβ induces greater CD33 SHP-1 recruitment than CLU alone, but only in CD33M expressing cells.** Quantification of western blots of anti-CD33 immunoprecipitates probed with SHP-1 or anti-HA antibodies reveals that treatment with CLU alone, and especially treatment with CLU + Aβ oligomers, increases SHP-1 recruitment in cells expressing either CD33M only or CD33^M^ plus CD33^m^. No SHP-1 recruitment was observed in cells expressing only CD33^m^, and very little in CD33^M-5X^ cells. n = 3 replications. Representative western blots are in Supplementary Data Figure 5D. e. **Treatment of human monocytes with 0.06 µM CLU for 15 min at 37°C increases binding of CD33 and SHP-1.** Human monocytes (genotype CD33 rs3865444^CC^) were treated with CLU and then a proximity ligation assay (PLA) was performed to examine CD33-SHP-1 interaction. Median fluorescence intensity of the PLA of CD33-SHP-1 was measured and analyzed in relation to untreated monocytes from the same individuals. Data were analyzed using paired t-test on GraphPad Prism. *, p<0.05. f. **CD33 rs3865444^CC^ risk genotype-dependent effect of CLUon monomeric Aβ1-42 uptake by human monocytes.** Human monocytes (genotype CD33 rs3865444^CC^) were treated with either CLU or desCLU, and the effect on Aβ1-42 uptake was assessed by flow cytometry. CLU significantly reduces the Aβ 1-42 uptake compared to the media control or desCLU treatment. Data were analyzed using geometric mean intensity of HiLyte™ Fluor 647 conjugated Aβ 1-42, n = 10 unrelated donors each examined once. Each dot represents an individual donor. Data were analyzed using ANOVA on GraphPad Prism. **, p<0.01; *, p<0.05.

To explore the role of differing surface abundance of CD33^M^ and CD33^m^, as well as the role of CD33 dimerization, we repeated these experiments in the HEK293 cell lines described in Figures 3 and 4. Based upon Western blotting studies, these cells express similar overall quantities of CD33 as either CD33^M^ only (CD33^M^/CD33^M^); CD33^M^ and CD33^m^ (CD33^M^/CD33^m^); CD33^m^ only (CD33^m^/CD33^m^); or CD33^M-5X^. As expected from the reduced effciency of CD33^m^ or CD33^M-5X^ dimerization at the cell surface (see Figure 2), the HEK293 cell lines expressing these proteins show little or no CD33 ITIM phosphorylation or SHP-1 recruitment (Figure 5c,d; Supplementary Data Figure 5C,D). In contrast, cells expressing CD33^M^ (i.e. CD33^M^/CD33^M^ cells and CD33^M^/CD33^m^ cells) robustly phosphorylate CD33 and recruit SHP-1 (Figure 5c-e; Supplementary Data Figure 5C,D). Moreover, in CD33^M^/CD33^m^ cells, only the CD33^M^ isoform is tyrosine phosphorylated (Figure 5c,d, but the overall efficiencies of both the CD33^M^ phosphorylation and the recruitment of SHP-1 is similar in CD33^M^/CD33^M^ and CD33^M^/CD33^m^ cells (Supplementary Data Figure 5C,D). To explain this observation, we suggest that CD33 ITIM signalling is likely to be regulated by two features.

First, in cells expressing both CD33^M^ and CD33^m^, CD33^m^ is much less prone to dimerize and to participaite in ITM signalling even if it gets to the cell surface. Second, in cells expressing both CD33^M^ and CD33^m^, sufficient numbers of functional CD33^M/M^ dimers are formed on the surface to support ITIM signalling in a manner similar to cells expressing only CD33^M^. The molecular mechanism in the secretory pathway that regulates the formation and abundance of CD33^M^ homodimers on the cell surface represents a fascinating future question for the field.

We replicated these results in human blood derived monocytes and in monocyte-derived microglial like cells (MDMi) expressing endogenous CD33. We use flow cytometry proximity ligation assays (flowPLA)^76^ to show that 60 nM CLU induces rapid recruitment (15 minutes) of endogenous monocyte SHP-1 to CD33^M^ (CD33-SHP1 PLA)(Figure 5e, Supplementary Figures 5E, p <0.05). In parallel, we use western blotting to show that CLU induced tyrosine phosphorylation of the endogenous monocyte CD33 ITIM domain (Supplementary Figures 5F). We show that treatment of human monocytes with 60 nM CLU suppressed Aβ_1-42_ monomer uptake, in contrast, treatment of human monocytes with desialylated CLU resulted in Aβ_1-42_ monomer uptake is unchanged compared to mock-treated monocytes (Figure 5f). Finally, we show that 15-minute treatment with 60 nM CLU supresses Aβ_1-42_ monomer uptake in human MDMi cells (Supplementary Figure 5G; p< 0.05, Wilcoxon Test).

Upon further examining these results, we noticed that CLU alone and CLU + Aβ oligomers have similar CD33^M^-binding avidities. However, despite this, CLU + Aβ oligomers consistently induce larger increases in CD33 ITIM signalling (Figure 5a, c, e; p<0.05 – 0.001). The structural mechanism by which CLU + Aβ oligomers lead to stronger CD33^M/M^ signalling agonism compared to CLU alone is unclear. The similar binding affinities for CLU and CLU + Aβ oligomers indicate that the different magnitudes of their agonist effects on CD33 signalling are not simply a consequence of extra “stickiness” imparted by Aβ oligomers. A more plausible explanation is that Aβ oligomers complexed with CLU could alter the shape of the ligand’s fit into the CD33^M/M^ binding pocket. This might result in more robust allosteric effects on the intracellular ITIM domain with enhanced CD33 signalling. Future structural studies will be needed to resolve this intriguing question.

Taken together, our experiments confirm that the binding of CLU (± Aβ oligomers) activates CD33 ITIM signalling, and this activation requires both the presence of an active sialic acid binding site and dimerization of CD33^M^.

### CLU but not desialylated CLU modulates CD33-dependent Aβ uptake in human monocytes

To explore the functional consequences of CLU-induced CD33 signalling (as measured by CD33 ITIM phosphorylation), we next exploited a previously described *ex vivo* assay which measures monomeric Aβ uptake in human monocytes^30^. This assay also capitalizes on the fact that CLU preferentially binds Aβ oligomers, not monomers^71,77^, thereby allowing us to deconvolute uptake of monomeric Aβ as phagocytic cargo from the effects of Aβ oligomers on CLU-mediated CD33^M^ signalling. We have previously shown that uptake of Aβ monomers by human monocytes is greatly reduced in monocytes from “CC” risk allele carriers compared to monocytes from “AA” protective allele carriers^30^. Crucially, the *in vivo* relevance of this assay is validated by the published observation that CD33 genotype-dependent differences in monocyte uptake of monomeric Aβ measured by this assay correlates well with the *in vivo* brain Aβ levels (as detected by amyloid PET) in the living donors of those monocytes^30^. We therefore compared uptake of Aβ monomers by monocytes from “CC” carriers (n = 9 unrelated donors) that were treated either with 60 nM wild-type CLU, or with desialylated CLU (desCLU) as a negative control. These experiments show that CLU treatment of CC monocytes inhibits Aβ uptake (Figure 5f). In contrast, desCLU treatment of monocytes from the same batch and same donor had no significant inhibitory effect on monomeric Aβ uptake (Figure 5f).

These results indicate that sialylated CLU, but not desialylated CLU, can suppress the engulfment of AD-relevant ligands (i.e. Aβ uptake) in human primary cells.

### CLU but not desialylated CLU inhibits phagocytosis of Aβ plaques by human monocytes

To further support this conclusion, we exploited the ability of human monocytes to clear human Aβ amyloid plaques in *ex vivo* brain slices from the 5XFAD transgenic mouse model of AD^78^ (Figure 6a-c). We note that CD33 modulates other microglial functions beyond engulfment of Aβ.

**FIGURE 6.**
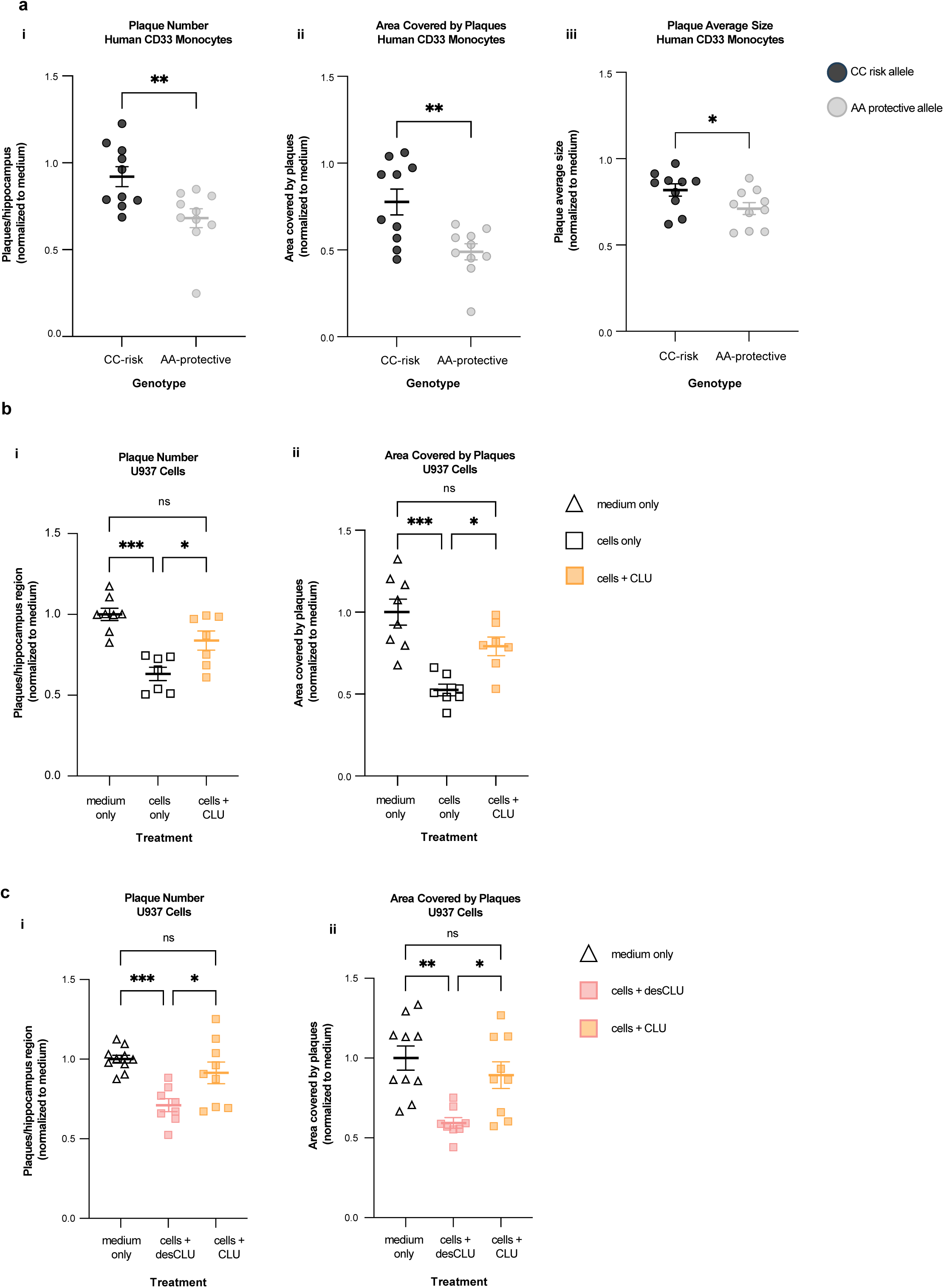
CLU down-regulates CD33-mediated amyloid plaque clearance by human monocytic cells. a. **Compared to monocytes from human CD33 “AA” protective allele carriers, monocytes from CD33 “CC” risk allele carriers** have reduced clearance of human amyloid plaques in *ex vivo* brain slices from 5XFAD transgenic human APP mice. Serial sections of unfixed brain from 5XFAD mice were incubated 48 hours with medium only, or with medium plus PBMCs from AA (protective) or CC (risk) allele carriers. Plaques were quantified by anti-Aβ immunofluorescent staining. The results were expressed as plaque number (i), brain area covered by plaques (ii) and plaque size (iii). The results were normalised to the results for medium only. N = 10 independent replications. ***p<0.001, *p<0.05; unpaired student’s t-test; error bars = mean ± SEM. Representative images are in Supplementary Data Figure 6A. **b. Human U937 monocytic cells clear amyloid plaques from *ex vivo* brain slices, and this is impaired by CLU.** Clearance of Aβ plaques from three serial sections of unfixed 5XFAD brain was quantified by immunofluorescent staining of hippocampal Aβ plaques following 48h incubation with medium only, with U937 cells only, or U937 cells + CLU. Results are expressed as number of plaques (i) or area covered by Aβ (ii), and were normalised to the result for slices exposed to medium only. *p<0.05, **p< 0.01, ***p<0.001; unpaired student’s t-test; error bars = mean ± SEM n = 8 replications. Representative images are in Supplementary Data Figure 6B. **c. Compared to CLU, desCLU does not downregulate clearance of Aβ plaques by human U937 monocytic cells. Clearance of Abeta plaques was quantified by** immunofluorescent staining of hippocampal Aβ plaques in three serial sections from 5XFAD brain following 48h incubation with medium only, with U937 cells plus medium containing desCLU, or with U937 cells plus medium containing CLU. Results are expressed as number of plaques (i) or area covered by Aβ (ii) and representative images are in Supplementary Data Figure 6C. *p<0.05, **p<0.01, ***p<0.001, one-way ANOVA Tukey’s multiple comparisons test; error bars = mean ± SEM, n = 10 replications.

These other microglial functions include release of microglial cytokines and clearance of neurotoxic membrane debris. However, our focussed assessment of Aβ plaque removal represents a tractable and disease-relevant assay of the impact of CD33 signalling on microglial function.

First, we validated the ability of human monocytes to remove human Aβ plaques in *ex vivo* brain slices from 6–8-month-old 5XFAD mice in a CD33-genotype-dependent manner. Briefly, monocytes are incubated with 20 µm coronal brain sections in RPMI medium for 48 hours at 37°C in the presence or absence of 60 nM human plasma-derived CLU. The sections are then stained and scored for amyloid plaque abundance and size. We find that compared to monocytes from “AA” protective allele carriers, monocytes from “CC” risk allele carriers only weakly clear amyloid plaques (as measured by the total number of plaques per hippocampus; the area covered by plaques; and plaque size; Figure 6a, Supplementary Data Figure 6A p < 0.001). This result nicely aligns with the previously reported reduced *in vivo* amyloid clearance (as measured by in vivo amyloid PET imaging) in CD33 “CC” risk allele carriers compared to protective CD33 “AA” allele carriers^30^.

Next, to determine how CLU might influence plaque clearance by human monocytes, we repeated this assay in the presence of 60 nM CLU or 60 nM desCLU (as a negative control). Because of the technical difficulty in obtaining sufficient numbers of human monocytes for this experiment, we had to use human U937 monocyte-like cells. U937 cells predominantly express CD33^M^ and mimic monocytes from homozygous and heterozygous “C” risk allele carriers (Figures 5a,b). This strategy also benefits from the fact that human U937 cells are a highly reproducible and widely accessible resource compared to primary PBMC cells.

We find that U937 cells significantly decrease the average size and number of amyloid plaques compared to medium only (Figure 6B, p <0.0001). This effect is inhibited when the U937 cells are co-incubated with physiologic concentrations (60 nM) of sialylated human CLU (Figure 6c, Supplementary Data Figure 6C, p <0.04). However, in good agreement with the biochemical studies described above, human desCLU induces significantly less immuno-inhibitory effect (Figure 6c, Supplementary Data Figure 6C, p < 0.02).

Together, these experiments validate a role for CLU ± Aβ oligomers both in modulating CD33 ITIM signalling and in regulating phagocytic clearance of amyloid plaques by myeloid macrophages. The CLU ± Aβ interaction with CD33 may also affect other AD-relevant microglial activities which cannot be tested in this assay. These other activities potentially include removal of neurotoxic membrane debris and secretion of inflammatory cytokines.

### CLU and CD33 genotypes interact to modulate AD risk

The biophysical biochemical and cell biological results described above raise the question of how the genotypes at both of these AD risk genes might interact to modulate AD risk and AD-related endophenotypes. To explore this question, we interrogated the comprehensive genetic, transcriptional, neuropathological and clinical phenotype data from the dorsolateral prefrontal cortex (DLPFC) for 1092 participants enrolled in the prospective ROSMAP cohort. Supporting the biophysical and biochemical analyses reported above, these genomic analyses reveal significant interactions between *CD33* and *CLU* gene expression and global AD pathology burden (p = 0.026), cognitive decline (p = 0.041), dementia with Lewy bodies (p = 0.007), and Lewy bodies in the neocortex (p = 0.018; n = 1092 participants; Supplementary Data Figure 6D). Causal mediation analyses reveal that CD33 mediates the association between CLU expression and both clinical AD dementia (p = 0.044) and amyloid burden (p = 0.006, n = 1092 participants; Supplementary Data Figure 6E). Expression quantitative trait locus (eQTL) analyses revealed *CD33* expression was significantly associated with the CLU SNP (rs9331947_G) (p = 0.005, n = 986 participants, Supplementary Data Figure 6F), but CLU expression was not associated with any CD33 SNPs (Supplementary Data Figure 6G). In addition, we examined the effect of *CD33* expression on AD traits in the presence of high and low CLU expression. We found that *CD33* was significantly associated with global AD pathology burden (p = 0.002), pathological AD (p = 0.0008), clinical AD dementia (p = 0.003), and β-amyloid levels (p = 0.005; n = 546 participants) in brains with low *CLU* expression (Table 2). Tangles, cognitive decline, and cerebral amyloid angiopathy were all nominally significant in the low *CLU* expression group as well. As discussed below, these genetic interactions are consistent with the biophysical, biochemical and cell biological studies reported here. They may also have important implications for personalised medicine approaches for management of people with CD33 and CLU risk alleles.

**TABLE 2.**
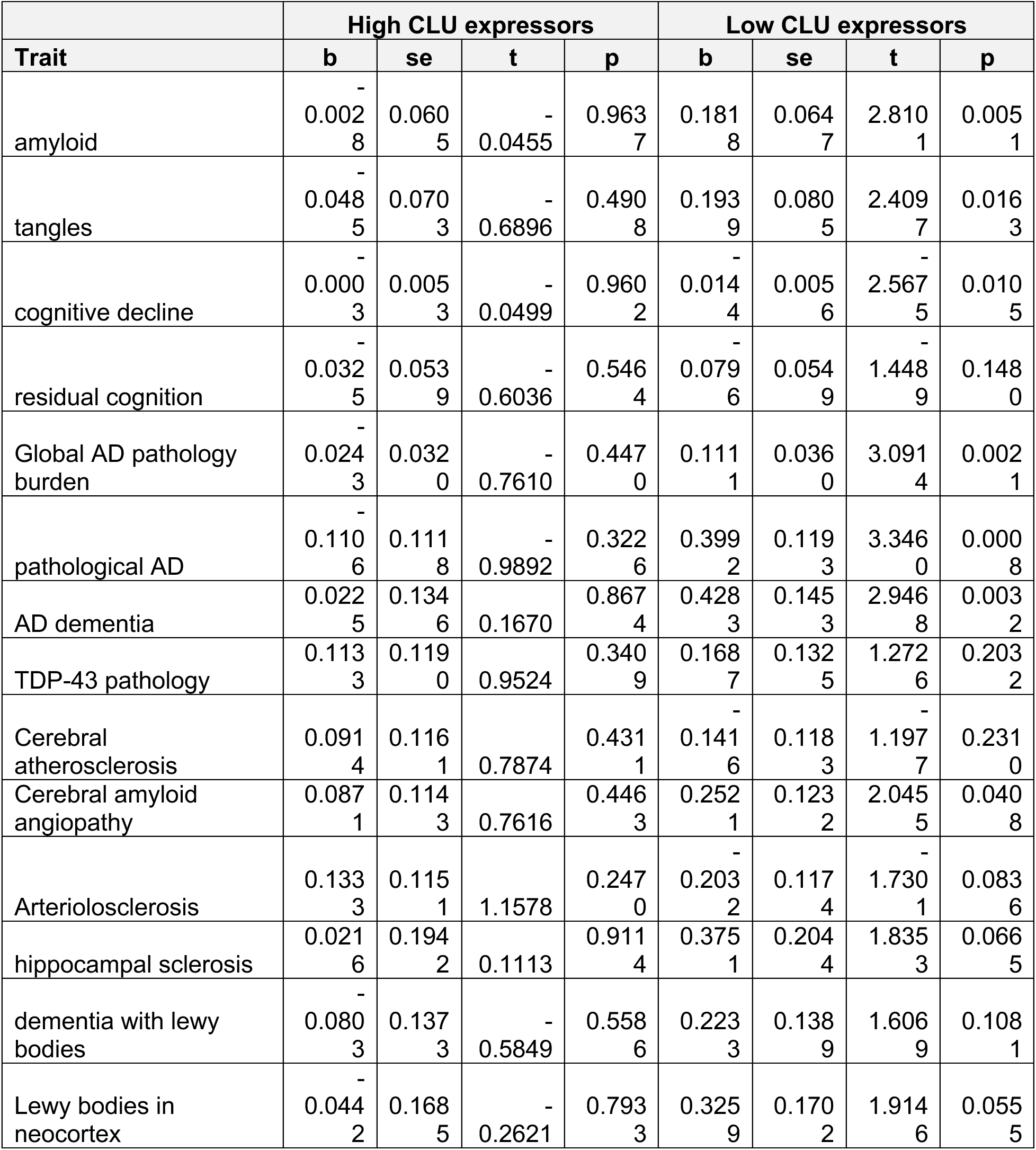
Association between CD33 gene expression and clinico-pathological traits of AD, stratified by CLU gene expression level. ROSMAP participants were stratified into two groups based on CLU gene RNA expression levels, using the median value (0.0083) as the cutoff: high CLU expressors (n = 546, CLU expression ≥ 0.0083) and low CLU expressors (n = 546, CLU expression < 0.0083). Within each CLU-defined subgroup, we tested whether CD33 gene RNA expression was associated with clinico-pathological traits of AD. Analyses were conducted in 1,092 participants with bulk RNA-sequencing data from the dorsolateral prefrontal cortex in the ROSMAP cohort, b: estimated effect size for CD33 gene expression on the trait; se: standard error of the estimate; t: t- statistic; p: p-value.

## DISCUSSION

The experiments described here offer several unexpected insights into the fundamental biology of the CD33 receptor and its role in the pathogenesis of AD.

The fundamental insights include the structure of the ligand binding site, the formation of dimers that facilitate trafficking of CD33^M^ to the cell surface, and the discovery of a shed sCD33^M^ fragment. The 2.24Å structure of the CD33 ectodomain delivers a detailed view of the sialic acid binding site and the spatial orientations of the Ig-like V-set and C2-set domains. The overall topology of the CD33^M^ sialic acid binding site is similar to those of other Siglecs, differing subtly in the architecture of the BC loop, CC′ loop and the orientation of the cognate Arg119. However, the more surprising feature of the CD33 structure is its dimerization via its C2-set extracellular domain. While this feature has not been explicitly emphasized for other Siglecs, some Siglecs (e.g. CD22) are thought to form functional nanoclusters at the cell surface^79^. Because there is a high degree of conservation of the residues in other human Siglecs at the equivalent position within CD33 dimeric interface, it will be interesting to now seek explicit evidence for dimerization in these other Siglecs.

The dimerization of CD33 likely has a profound effect on its biology, potentially contributing both to subcellular localisation and to ligand binding and specificity. Dimerization creates two ligand binding sites for each CD33 receptor, with the two ligand binding sites having divergent spatial orientations relative to the central axis of the receptor. This “open” configuration suggests that CD33^M^ might be specialised for binding larger sialylated molecules such as CLU. This possibility is supported by the strikingly higher binding avidity (Kd = 28.9 nM) for CLU (which has multiple sialylated residues) compared to those for simple sialyllactose molecules (Kd = 6.1 mM), which fit the CD33^M^ binding pocket well. We propose that the higher avidity between CD33^M^ and CLU is likely to be driven by the simultaneous engagement of both sialic acid binding sites on the CD33^M/M^ dimer with two glycan chains from the same CLU ligand molecule. It is conceivable that following the double glycan binding, there are additional interactions between the protein backbones of CD33 and CLU. If so, these additional interactions might contribute further to the high avidity binding of large sialylated ligands like CLU compared to small sialyllactose molecules.

We also demonstrate that dimeric full-length CD33^M^ is enriched at the cell surface whereas CD33^m^ dimers are not (see Figure 2). The molecular basis for the preferential enrichment of CD33^M^ dimers at the cell surface is not fully clear. Prior work has suggested that a peroxisome targeting sequence in the cytoplasmic tail of all splice forms of CD33 encourages partitioning of CD33^m^ into peroxisomes^39^. This is likely correct. However, because this putative targeting sequence is also present in the C-terminus of CD33^M^, it does not explain why CD33^M^ is not also targeted to peroxisomes. Conversely, the dimer site discovered in the crystal structure of the CD33^M^ ectodomain is predicted to also be present in CD33^m^. Future dynamic studies of the intracellular flux of various CD33 species will be required to resolve the underlying molecular mechanisms. One possibility is that motifs in the V-set domain contained in CD33^M^ (and CD33^MΔ4bp^ but not CD33^m^) may play a role in facilitating formation of CD33^M/M^ dimers (and CD33^M^/CD33^MΔ4BP^ complexes) at the cell surface (Figure 2e-f,k, Supplementary Data Figure 2J,K,M). However, regardless of the underlying mechanism, the enrichment of CD33^M^ at the surface of cells expressing either CD33^M^ only or both CD33^M^ and CD33^m^ cells is functionally reflected in their similar CD33^M/M^ dimer ITIM signalling properties. Specifically, in response to CLU ± Aβ oligomers, CD33^M/m^ and CD33^M/M^ cells display similar levels of CD33 ITIM tyrosine phosphorylation and SHP-1 recruitment (shown using transfected HEK293 cells - Figure 5c,d and Supplementary Data Figure 5C,D; and human AA vs CC monocytes - Supplementary Data Figure 5F top panel). This similarity provides a molecular explanation for the prior observation of equivalent clinical phenotypes in homozygous and heterozygous carriers of the rs3865444^CC^ risk allele (i.e. equivalently impaired Aβ clearance by *ex vivo* monocytes and equivalently increased *in vivo* brain amyloid loads)^30^.

Another remarkable observation arising from our structural and cell biological studies is the tight binding (and agonism) between CD33 and CLU, but the conspicuous absence of tight binding between CD33 and ApoE. Both ApoE and CLU are sialylated proteins that share other binding partners (ApoER2)^62,80^. Both also play a role in the pathogenesis of AD^81–86^. The present atomic structure of CD33^M^ does not immediately explain why CD33^M/M^ dimers preferentially bind CLU, but not ApoE. However, we speculate that the concurrent binding of a ligand to two binding sites on a CD33 dimer would require a specific configuration of the glycan chains on the ligand. The different affinities for CLU and ApoE could therefore reflect the specific orientation and/or type of glycans on CLU versus ApoE. In addition, as noted above, the spatial orientation of the CD33 CC’ loop may allow it to play a role in selective binding to specific larger sialylated ligands. Future studies using cryo-electron microscopy of CD33:CLU complexes (with or without Aβ oligomers) may provide some insights into the structural basis for the high avidity CD33:CLU interaction. Such information could be useful for generating small molecule inhibitors of CLU binding.

Another unexpected observation is that CD33^M^ undergoes shedding of its soluble N-terminal domain (sCD33^M^) both in HEK293 cells exogenously expressing the CD33^M^ cDNA, and in human iPSC derived microglia endogenously expressing the receptor (Figure 2i-k). Further proteomic and RNA-Seq studies will be required to fully exclude the possibility of differential splicing to account for the sCD33^M^ species. However, Intriguingly, the CD33^MΔ4bp^ mutant produces an extracellular soluble species that could mimic the effect of sCD33^M^. Our observation that both sCD33^M^ and CD33^MΔ4bp^ proteins can be co-immunoprecipitated with full length CD33^M^ and CD33^m^ proteins in HEK293 cells (Figure 2j), raises the possibility that both soluble truncated proteins could affect signalling by CD33^M^ itself, and possibly byother monocyte and microglial cell surface receptors (e.g. CD45 – see accompanying manuscript by Vo et al). Additional studies will be required to replicate this result in monocytes and microglia expressing endogenous CD33, and to understand the molecular/structural basis for this potential interaction (CD33^MΔ4bp^ does not contain the dimer interface sequence). It will also be of interest to determine whether sCD33^M^ and CD33^MΔ4bp^ proteins have similar biological properties. An analogous situation exists with the soluble extracellular domain of TREM2 (sTREM2), which protects against Aβ toxicity and AD progression^48,49,87–90^. For example, chimeric dimers containing CD33^M^: CD33^MΔ4bp^ or CD33^m^: CD33^MΔ4bp^ might support CD33^M^-type ITIM signalling (see Supplementary Data Figure 2L). Alternatively, sCD33^M^ and CD33^MΔ4bp^ proteins might interact with other cell surface receptors, perhaps activating additional deleterious signalling pathways. Crucially, because sCD33^M^ is a product of “risky” CD33^M^ but not “protective” CD33^m^, if sCD33^M^ (and CD33^MΔ4bp^) do have deleterious biological properties, their absence in cells predominantly expressing CD33^m^ could contribute to the “AD-risk effect” of CD33^M^ and CD33^MΔ4bp^, and to the apparent “anti-AD-risk effect” of CD33^m^.

A major outcome of our work is that it sheds light on how CD33 and CLU contribute to AD pathogenesis. Specifically, our discovery that physiological concentrations of sialylated CLU ± Aβ oligomers are high avidity ligands for CD33^M^ whereas ApoE is not, is of acute interest to the field. This unexpected observation emphasises the very different mechanisms by which CLU and ApoE modify AD risk. ApoE alters AD risk in part by impairing endolysosomal transport in an isoform-specific (not sialylation-specific) manner^91^. In contrast, we propose that CLU modulates AD risk in part by altering CD33^M^ ITIM signalling (as shown here). There are likely to also be CD33- independent mechanisms for CLU (e.g. blocking Aβ oligomerisation, protection of synapses and reduction of astrocyte reactivity^63,71,77,92,93^). Further work will focus on determining how the CD33: CLU interaction regulates the function of brain-resident microglia and infiltrating monocytes, and in peripheral blood monocytes.

The genetic and functional observations reported here provide powerful evidence for a previously unrecognised complex link between three genetic risk factors for AD: namely APP / Aβ oligomers, CLU and CD33. Specifically, our genetic studies support interactions between CLU and CD33 in modulating AD risk (Supplementary Data Figure 6E-G). This interaction is confirmed by the biophysical and cell biological studies which show that CLU binds CD33^M/M^ dimers (Figure 3-4) and activates ITIM signalling (Figure 5). This activation of CD33 ITIM signalling in turn leads to inhibition of microglial phagocytosis and plaque clearance (Figures 6). The QTL data suggest that this AD-risk promoting interaction is most profound at low CLU transcription levels. At higher CLU transcription rates, CLU appears to be protective. Future dynamic and kinetic systems biology experiments will be needed to characterise this complex relationship. However, we suggest that at lower levels of CLU expression, CLU-mediated activation of deleterious inhibitory CD33^M^ signalling might predominate. The CLU-mediated activation of CD33^M^ signalling may be saturable. If so, it is possible that at higher CLU expression levels generated by protective CLU allleles^93^, the saturable inhibitory CLU-CD33^M^ effect might then be overridden by the beneficial effects of higher CLU expression mediating the protective effects of CLU on Aβ, astrocytes and synapses ^63,71,77,92,93^. This complex interaction could be an important factor to include in future clinical trials and in personalised medicine approaches with therapeutics directed at CD33 or CLU.

Our work supports the existing notion that down-regulation of CD33^M^-based ITIM signalling is a potential target for preventative or disease-modifying therapies to mitigate deleterious monocyte and microglial responses to Aβ in AD. One such therapeutic approach already mentioned in the literature is the use of function-blocking anti-CD33^M^ antibodies^94–97^. However, given the difficulty in getting antibodies into the CNS and their occasional complication by amyloid-related imaging abnormalities (ARIA)^98^, other more tractable strategies might need to be considered. Specifically, our data provide 3D structures of the CD33^M^ ligand binding site and of the crucial dimer interface necessary for CD33^M^ stability and expression at the cell surface. These structures can now be used to design small molecule mimics to interrupt dimer formation. Such approaches will require the design of brain-penetrant molecules with higher affinity than the dimer site itself (10^-7^ M). If disruption of the dimer site needs to occur during assembly in the endoplasmic reticulum and Golgi, such molecules will also need to be cell penetrant. Candidate dimer-disrupting molecules include small molecules, peptides or aptamers. Additionally, antisense oligonucleotides designed to specifically down-regulate CD33 mRNAs containing exon 2 can be envisaged as a complimentary strategy to cripple CD33^M^ signalling. The restricted cell- and tissue-specific patterns of CD33 expression raise the hope that mechanism-based, non-CNS side-effects of such anti-CD33^M^ therapies could be minimal. Furthermore, the genetic analyses reported here suggest a personalized approach to the application of such anti-CD33 strategies might be especially useful. Specifically, persons with risk alleles at CD33 but with low expression eQTL alleles at CLU might be most responsive to anti-CD33 strategies.

## METHODS

### Protein expression and purification

A construct encoding residues 21-232 of CD33 (encompassing the V-set and C2-set immunoglobulin-like domains) was PCR amplified from an IMAGE clone (id 5217182) and inserted into pHLSec (a kind gift from Prof Radu Aricescu)^99^ via Gibson assembly. IMAGE clone 5217182 contains a sequence for CD33 corresponding to the R69G variant, the most common sequence observed in European (25%) and American (44%) populations (transcript CD33-201, www.ensembl.org). pHLSec encodes an N-terminal signal peptide (derived from RPTPσ) and a C-terminal hexahistidine tag. Protein was expressed by transient transfection of HEK 293F cells grown in suspension in Freestyle 293 medium (Life Technologies)^100^. Briefly, cells were centrifuged at 300 *g* and resuspended in fresh medium at 2.5 x 10^6^ cells/ml immediately prior to transfection. Plasmid DNA was added to the culture at a final concentration of 3 μg/ml and the culture incubated at 37°C for 5 min. Following incubation, polyethyleneimine at a concentration of 9 μg/ml (from a 1 mg/ml stock in 25 mM HEPES, 150 mM NaCl, pH 7.4) and the mannosidase I inhibitor kifunensine at 5 μM were added. 24 hours later, the culture volume was doubled, the HDAC inhibitor valproate added at 3 mM (to boost protein expression)^101^ and the kifunensine supplemented to return the concentration to 5 μM.

On Day 6 post-transfection, the culture was harvested and the medium mixed with 5 ml pre-washed Ni-Sepharose Excel resin (GE Healthcare) for 60 min at 4°C. The resin was poured into a column and washed with 200 ml Ni buffer A – 50 mM HEPES, 500 mM NaCl, 25 mM imidazole, pH 7.5. CD33 ECD was then eluted with Ni buffer B – 50 mM HEPES, 500 mM NaCl, 250 mM imidazole, pH 7.5 – concentrated to 10 ml in a Vivaspin 20 10K MWCO device and loaded on a Superdex 200 26/600 pg column pre-equilibrated in 20 mM HEPES, 200 mM NaCl, pH 7.5. Eluted CD33 ECD was subsequently concentrated to 1 mg/ml, buffer exchanged into 20 mM MES, 200 mM NaCl, pH 6.0 and incubated with 8,500 units Endo H_f_ per mg protein for 6 hours at RT. Endo H_f_ was removed by incubation with 1 ml amylose resin for 30 min and CD33 ECD was concentrated to 9 mg/ml and snap frozen in 100 μl aliquots in thin-walled PCR tubes in liquid nitrogen. The final yield was ∼ 60 mg/litre of culture.

Human plasma CLU used for BLI was purified from human plasma as described by Wilson et al.^102^. The purity analysed by Coomassie Brilliant Blue (CBB)-stained SDS PAGE is almost 100% (Extended Figure 3A). To remove conjugated sialic acid groups, purified human CLU (dialysed into sodium acetate buffer, pH 5.0) was supplemented with neuraminidase (from *Clostridium perfringens*, 1U, Sigma Aldrich) and incubated for 6 hours at 37°C with gentle mixing on an orbital shaker. Proteins were subsequently analysed by SDS PAGE (4-12% gradient Bis-Tris Bolt Plus pre-cast gel, ThermoFisher Scientific) and stained with Imperial^TM^ protein stain (ThermoFisher Scientific). Samples were also analysed by Western blot; 500 µg of treated and untreated hCLU were subjected to SDS PAGE and then electrophoretically transferred to a nitrocellulose membrane, which was blocked for 30 min with 2% (w/v) BSA in PBS containing 0.1% (v/v) Tween 20. Sections of the membrane were then cut and incubated with either G7 (hCLU-specific mouse monoclonal antibody^102^, as hybridoma culture supernatant), or 2.5 µg/ml recombinant human lectin Siglec-1 (6xHis-fusion protein which binds to sialic acid groups; R&D Systems, InVitro Technologies). After an overnight incubation at 4°C, the blots were washed three times with PBS containing 0.1% (v/v) Tween 20 and then incubated with the appropriate HRP-conjugated secondary antibody (0.32 µg/ml anti-6xHis-HRP antibody, Abcam; 0.4 µg/ml goat-anti-mouse-HRP, Dako, Agilent). Siglec-1 and HRP-conjugated antibodies were diluted in 2% (w/v) BSA in PBS containing 0.1% (v/v) Tween 20. Finally, the blots were washed as before and then developed using Pierce SuperSignal West Pico Chemiluminescent Substrate (ThermoScientific).

### Crystallisation and data collection

CD33 ECD was initially set up in 5 x 96 well vapour diffusion screens by an Innovadyne Screenmaker robot and crystals were identified in approximately 10% of drops. The best initial hit was in condition H9 of the MORPHEUS screen – 10% PEG 20,000, 20% PEG MME 500, 100 mM Bicine/Tris base pH 8.5, Morpheus amino acids. Optimised crystals grew as clusters of rods in 800 nl + 800 nl drops in 2% PEG 20,000, 4% PEG MME 500, 100 mM Bicine/Tris base pH 8.5, Morpheus amino acids over the course of 1-3 days. Individual rods were isolated, soaked for ∼20 s in mother liquor with PEG MME 500 supplemented to 15% for cryoprotection, and cryocooled by plunging into liquid nitrogen. Data were collected at 100 K on beamline I04-1 at the Diamond Light Source, a fixed energy beamline with a wavelength of 0.920 Å.

### Ligand soaking

3ʹ and 6ʹ-sialyllactose (Carbosynth Ltd) were dissolved in crystallisation mother liquor (2% PEG 20,000, 4% PEG MME 500, 100 mM Bicine/Tris base pH 8.5, Morpheus amino acids) at a final concentration of 20 mM. Crystals were transferred into 4 μl drops of either oligosaccharide solution and soaked for 10 days before soaking in cryoprotectant solution and cryocooling as for apo crystals. Data for the ligand-soaked crystals were collected on beamline I04-1 at the Diamond Light Source.

### Structure solution and refinement

Data were integrated and scaled using Xia2^103^ with the 3dii protocol (XDS^104^ for integration and processing, XSCALE for scaling), exhibiting an I/σI of 1.90 in the highest resolution shell (2.32 - 2.24 Å, Table I). Crystals were in the orthorhombic space group *P*2_1_2_1_2_1_ and the MATTPROB^105^ calculator suggested the presence of 4 copies of CD33 ECD in the asymmetric unit. The structure was solved using PHASER^106^ with a model derived by SCULPTOR^107^ from the structure of Siglec-5 (pdb id 2ZG2, 56% sequence identity)^8^. The initial model had a good fit to the density for the V-set domain, but a very poor fit for the C2-set domain. The model was improved by the application of autobuild^108^ from the PHENIX^109^ software suite, which reduced R_free_ to approximately 0.33. The model was then subjected to multiple iterations of refinement in phenix.refine^110^ coupled with manual rebuilding in COOT^111^, including the use of phenix feature-enhanced maps^112^. Torsion based non-crystallographic symmetry restraints were applied in phenix.refine, restraining dihedral angles between NCS-related copies. TLS groups were identified automatically using the phenix.find_tls_groups tool and applied during refinement. Where density was supportive of inclusion in the model, *N*-linked *N*-acetylglucosamine residues were added and linked to Asn residues using phenix.carboload. Some level of positive difference density was observed for all five potential *N*-linked glycosylation sites, suggesting that all are glycosylated within cells to some degree. However, *N*-acetylglucosamine residues were only modelled at positions Asn100, 160 and 209 (and not at Asn113 and Asn230). The final model was refined to an R/R_free_ of 0.19/0.23.

Data for the ligand-bound structures were processed and scaled as for the ligand-free structure and extended to a resolution of 2.66 Å (3ʹ-sialyllactose) and 2.48 Å (6ʹ-sialyllactose). The models were initially placed by rigid body refinement of the apo structure and similarly built and refined using iteration of COOT and phenix.refine. Where electron density supported building of ligands, monosaccharide residues were manually placed in COOT and refined in PHENIX. All structural figures were generated using PyMOL^113^.

### Analytical Ultra-Centrifugation

Sedimentation velocity (SV) analysis was performed in a ProteomeLab XL-I (BeckmanCoulter) analytical ultracentrifuge. Protein samples were loaded in 2-channel cells and centrifuged in an An-50 Ti 8-place rotor at 40,000 rpm, 20 °C for 15 hours. Absorbance at 280 nm was used for detection. The samples contained 0.4 mg/ml glycosylated or deglycosylated CD33 ECD, 20 mM HEPES, 200 mM NaCl, pH 7.5. SV data were analysed using Sedfit software (P. Schuck, NIH/NIBIB).

Sedimentation equilibrium (SE) experiments were performed in a ProteomeLab XL-I (Beckman Coulter) analytical ultracentrifuge. Samples of glycosylated and deglycosylated CD33 ECD at concentrations 0.2, 0.1 and 0.05 mg/ml were loaded in 6-channel equilibrium cells and centrifuged in an An-50 Ti 8-place rotor at 16,000 or 20,000 rpm and 20 °C until equilibrium was reached for each speed. Sample buffer was 20 mM HEPES, 200 mM NaCl, pH 7.5. SE data were analysed using HeteroAnalysis software (by J.L. Cole and J.W. Lary, University of Connecticut).

### Multi-angle light scattering

SEC-MALS was used to determine the mass of Endo H_f_-treated and non-deglycosylated samples of CD33 ECD. These were resolved on a Superdex 75 10/300 GL analytical SEC column GE Healthcare) in 20 mM HEPES, 200 mM NaCl, pH 7.5 and elution monitored using absorbance at 280 nm (Agilent 1200 MWD), light scattering (Wyatt Heleos II) and refractive index (Wyatt Optilab rEX). CD33 ECD samples were loaded at two concentrations: 2.5 mg/ml and 0.5 mg/ml (equivalent to 100 µM and 20 µM, CD33 ECD respectively). Due to the SEC step there is a significant dilution of sample concentration during the SEC-MALS experiment meaning maximal concentration achieved are > 10 times less than the loaded concentrations (i.e., < 10 and 2 µM CD33 ECD). In the case of the glycosylated samples, the masses of the protein and carbohydrate components were determined using the dual detection method as implemented in Wyatt’s ASTRA software as conjugate analysis. The protein refractive index increment^114^ based on the CD33 ECD sequence used was 0.1885 ml g^-1^ and the extinction coefficient for absorbance at 280 nm was 1252 ml g^-1^ cm^-1^. The *dn/dc* value used for carbohydrate^115^ was 0.145 ml g^-1^ and its UV absorbance at 280 nm zero. The inter-detector delay volumes and associated band broadening constants, as well as the detector intensity normalization constants for the Heleos and the UV intensity calibration, were determined as part of each set of measurements using a known protein standard (BSA).

QELS measurements (DLS) were obtained during SEC MALS runs using a fast response detector in the Helios instrument that replaces one of the static light scattering detector angles. Autocorrelation functions were fit to a translational diffusion of a single species (appropriate because of the SEC step) and converted to hydrodynamic radius (Rh) using a buffer viscosity of 0.93 cP using Wyatt’s ASTRA software.

### Blue-Native PAGE in cells expressing endogenous CD33

1 x 10^6^ THP-1 cells or peripheral blood monocyte cells from human subjects (obtained with institutionally approved informed consent protocols) were lysed in 100 µl PBS, 1% digitonin (plus protease inhibitors) on ice for 30 min. Insoluble material was pelleted at 20,000*g*, 5 min, 4 °C. 10 µl of the lysate was run on a BN-PAGE gel in 30 µl total volume, including G-250 additive at 0.08%. Protein was denatured in gel post-run by soaking in 1% SDS, 15 min and then blotted to PVDF. The membrane was blocked in 5% non-fat milk/PBS/0.1% Tween-20 and probed with rabbit anti- CD33 mAb EPR4423 at 1:1,000. Following washing, the blot was probed with goat anti-rabbit-HRP secondary at 1:20,000 and visualised with SuperSignal West Dura substrate on film.

### CD33 Dimer formation

The Myc-tagged CD33^M^ containing vector was obtained from Origene (RC207023L1). The vector for Myc-CD33^m^ was made by removing exon 2 using the following primers and the Q5 Site Directed Mutagenesis Kit (NEB). Fwd: ACTTGACCCACAGGCCCAAAATC; Rev: CTGCCCACAGCAGGGGCA. The Myc tag was then replaced with a 3X HA tag in both vectors, resulting in vectors for Myc-CD33^M^, Myc-CD33^m^, HA-CD33^M^ and HA-CD33^m^. 10 µg of paired vectors were transfected into HEK 293T cells using Lipofectamine 3000 in the following combinations: Myc-CD33^M^/HA-CD33^M^, Myc-CD33^M^/HA-CD33^m^, Myc-CD33^m^/HA-CD33^m^. After 48 hours, the proteins were immunoprecipitated with either anti-Myc (Cell Signaling, 2276S) or anti-HA (Abcam, 9110) antibodies and probed with the same antibodies or anti-CD33 (Santa Cruz, 28811) antibody.

### CD33:CLU Co-IP

10 cm dishes containing HEK 293T cells were transfected with Lipofectamine 2000 according to manufacturer’s protocol with pCMV6 plasmids containing either HA-CD33^M^or HA- CD33^m^-HA alone, or with a 50:50 mixture of pCMV6 plasmids containing either HA-CD33^M^ or myc- CD33^m^. After three days cells were harvested and incubated with or without 30 nM CLU purified from human plasma as previously described for 1 hour on a rotating wheel at 4 °C^92^. Cells were pelleted, extensively washed with PBS to eliminate excess CLU and lysed in 1%CHAPSO/TBS (100 mM NaCl, 50 mM TRIS, 1 mM EDTA, pH 7.4). Lysates were subjected to immunoprecipitation with rabbit anti-HA antibody and protein complexes were precipitated on a protein-G-sepharose matrix in 0.5% CHAPSO/TBS buffer. After 2 washes with IP buffer and one wash with PBS, proteins were eluted with SDS-PAGE loading buffer at 65°C for 10 minutes. Proteins were separated on a Novex NuPage Bis-Tris 4-12% gel and blotted onto a PVDF membrane. Membranes were probed for CLU with G7 and 41D mouse anti-CLU antibodies^92^, or mouse anti-HA antibody for CD33 tagged with HA.

### CD33/ApoE Co-IP

20µg wild type CD33^M^ or R119A ligand incompetent CD33^M^ proteins or ApoER2 ectodomain fused to His-tag ^66^ were incubated with 50µl ‘Dynabeads His-Tag Isolation and Pulldown’ (Thermo Fisher Scientific, Catalog # 10103D) at 4°C overnight. Beads were briefly spun down, and washed three times with IP buffer, followed by incubation with 500µl culture supernatant containing naturally secreted and receptor binding-competent ApoE3 or ApoE4 (3µg/ml) prepared as previously described^80,116^, for an additional 4 hour at 4°C. Beads were washed three times using wash buffer [500 mM NaCl, 15 mM TrisHCl, 0.5% Triton X-100 (pH8.0)]. The bound ApoE was separated on 4%-15% SDS-PAGE and analyzed by Western blotting using a polyclonal antibody directed against human ApoE.

### CD33 phosphorylation and SHP1 recruitment

HEK293 cells were transfected as above with pCMV6 plasmids containing HA-CD33^M^, or HA-CD33^m^ alone, or HA-CD33^M-^5X alone or with 50:50 mixtures of pCMV6 plasmids containing either HA-CD33^M^ or HA-CD33^m^ for two days. Cell media were removed and washed with PBS four times, then incubated with CLU 60 nM, Aβ 2µM,or CLU 60 nM plus Aβ 2µM for 0, 15 minutes, cells were lysed in 1X RIPA buffer with PhosSTOP cocktail tablet (Roche) and proteinase inhibitor (Sigma-Aldrich), 1mg total protein lysis buffer for each sample was performed for immunoprecipitation (protein concentration measured by BCA, Pierce), Rabbit HA antibody (ab9110, Abcam) was added and incubated for 1 hour at 4 °C, then Protein G beads were added and rotated for overnight at 4 °C. Beads were washed three times with 1XRIPA washing buffer, eluted in 1 X Nupage LDS buffer at 65 °C for 10 minutes, supernatant was subjected for western blot with mouse anti-phosphotyrosine antibody (PY20, ab10321, Abcam), or mouse anti-SHP1 antibody (14D5, Invitrogen), or mouse anti-HA antibody (H3663, Sigma-Aldrich).

### Bio-Layer Interferometry

Binding avidity of CLU to CD33^M^ ECD was analysed by bio-layer interferometry (BLI) using the Octet RED384 system (ForteBio). Assays were performed at 30 °C in the assay buffer (20 mM HEPES, 200 mM NaCl, 1 mg/mL casein, pH 7.5) with vibration at 1,000 rpm. Ni-NTA biosensors loaded with hexahistidine tagged deglycosylated CD33^M^ ECD wild-type (WT) or R119A were incubated with CLU in a range of 0.0025-5 µM to obtain the association curves. Equilibrium dissociation constant (Kd) was analysed using the averaged binding responses at 895-900 seconds.

Binding affinity of 3’-sialyllactose to CD33^M^ ECD was analysed by BLI at 30 °C with vibration at 1,000 rpm in the same assay buffer with CLU:CD33^M^ ECD interaction. Super Streptavidin biosensors loaded with 3’-sialyllactose-sp-biotin (Sigma Aldrich) were incubated with CD33^M^ ECD WT or R119A in a range of 15.6-250 µM to obtain the association curves, and subsequently in the assay buffer to obtain the dissociation curves. Kinetics data were analysed using a 1:2 bivalent analyte model.

### Microscale Thermophoresis

Microscale thermophoresis analysis was performed with the Monolith NT.115 instrument (NanoTemper) using excitation power of 20% at 23°C. Deglycosylated CD33 ECD was labelled with Alexa Fluor 647 via primary amine coupling. The proteins were diluted in 20 mM HEPES (pH 7.5), 150 mM NaCl, 0.05% Tween-20. Kd values were calculated using the MO Affinity Analysis software.

### Far-UV circular dichroism (CD)

Far-UV CD spectra were recorded on a Jasco J-720 spectrometer using a 1 mm path length cuvette at 4 °C. The wild-type (WT) CD33^M^ ECD and R119A (0.2 mg/mL) were prepared in a buffer (20 mM HEPES, 200 mM NaCl, pH7.5). CD spectra were obtained from 205 to 260 nm, with a 1- nm bandwidth, and 8-sec response time and a scan speed of 20 nm/min. Data are averaged from five consecutive scans.

### Flow Cytometry

HEK 293T cells were plated into 6-well plates one day before transfection. Cells were transfected with GFP (1µg) plus CD33^M^ (2 µg), CD33^m^ (2 µg) or CD33^M^+CD33^m^ (1 µg+1 µg) plasmids using Lipofectamine3000. Twenty-four hours after transfection, cells were collected by trypsinization and then washed twice with 1xPBS. Cells then were stained by Viability Dye eFluor450 (65-0863, eBioscience) at 4°C for 30 min. After washing, cells were resuspended with 1x PBS and 1 million cells per test were then stained with primary antibodies (WM53-PE and A16121H for the double staining; HIM3-4-PE for the single staining) for 30 min at 4°C. After washed with 1x PBS 3 times, double stained cells were stained with secondary antibody (anti-ratIgG2a-AF647) for 30 min at 4°C. After being washed 3 times, cells were transferred to flow cytometry test tubes and data were acquired by BD FACS LSRII with corresponding compensation controls. Data were analysed by FlowJo v10. GFP positive cells were analysed after sequential gating to exclude debris, doublets, dead cells, GFP negative cells and cells with very high GFP expression that could not be compensated properly.

CD33^M-5X^ cells were not included in the flow cytometry experiments because the 5X dimer site mutant disrupts the recognition motif for the WM53 anti-CD33 antibody used to study CD33^M^ in fluorescence flow cytometry. These cells were studied by surface labelling.

### SNAP-tagged plasmid construction for smFRET studies

Plasmids coding for various CD33 receptors tagged at their amino-terminus with the catalytically enhanced version of the SNAP tag - SNAP_fast_ (SNAP_f_) were constructed using standard subcloning methods by replacing the dopamine D2 receptor coding region of the previously described pcDNA5/FRT/TO-IRES SNAP_f_-D2 vector^117^ with the following coding regions: human CD33^M^ (positions 17-364); human CD33^M-5X^; and human CD33^m^ (positions 13-237). These vectors contain a weak Kozak sequence with thymine at the -3 and +4 positions and a crippled CMV promoter in which the enhancer sequence was partly deleted. All constructed and commercial plasmids were confirmed by whole-plasmid DNA sequencing (Plasmidsaurus).

### Cell culture and generation of stable CHO cell lines for smFRET studies

All Chinese Hamster Ovary (CHO) cell lines were maintained in Ham’s F-12 medium (Corning), 10% Fetal Bovine Serum (Corning), 2 mM L-glutamine (Corning) at 37 °C in 5% CO_2_. For parental LEx-Flp-In T-REx (FITR) CHO cells, a previously described FITR cell line containing an FRT recombination target site^118^ in the CHO genome that confers low expression levels and that also allows for tetracycline-inducible expression^41,119^, the medium contained 15 μg ml^−1^ blasticidin (InvivoGen) and 50 μg ml^−1^ zeocin (Invitrogen). For stable LEx-FITR CHO lines expressing CD33 receptors, the medium contained 500 μg ml^−1^ hygromycin (Thermo Fisher Scientific) and 15 μg ml^−1^ blasticidin. All CHO lines were grown to ∼70-80% confluency in six-well tissue culture dishes before transfection for generating stable lines or tetracycline (MilliporeSigma) induction and/or labelling for microscopy.

To generate stable CHO cell lines expressing SNAP_f_-CD33M, -CD33M-5X, and -CD33m, 200 ng of the pcDNA5/FRT/TO-IRES vectors encoding these receptors was co-transfected with 1,800 ng of the pOG44 Flp-In Recombinase vector (Invitrogen) into the LEx-FITR CHO lines using Lipofectamine LTX according to the manufacturer’s protocol. Stable cells were selected with 500 μg ml^−1^ hygromycin (Thermo Fisher Scientific) in the presence of 15 μg ml^−1^ blasticidin.

### Sample preparation for TIRF Microscopy

The LEx-FITR CHO stables expressing SNAP_f_-CD33^M^, -CD33^M-5X^, or -CD33^m^ were induced with tetracycline (0, 200, and 1000 ng.ml^-1^ for SNAP_f-_CD33^M^, -CD33^m^, and-CD33^M-5X^ expressing stable lines, respectively.) 18–24 hours before labelling and imaging to yield comparable cell surface expression levels suitable for resolving single molecules. The CHO cells were then prepared for single-molecule imaging by first stochastically labelling with the membrane-impermeant, self-healing SNAP_f_-reactive Lumidyne dyes LD555p-benzylguanine (BG) (Acceptor, 666 nM) and LD655-BG (Donor, 333 nM)^119,120^in conditioned culture medium for 20 minutes at 37 °C in 5% CO_2_ under dark conditions. The donor-only control was labelled with 333 nM LD555p- BG in the absence of the acceptor dye following the same procedures above. After labelling, cells were extensively washed with Dulbecco’s Phosphate-Buffered Saline (DPBS) supplemented with 5 mM Glucose (Sigma Aldrich) and 1 mg.ml^-1^ Bovine Serum Albumin (BSA; Sigma Aldrich) and then dissociated with PBS-based enzyme-free cell dissociation buffer (Sigma Aldrich). Cells were then centrifuged at 500 *RCF* for 5 minutes, resuspended in FluoroBrite DMEM (ThermoFisher) and seeded onto fibronectin-coated (10 ng µl^−1^, Sigma Aldrich) high-index glass coverslips (1.7 numerical aperture (NA) HIGHINDEX-CG 20 mm diameter, Olympus) and allowed to adhere for 60 minutes at 37 °C in 5% CO_2_. Prior to use, the coverslips were chemically cleaned in an ultrasonic bath sonicator (CPX3800, Branson Ultrasonics) using the following sequence of steps: 20 minutes in 10% Alconox (Sigma Aldrich), 10 minutes in ultrapure water, twice for 40 minutes in potassium hydroxide (1 M, Sigma Aldrich), followed by 20 minutes in ultrapure water. Once chemically cleaned, the coverslips were further processed under argon plasma (Zepto RIE, Diener Electronic) for 5 minutes and then coated with 10 ng.μl^-1^ fibronectin (Sigma Aldrich) for 60 minutes before seeding with labelled cells.

After the cells adhered onto the coverslips, and immediately before imaging, they were washed with excess DPBS and then assembled onto a microscopy chamber^117^, where they were incubated at room temperature for 30 minutes in dark conditions in imaging buffer containing 1 mM 3,4-dihydroxybenzoic acid (PCA) and 50 nM protocatechuate 3,4-deoxygenase (PCD) (Sigma-Aldrich) to deplete dissolved ambient oxygen by 50% as previously described^119^.

### TIRF microscopy and smFRET imaging

Image sequences were collected at a time resolution of 15 ms using a customized, objective-based IX81 TIRF microscope equipped with a cellTIRF-4Line illuminator system (Olympus Life Sciences/Evident) containing a custom microscope enclosure to create a controlled, lightproof environment (Precision Plastics Inc.). A 100× oil-immersion 1.70-NA (APON100×HOTIRF, NA 1.70, Olympus) objective was used with a 1.705 NA immersion oil (Cargille Labs) for imaging the samples. An evanescent TIRF field with an approximate penetration depth of 80 nm was used to excite labelled receptors at the proximal plasma membrane using a 532-nm (Opus 3600, Laser Quantum, 3W)^41^ and/or 640-nm (LAS-640-100D, Cell^tirf^, Olympus, 100 mW) lasers. Fluorescence emission was separated from the excitation light using a dual-band laser filter set (ZT523/640rpc, ZET532/640m, ZET532/640x, Chroma) in combination with an emission side image splitter (OptoSplit 2, Cairn) equipped with a FRET filter set (ZT640rdc, ET585/65m, ET655lp, Chroma) to spatially separate and project donor and acceptor emission signals side-by-side onto a single quantitative complementary metal oxide semiconductor (qCMOS, ORCA Quest C15550-20UP, Hamamatsu Photonics) camera for detection. The dual-band TIRF–FRET filter configuration allows for the excitation of donor and acceptor fluorophores separately or simultaneously. The microscope is also equipped with nano- and micro-drive stage positioning systems (Mad City Labs Inc.) for locating single living cells.

Before smFRET imaging, cells were briefly excited with both 532-nm and 640-nm laser lines at 10 mW measured at the back focal plane to generate an initial image for quantifying the density of donor- and acceptor-labelled CD33 receptors. SmFRET imaging was then performed by exciting the same cells with only the 532-nm laser (10 mW at back focal plane) and acquiring 5,000 frames per image sequence in both donor and acceptor channels.

### Determination of surface density by single-particle detection

The surface density of labelled molecules was determined from an initial image of cells taken from direct and simultaneous excitation before smFRET imaging. The number of donors and acceptors within a region of interest for each cell was determined by the DoG particle-detection function of TrackMate in ImageJ^121,122^. The cell surface area and region of interest was determined by boundary tracing on the projected image, generated from the donor image stack using the ImageJ plugin ZProject with projection type ‘Standard Deviation’^122^.

### Tracking smFRET in living cells

The smFRET image sequences were processed using the previously described and publicly available smCellFRET platform ^3^ for tracking and subsequent analysis of smFRET in living cells to detect and quantify the number of sensitized acceptor events and to generate their corresponding smFRET trajectories, fluorescence and smFRET time traces. The tracking constants for donor and acceptor are shown in **Supplementary Table 1**; all other calculations were performed as previously reported ^3^. For freely diffusing-trajectory (FDT) analysis, freely diffusing smFRET trajectories, as well as freely diffusing segments from smFRET trajectories with more than one diffusion state, were determined by divide-and-conquer, moment-scaling spectrum (DC-MSS) analysis^41,123,124^.

### Plotting and statistics for smFRET data

Plotting, distribution fitting and statistics for all single-molecule data were carried out using GraphPad Prism (Version 10.3.1 (509) GraphPad Software LLC). To determine P values, ordinary one-way ANOVA with Šídák’s multiple comparisons test was performed for multisample comparisons.

### CD33^MΔ4bp^ co-IP

HEK293 cells were transfected as above with single pCMV6 plasmids containing: i) empty vector (control) alone; or ii) HA-CD33^M^ alone; or iii) HA-CD33^m^ alone; or iii) CD33^ΔM4bp^ alone; or iv) with equal amounts of two pCMV6 plasmids, one of which contained HA-CD33^MΔ4bp^ and the other contained either HA-CD33^M^ or HA-CD33^m^. Cell were grown for two days. Supernatant media were removed. Cells were washed with cold PBS and lysed in 1X IP lysis buffer with PhosSTOP cocktail tablet (Roche) and proteinase inhibitor (Sigma-Aldrich). Equal amounts of cell lysates from each sample were used for immunoprecipitation. Anti-HA magnetic beads (SAE0197, Sigma-Aldrich) were added and incubated for overnight at 4 °C. Beads were washed three times with 1X IP lysis buffer, eluted in 1 X NuPage LDS buffer at 70°C for 10 minutes. The supernatant were western blotted and probed with rabbit anti-CD33 Ig-like V type domain antibody (ab134115, Abcam).

### hiPSC Maintenance

The FA0000010 (FA10) hiPSC line was generously provided by Barbara Corneo^125^. Undifferentiated hiPSCs were cultured in StemFlex medium (ThermoFisher) supplemented with penicillin/streptomycin (ThermoFisher) on reduced growth factor Cultrex BME (Bio-Techne). Cells were routinely passaged 1–2 times per week using ReLeSR (STEMCELL Technologies) without ROCK inhibitor (Y-27632, Selleck Chemicals) as previously described^126^.

### iPSC-derived Microglia (iMG) Differentiation

hiPSCs were differentiated into iMGs following previously established protocols with minor modifications^126–128^. Hematopoietic precursor cells (HPCs) were generated using the STEMdiff Hematopoietic Kit (STEMCELL Technologies), following the manufacturer’s instructions. Floating HPCs were gently harvested and cryopreserved at three separate time points: days 11, 13, and 15 (or alternatively, days 12, 14, and 16). Previously cryopreserved HPCs were terminally differentiated into iMGs by plating at a density of 60,000 cells/cm² on reduced growth factor Cultrex BME in microglia differentiation medium composed of DMEM/F12 supplemented with 2X insulin- transferrin-selenium, 2X B27, 0.5X N2, 1X GlutaMAX, 1X non-essential amino acids, 400 µM monothioglycerol, and 5 µg/mL human insulin (ThermoFisher). The medium was freshly supplemented every other day with 100 ng/mL IL-34, 25 ng/mL M-CSF (R&D Systems), and 50 ng/mL TGFβ1 (STEMCELL Technologies) through day 24. On day 25, 100 ng/mL CD200 (Bon Opus Biosciences) and 100 ng/mL CX3CL1 (R&D Systems) were added to the existing supplement regimen and maintained for the remainder of the culture period. For all experiments in this study, iMGs were stimulated and collected between days 35–37.

### CLU Stimulation and Lysate/ Media Collection

Native human plasma-derived clusterin (BioVendor R&D) was reconstituted in sterile water according to the manufacturer’s instructions. iMGs were treated with 300 nM clusterin or sterile water in complete microglia medium for 24 hours. After 24 hours, media was collected, centrifuged at 14,000 rpm for 10 minutes at 4°C, aliquoted, and snap-frozen on dry ice. iMGs were lysed using Pierce™ IP lysis buffer (ThermoFisher) supplemented with 1X Halt™ Protease and Phosphatase Inhibitor Cocktail and 1 mM sodium orthovanadate (New England Biolabs).

### Full length and soluble CD33 IP from human iMG

CD33 antibody (ab134115, Abcam) and protein A Sepharose beads (GE17-0780-01, Millipore Sigma) were incubated at 4 °C overnight. After washing 2 X with 0.2 M Na Borax/Boric acid buffer (pH 9.0), beads were suspended in 0.025 M Borax (pH 9.45) containing 5.2 mg of DMP (dimethyl pimelimidate, D8388, Sigma-Aldrich) and incubated for 30 minutes at RT. Beads were washed and incubated with 0.2 M ethanolamine (pH8.0) for 2 hours at RT to stop crosslinking. Uncrosslinked antibodies were removed and beads were washed sequentially with 10 mM NH_4_HCO_3_, 100 mM glycine, 100 mM Tris/HCl (pH 8.0) and PBS. The crosslinked beads were kept at 4°C and used for the following CD33 IP.

Cell lysates and media supernatants from cultures iMGs were incubated overnight at 4°C with crosslinked beads. Beads were washed 3 X with 0.5X IP lysis buffer and 1X with H_2_O and then eluted three times in 0.3% trifluoroacetic acid at RT for 5 min. Eluates were neutralized with 1M Tris buffer, and then concentrated by speed vacuum. The concentrated IP products were mixed with 4XNuPage LDS buffer, heated at 70°C for 10 minutes and then subjected to western blotting with mouse anti-CD33 Ig-like C2 type domain antibody (PWS44, Sigma-Aldrich).

### Proximity Ligation Assay (PLA)

The PLA assay was performed on formalin-fixed paraffin-embedded brain slices from three individuals with Alzheimer’s disease from the Maritime Brain Tissue Bank following the kit instructions (Duolink *In Situ* Detection Reagents Orange, Sigma Aldrich DUO92007). Briefly, after deparaffinization and antigen retrieval with sodium citrate buffer (pH6.0), tissues were permeabilized and blocked non-specific binding then the primary antibodies, rabbit anti-human CD33 (Sigma Aldrich HPA035832, 1:100) and mouse anti-human clusterin-a (Santa Cruz sc- 32264, 1: 50), were performed overnight at 4°C. After several washes with PBS-T (0.1% tween), secondary antibodies conjugated with oligonucleotides (PLA probe mouse PLUS and PLA probe rabbit MINUS, Sigma Aldrich) are added to the reaction and incubated. The samples were washed with TBS-T (10mM Tris, pH 7.5, 150mM NaCl, 0.05% tween, prepared freshly) two times. The Ligation solution, consisting of two oligonucleotides and Ligase, is added and the oligonucleotides hybridize with the two PLA probes and join to a closed circle if they are in proximity theoretically ranging from 0 to 40 nm. Then the samples were washed with TBS-T two times. The Amplification solution, consisting of nucleotides and fluorescently labelled oligonucleotides, was added together with Polymerase. The samples were washed with wash buffer B (0.2 M Tris, pH 7.5, 0.1M NaCl) three times then, incubated with primary antibodies, rabbit anti -Iba1 (Wako, 01919741, 1:100) and mouse anti-Aβ (Creative Biolabs, NAB61, 1: 50) for 1.5hr at room temperature or overnight at 4°C. After several washes, samples are incubated with secondary antibodies, goat anti-rabbit IgG Alexa Fluor 488 (Jackson ImmunoResearch, 1:200) and goat anti-mouse IgG Alexa Fluor 647 (Jackson ImmunoResearch, 1:200). After several washes with PBS, samples were incubated with 1X TrueBlack for 30 seconds and washed three times with PBS then mounted with Duolink *In Situ* Mounting Medium with DAPI (Sigma Aldrich). The PLA signal was detected with Cy3 filter by confocal fluorescence microscopy (Zeiss LSM880) and analyzed by ZEN (Zeiss).

### Antibodies used

The following antibodies were used: **Myc:** (Cell Signalling, 22776S); **HA:** (rabbit polyclonal, Abcam, ab9110; mouse monoclonal clone HA-7, Sigma-Aldrich, H9658; mouse monoclonal, Sigma-Aldrich, H3663 ); **CD33:** (Santa Cruz, 28811; rabbit monoclonal, Abcam, Ab134115; mouse monoclonal, Millipore Sigma, 133M-1, (PWS44); mouse monoclonal EMD Millipore, CBL163 (Clone WM53); eBioscience, 12-0339-41; eBioscience, 12-0338-42 (WM53-PE); Biolegend, 94835 (A16121H - not yet commercially released); eBioscience, #12-0339-41 (HIM3-4-PE); rabbit polyclonal, Sigma Aldrich HPA035832); **SHP1:** (Thermo Fisher Invitrogen 14D5); Anti-phosphotyrosine (PY20, Abcam Ab10321). **CLU**: (The anti-clusterin monoclonal antibodies were obtained from Mark Wilson (G7 and 41D hybridoma); mouse monoclonal, Santa Cruz sc-32264, clone 78E); **Iba1:** (rabbit polyclonal, Wako, 01919741,); **Aß:** (mouse monoclonal, Creative Biolabs, TAB0792CLV, clone NAB61); **Isotype control**: **rat IgG2a** (Biolegend #405416, AF647, 1:80 for flow cytometry); **mouse IgG1k** PE (eBioscience #12-4714-82, 1:80 for flow cytometry); **mouse IgGκ** (from Wilson Lab for PLA); **rat IgG2ak** (Biolegend #400501, for flow cytometry); **mouse IgG1** (EMD Millipore MABC002, for CBL163 WM53 control). All antibodies have been used in previously published work by other groups.

### CLU and CD33-mediated modulation of degradation of amyloid plaques in ex vivo brain slices

Coronal brain sections (20µm) were prepared from 6-8 month old 5XFAD mice (PMID: 17021169 ) using a cryostat as previously described (PMID: 21280087). U937 monocyte cell line 3 x10^5^ cell per 100µl in RPMI 1640(biological industries, 01-100-1A) were plated on unfixed brain sections and incubated at 37°C 5% CO2 for 48 hours with or without CLU (60nM) or 10µg/ml WM53 anti-CD33 antibody or isotype control. Quantification of hippocampus Aβ burden was done as described (PMID: 21280087). Sections were washed three times with PBS (Sigma D8537) and fixed with 4% PFA (paraformaldehyde) for 10 minutes. Sections were washed three times for five minutes with PBS. Immunostaining was done with mouse anti-human 6E10 antibody (1:500) (biolegend;cat 803001) for 1 hour at room temperature. Sections were than washed three times with PBS and stained for goat anti-mouse secondary antibody FITC- conjugated (Invitrogen; A11001) for 1 hour at room temperature. Sections were washed three times for five minutes with PBS and quantification was done in a blinded fashion using Imaging Research software from the National Institutes of Health in an unbiased stereological approach.

### Collection of Human Peripheral Blood Mononuclear Cells

De-identified blood was collected from the New York Blood Center (NYBC) or Partners’ Crimson Biospecimen Repository Core. All blood draws and data analyses were done in compliance with protocols approved by the Institutional Review Boards of each institution. Both males and females were used. Monocytes were separated by Lymphoprep gradient centrifugation (StemCell Technologies). Monocytes were frozen at a concentration of 2 × 10^7^ cells/mL in 10% DMSO (Sigma-Aldrich)/90% foetal bovine serum (vol/vol, Corning). After thawing, Monocytes were washed in 10 ml of phosphate-buffered saline according to a previously published protocol (Ryan et al., 2017). Samples were genotyped for rs8356444 using primers and the TaqMan SNP genotyping assay from Applied Biosystems (now ThermoFisher Scientific).

### Amyloid-β (Aβ 1-42) uptake assay

Monocytes were isolated from peripheral blood mononuclear cells (PBMC) by using anti- CD14+ microbeads (Miltenyi Biotech). 100,000 monocytes per well were plated in RPMI with 1% fungizone, 1% penstrep, and 10% FBS in a 96-well plate. The following final concentration was used for the Aβ 1-42 uptake assay: 60nM Clusterin (BioVendor, NC), 250nM beta-Amyloid (1-42), HiLyte™ Fluor 647-labeled (Anaspec), 60nM desialylated CLU. Clusterin or desCLU was preincubated with Aβ 1-42 for 15 min. in the cell culture incubator before being added to the monocytes. For the uptake assay, the preincubated clusterin and Aβ 1-42 mixture was added to the cells and incubated for 1 hour at 37°C. A total of n=10 individuals were used in each treatment condition. After the incubation, cells were washed using staining buffer (1% FBS in PBS) and then followed by incubation with fixable dead cell stain (ThermoFisher, L34955). Cells were fixed with 4% PFA. Aβ 1-42 uptake was measured by using NovoCyte Quanteon Flow Cytometer (Agilent, Santa Clara, CA).

### flowPLA

The flowPLA assay was performed as per the manufacturer’s instruction (Sigma Aldrich). Briefly, primary monocytes were permeabilized for 10 minutes on ice, then fixed for 30 minutes on ice using the Invitrogen Fix & Perm Cell Permeabilization Kit. In the first antibody incubation, rabbit anti-human CD33 (Bioss, bs-2042R, 1:400) and mouse anti-human SHP-1 (Thermo Scientific, MA5-38614, 1: 400), were performed overnight at 4°C. After two washes with Duolink wash buffer (Sigma Aldrich), secondary antibodies conjugated with oligonucleotides (PLA probe MINUS and PLA probe PLUS, Sigma Aldrich) were added to the reaction and incubated for one hour at 37 °C. The samples were then washed with Duolink wash buffer twice. The Ligation solution, consisting of two oligonucleotides and Ligase, was added and the oligonucleotides hybridized to the two PLA probes. Then the samples were washed with Duolink wash buffer two more times. The Amplification solution, consisting of nucleotides and fluorescently labeled oligonucleotides, were added together with the polymerase. The oligonucleotide arm of one of the PLA probes acts as a primer for a rolling-circle amplification (RCA) reaction using the ligated circle as a template, generating a concatemeric (repeated sequence) product. The fluorescently labeled oligonucleotides will hybridize to the RCA product. The samples were washed with Duolink wash buffer and then incubated with the Duolink flowPLA Violet Detection solution. The signal is visible as a distinct fluorescent spot and analyzed by flow cytometry (Novocyte Quanteon Cell Analyzer).

### Statistical Analysis of CLU-CD33 interaction in the ROSMAP data

CLU-CD33 gene expression interactions were investigated using the ROSMAP RNA-sequencing and GWAS data that have been previously described to examine the interplay between CLU and CD33 using three models^129,130^. When ROSMAP participants are enrolled, they are not known to have dementia and have agreed to detailed clinical evaluation and brain donation at death^131^. ROSMAP clinical and pathologic methods of characterization have been previously described^132–137^. Linear regression models for quantitative traits and logistic regression models for binary traits were used to test the association of CLU gene expression with CD33 expression in the dorsolateral prefrontal cortex (DLPFC) with AD neuropathological measures. The gene expression levels were adjusted for age, sex, RIN score, postmortem interval, and other technical covariates for RNA-sequencing and the residual expression values were used for interaction testing. A quantitative trait loci approach was used to test whether genetic variation in CLU regulates CD33 expression levels and vice versa. Common SNPs (MAF>0.01) were tested for association with residual gene expression levels in a linear regression model adjusting for age, sex, and population substructure variables. Lastly, the R mediation package was used to test whether the association of CLU expression with clinical AD, AD neuropathology and cognition is mediated by CD33 levels. This package was also used to test mediation in the other direction (CLU mediates CD33 association with traits) to establish causality. Additionally, we tested the association of CD33 with AD neuropathology independently in the top and bottom 50^th^ percentiles of CLU expression levels. This was to assess the difference in CD33 association with AD neuropathology in the presence of high or low CLU expression.

### Sex and Gender

Both male and female human brain samples and human peripheral blood monocytes were used in this study. Female animals were used in the plaque phagocytosis studies. U937 cells are male cells

### Human Subjects

Monocytes were derived from the peripheral venous blood of healthy control volunteers from the BWH blood bank. Samples were genotyped for rs8356444 using primers and the TaqMan SNP genotyping assay from Applied Biosystems (now ThermoFisher Scientific). All blood draws and data analyses were done in compliance with protocols approved by the Institutional Review Boards of each institution. Anonymized data were used from individuals enrolled in the ROSMAP study.

### Statistical Analyses

For simplicity, p-values are reported in figures in the style p<0.01. The exact p-values and statistical tests are reported in Supplementary Data Table 7.

## DATA AVAILABILITY

**Accession Numbers:** Coordinates and structure factors have been deposited in the Protein Data Bank under accession codes 5IHB (Apo CD33), 5J06 (3-SL complex) and 5J0B (6-SL complex).

**Source Data:** has been deposited in Figshare at: https://doi.org/10.6084/m9.figshare.29604716.

**Single Molecule Image Data Availability:** The raw single-molecule image data generated and analyzed that support the findings of this study are available from WBA, JAJ upon request. These image data are not deposited in a public database because of their large file sizes.

## CODE AVAILABILITY

The smCellFRET data analysis pipeline is freely available for academic use. The software and updated versions can be downloaded at http://innovation.columbia.edu/technologies/CU15268.

Other software used to collect and analyze data for this work as described in the Methods either was published previously or is commercially available.

## ACKNOWLEDGEMENTS

We thank the staff at the Diamond Light Source including Alex Dias and Alice Douangamath for advice on data collection, Prof Randy Read for discussions during preparation of the manuscript, and Ms Christina Yung for technical assistance in the early phase of the work. This work was funded by grants from the Ludwig Family Foundation, Wellcome Trust Collaborative Award 203249/Z/16/Z, Zenith Award of Alzheimer Association ZEN-18-529769 (PHStGH); Canadian Institutes of Health Research (PHStGH 406915, PEF, MI), National Science and Engineering Research Council (MI); Canada Foundation for Innovation (MI); Princess Margaret Cancer Fund (MI); UK Medical Research Council (PHStGH); Canada Research Chair in Cancer Structural Biology (MI) and National Institutes of Health (R01AG043617 RFIAG058852 (EMB) T32AI148099 (MR), U01AG072572 and U01AG072572-04S1 (PDJ, PHStGH). DAB (ROSMAP) is supported by P30AG10161, P30AG72975, R01AG17917. R01 AG015819, U01 AG072572, and U01 AG046152. MW is supported by Australian Research Council Discovery Project grant (DP160100011). JH is supported by R37 HL63762, R01 NS093382, RF1 AG053391, the Brightfocus Foundation and the Bluefield Project. DF is supported by JPco-fuND- European research For Neurodegenerative Disease and Israel Ministry Of Health CSO-MOH #30000-12631, Israel Ministry Of Science And Technology 3-12808. We wish to thank Yuval Nash from Tel Aviv University for analysis of *in situ* experiments. The pHLSec vector was a kind gift from Dr Radu Aricescu (University of Oxford). The single molecule work was supported by NIH grant MH054137 (J.A.J), the Hope for Depression Research Foundation (J.A.J.), and Miriam’s Magical Memorial Mission (J. A. J.). We thank Peter Geggier for discussions related to analysis of the smFRET imaging data and Scott C. Blanchard for providing the SNAP_f_-reactive Lumidyne fluorophores.

## AUTHOR CONTRIBUTIONS

RBD, ME, YZho, KS, YZha, FC contributed equally. RBD and PHStGH conceived the project. CMJ, MI, JH, MW, PEF, DAB, PLDJ, JJM, JAJ, WBA, EMB and PHStGH contributed to experimental design. Cloning, protein expression and purification were carried out by RBD and WM. Crystallisation, data collection and structure determination were performed by RBD. SEC-MALS was performed by CMJ. Binding experiments were performed by RBD, ME, YZha, KS, CB, JW, MAK, SS, SP, AAB, ES, PEF, MI, SQ, FC, XX, JH. Ex-vivo dimer coIP experiments were performed by CDR, FC, EMB, and PLDJ. Flow cytometry and floPLA were done by SJR, YZho, JLH, KAT. Human plasma CLU was purified by JW, MAK, SS and MRW. Human ApoE purification and CD33 binding was done by XX, JH. In vivo plaque clearance studies were done by AR, ZF, DF. Phosphorylation and signalling experiments were done by YZha, CB, YZho, JLH, KAT, EMB, AT, PEF, JS, MR. smFRET experiments were designed by WBA, JAJ, and JJM, and the expression plasmids and stable CHO cell lines were generated by WBA and GVD. JJM and GVD performed all live-cell smFRET imaging experiments, and JJM analyzed the data with input from JAJ and WBA. WBA supervised the single-molecule component of the project. JLH and KAT respectively performed the CD33-SHP1 PLA and MDMi Aβ uptake studies. AJL, BNV, and ML did the genetic, eQTL and mendelian causal analyses. RP and MJL generated the human iPSC-derived microglia and with YZho, JG and ESW performed the sCD33^M^ analyses. Source data acquisition, display, curation and sharing was done by BA, ASB and DG. All authors contributed to the writing of the manuscript. Current addresses: RBD: Biologics Engineering, SJR: Clinical Pharmacology and Safety Sciences at R&D, AstraZeneca, Cambridge, UK;.

## COMPETING INTERESTS

The authors have no competing interests to declare.

## SUPPLEMENTARY DATA

- Source images for western blots are at the end of this file.
- Numerical data is available in separate Excel spread sheets for main data and supplementary data.
- Exact P-values are separately provide in Supplementary Table.

**SUPPLEMENTARY FIGURE 1A.**
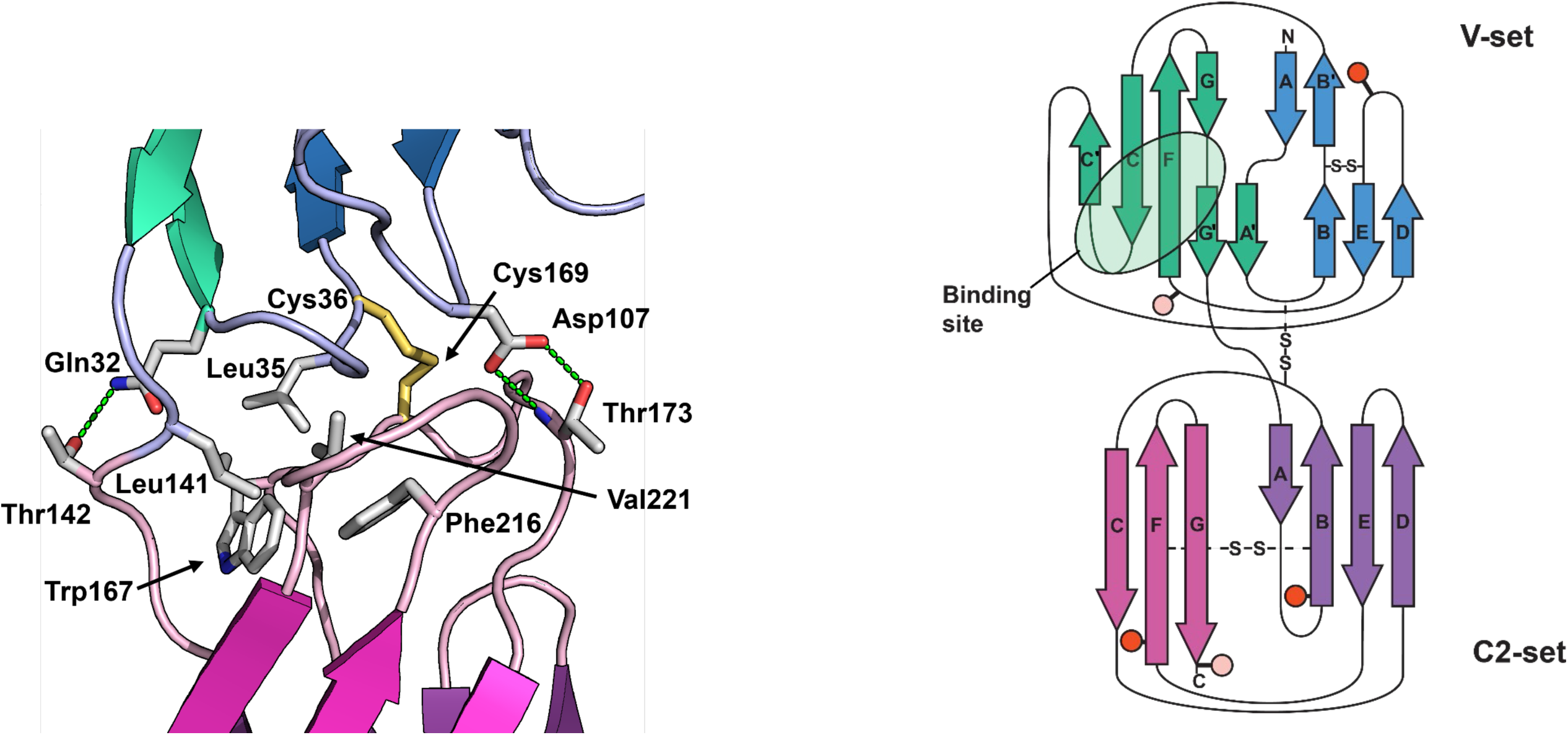
**LEFT**: Details of the interface between the V-set and C2-set domains. Indicated are the hydrophobic side chains in the centre of the interface and the polar side chains involved in hydrogen bonds (green dotted lines). The interdomain disulphide bond between Cys36 and Cys169 is shown in yellow. **RIGHT**: The topology of the V-set and C2-set domains, coloured as in (ii). The glycosylation sites are indicated by red circles (fully occupied) and pink circles (partial occupancy).

**SUPPLEMENTARY DATA FIGURE 1B.**
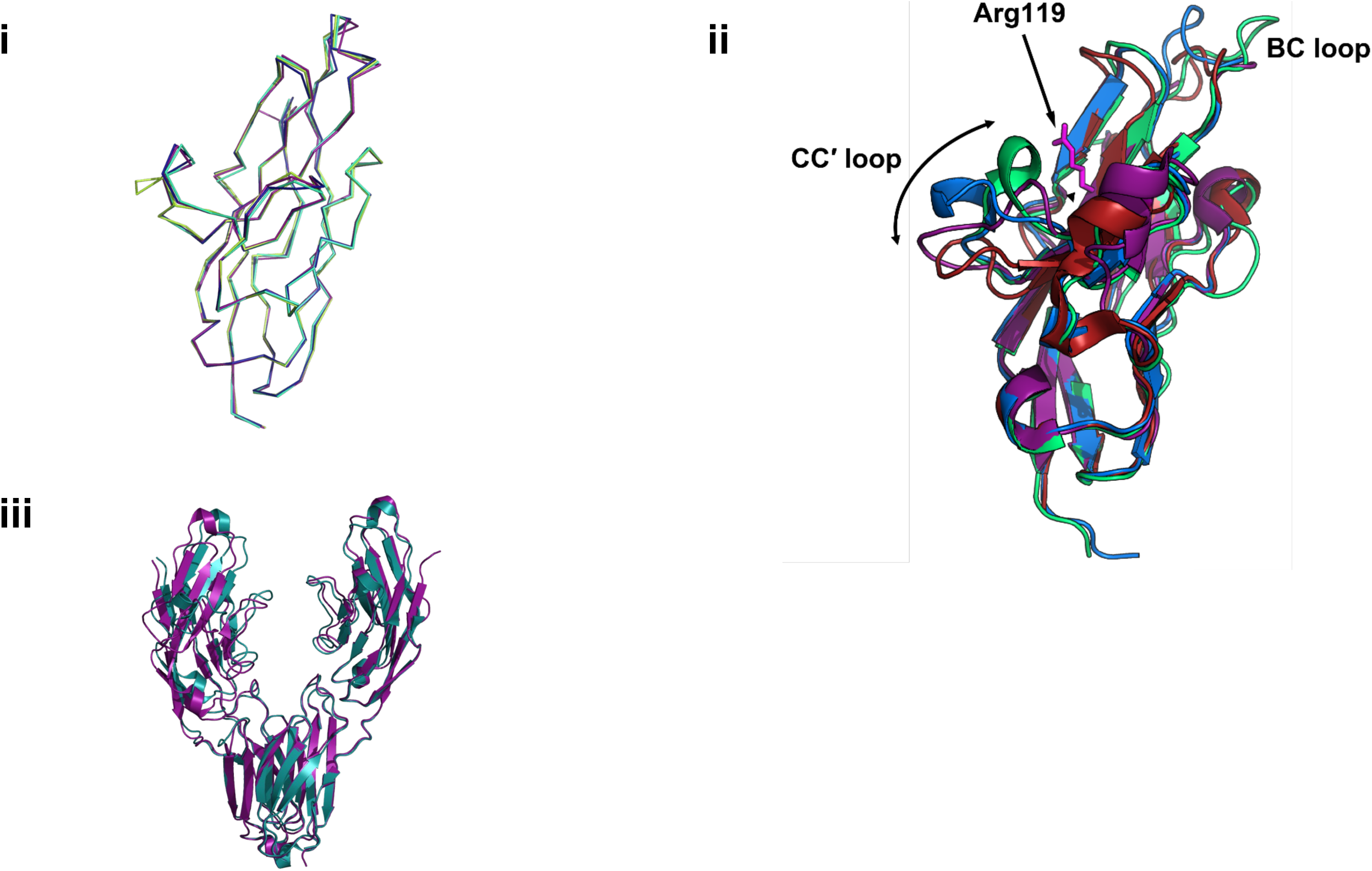
Structural comparisons to additional CD33 structures and paralogue siglec structures. **Structural alignments of i – four CD33 V-set domain structures: 5IHB (lime), 6D48 (purple), 7AW6 (blue) and AF_P20138_F1 (AlphaFold2 model, green); ii V-set domains of sialoadhesin (1OD9, red), CD33 (blue), Siglec-5 (2ZG2, green) and Siglec-7 (2HRL, purple).** Significant deviations are limited to the BC loop and the CC′ loop. The CC′ loop occupies conformations varying from more open (CD33^M^) to closed (Siglec-5) over the binding site; **iii full-length ECDs of 5IHB (cyan) and 7AW6 (purple) aligned on C2-set domain only. All structures were aligned using the align algorithm in PyMOL.**

**SUPPLEMENTARY DATA FIGURE 1C.**
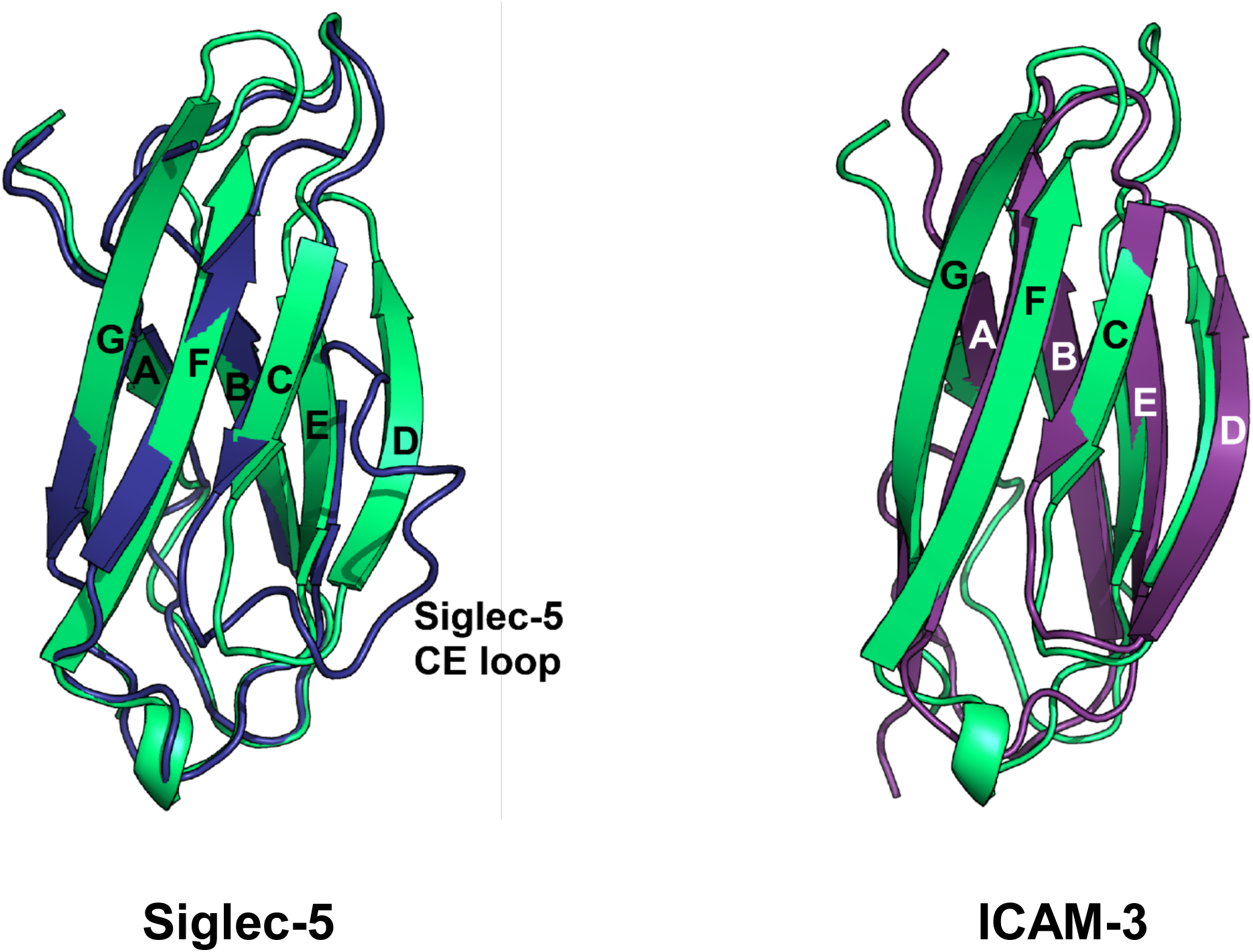
Structural alignments of CD33 C2-set domain. *Left panel -* Cartoon showing a structural alignment between the C2-set domains of CD33 (green) and Siglec-5 (blue), with an RMSD = 0.93 Å. Whereas the CD33 domain possesses a D strand, the region between the C and E strands in Siglec-5 occupies a non-regular loop conformation *Right panel:* Structural alignment of the C2-set domains of CD33 (green) and ICAM-3 (purple). Both domains possess a D strand. RMSD = 1.9 Å.

**SUPPLEMENTARY DATA FIGURE 1D.**
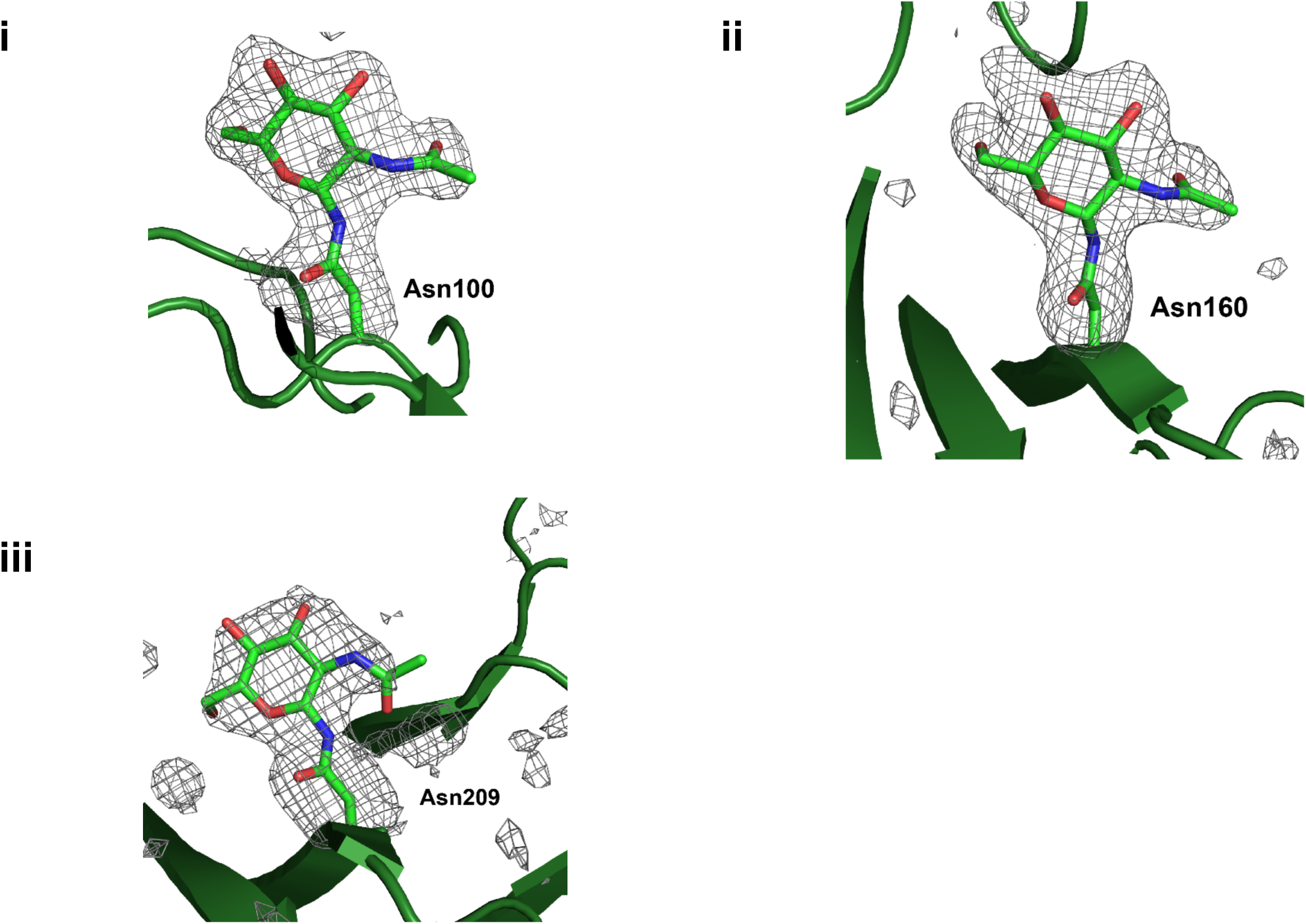
Details of the three *N*-linked GlcNAc residues built per chain. The protein chains at the three glycosylation sites, **i** – Asn100, **ii** – Asn160, **iii** – Asn209, are shown in cartoon form, with the Asn side chains and GlcNAc residues shown as sticks. Simulated annealing *mFo-DFc* omit maps (Asn and GlcNAc omitted) are displayed as a grey meshwork at 2.0 σ.

**SUPPLEMENTARY DATA FIGURE 1E.**
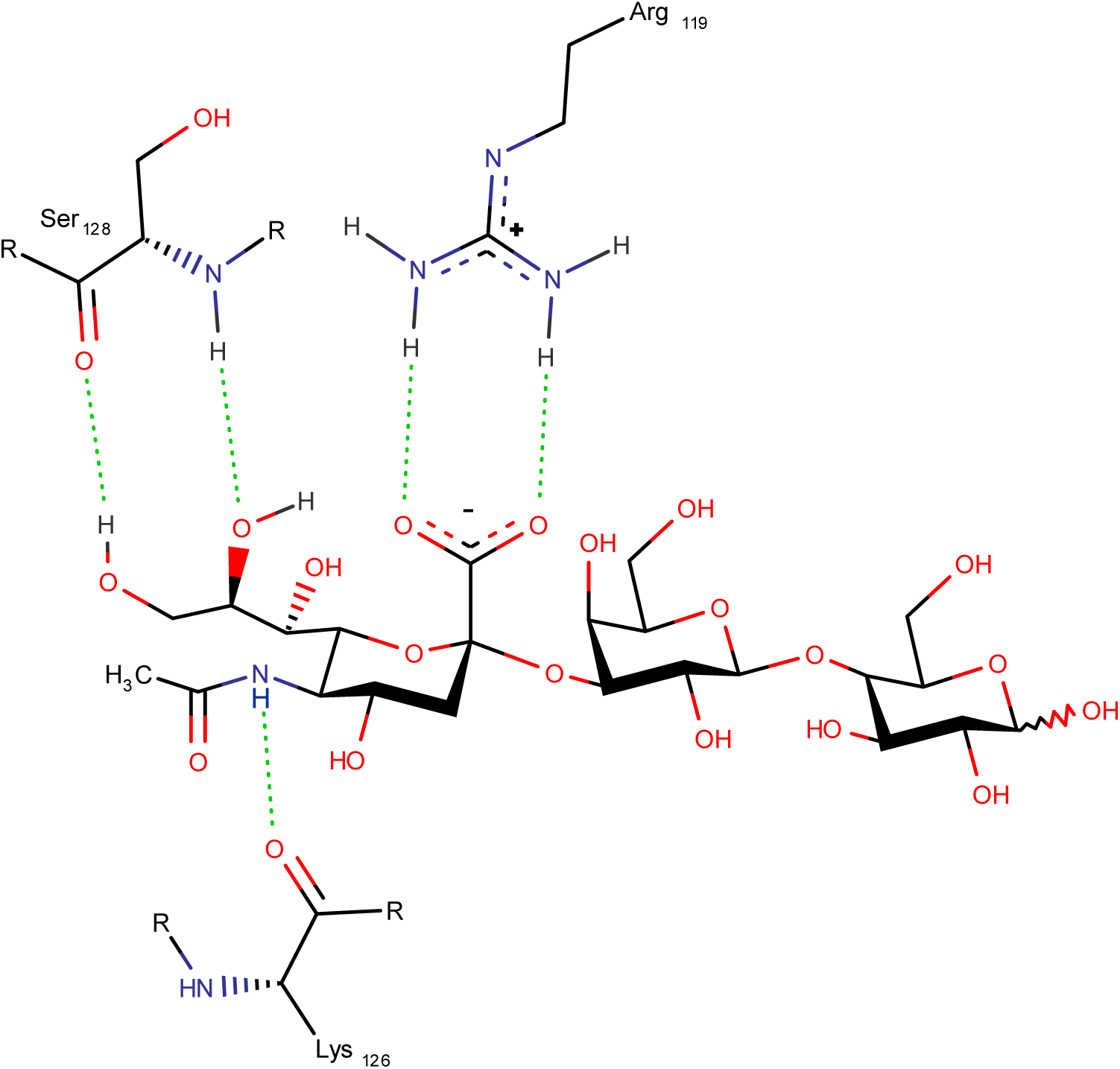
A schematic of the ligand binding site showing the interactions between CD33 and 3’-sialyllactose (identical interactions are seen for 6’-sialyllactose). No interactions are made with the galactose or glucose residues. Figure prepared using PoseView.

**SUPPLEMENTARY DATA FIGURE 1F.**
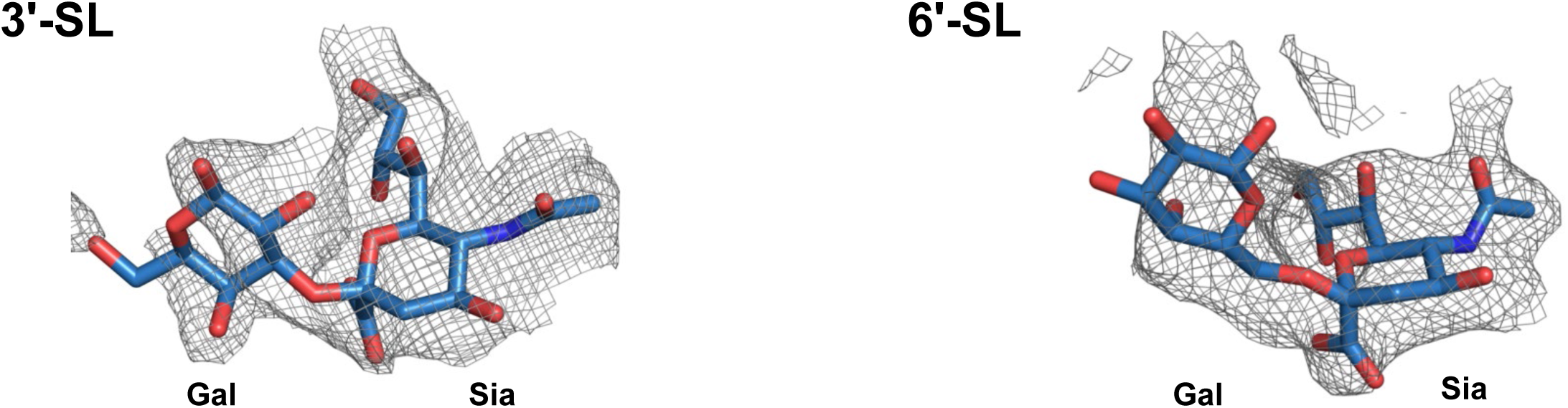
**Top panel:Electron density maps for 3’-sialyllactose and 6’-sialyllactose ligands** The two ligand molecules (each from their respective chain B binding site) are shown in stick format. Simulated annealing *mFo-DFc* omit maps (ligand molecules omitted) are displayed as a grey meshwork at 2.0 σ.

**SUPPLEMENTARY DATA FIGURE 1G.**
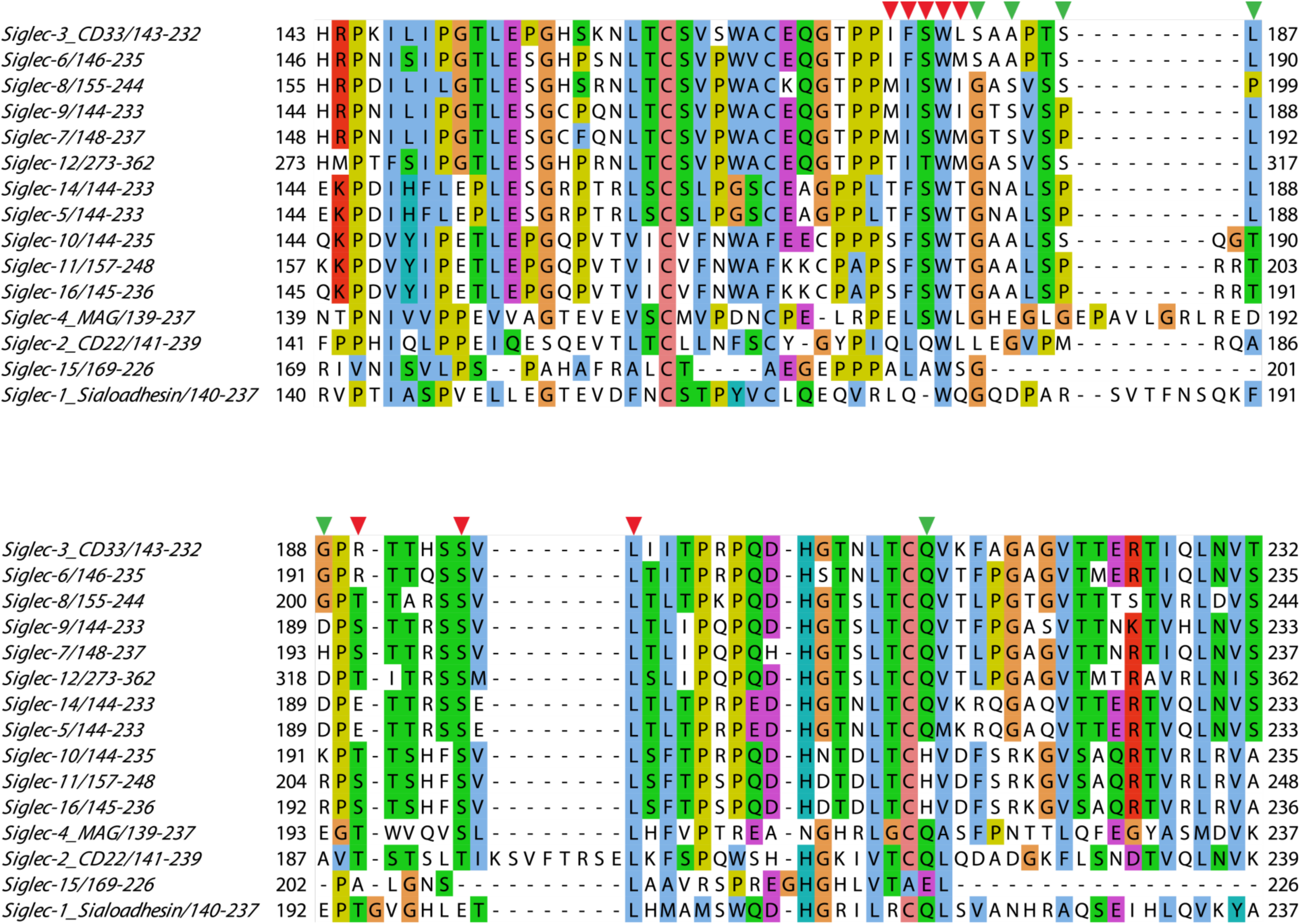
Sequence conservation of dimer interface residues in CD33 C2-set domains of multiple human Siglecs. This region was identified by the EPPIC program as likely to form biologically authentic dimer interfaces. Red arrows depict residues determined by EPPIC to have > 95% buried surface area in the interface, green arrows residues with > 70% buried surface area. These are highly conserved in a subset of the sequences.

**SUPPLEMENTARY DATA FIGURE 1H.**
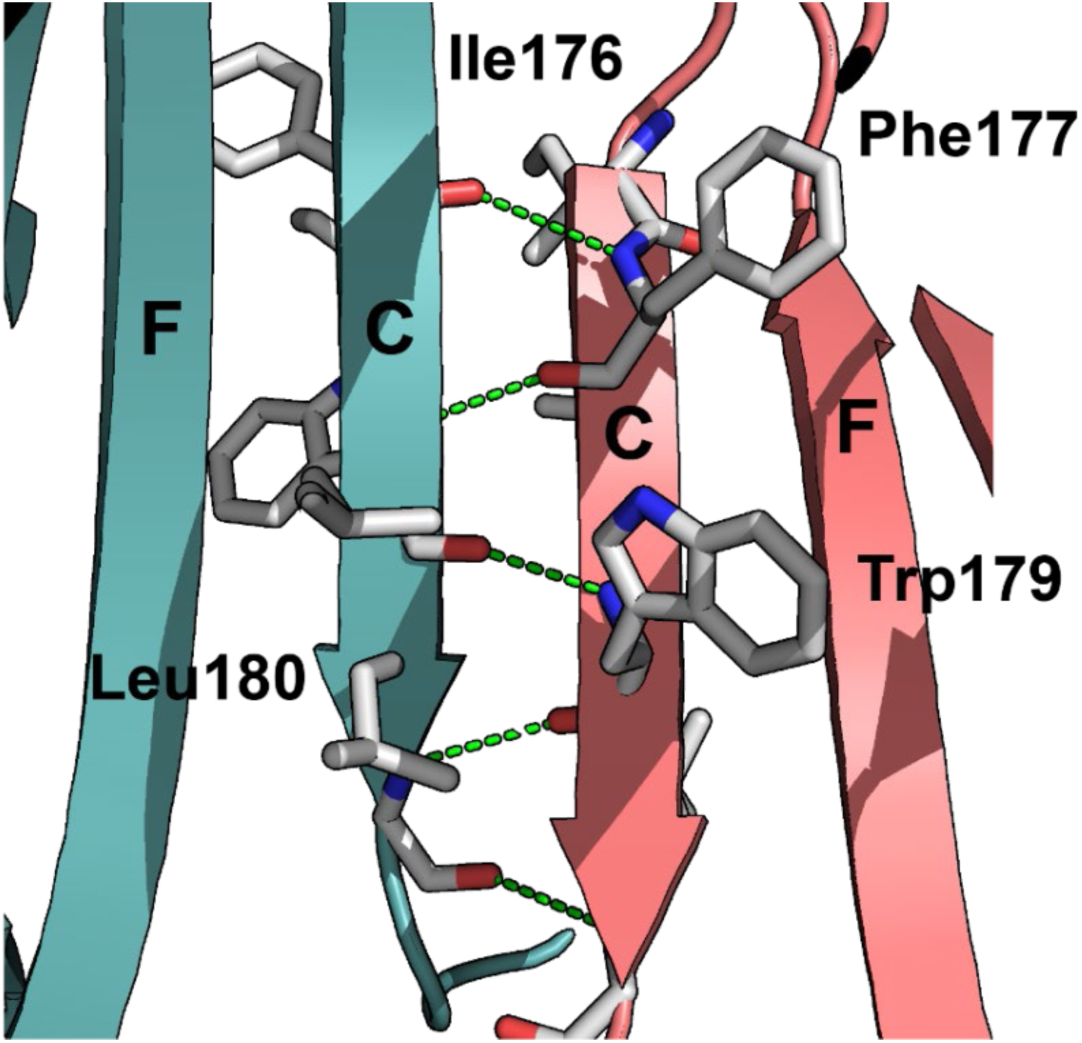
Detail of the β-strand interactions at the centre of the dimer interface, showing parallel arrangement of the two C-strands, with a series of main chain hydrogen bonds (green dotted lines), and with the side chains of several hydrophobic residues forming critical packing interactions (Leu176, Phe177, Trp179 and Leu180). Mutagenesis experiments show that all residues contribute to dimer formation.

**SUPPLEMENTARY DATA FIGURE 1I.**
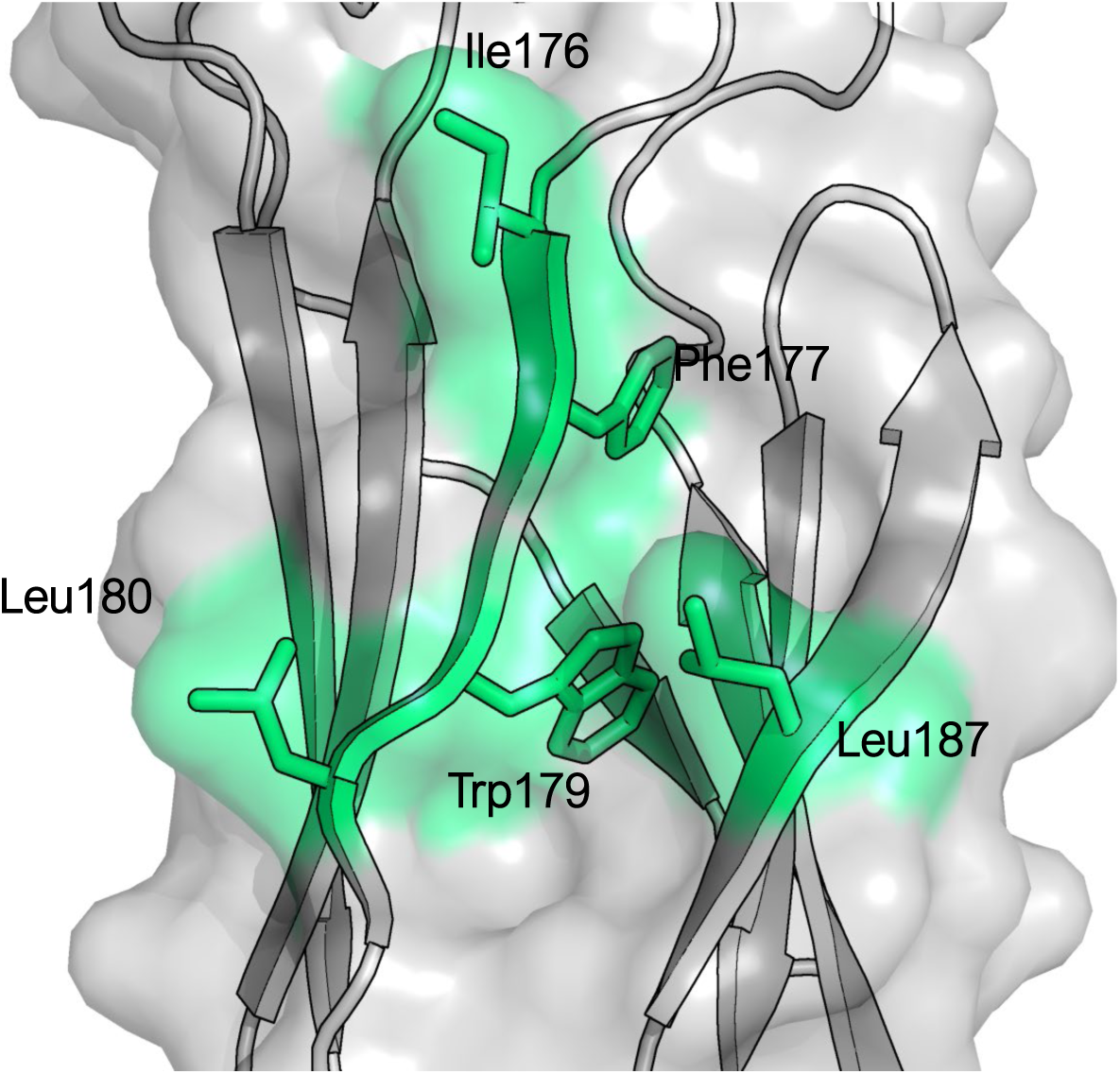
Detail of the hydrophobic residues interacting at the dimeric interface. The surface is coloured green for hydrophobic residues and grey for polar residues. The 5 hydrophobic side chains (Ile176, Phe177, Trp179, Leu180 and Leu187) are shown as sticks.

**SUPPLEMENTARY DATA FIGURE 1J.**
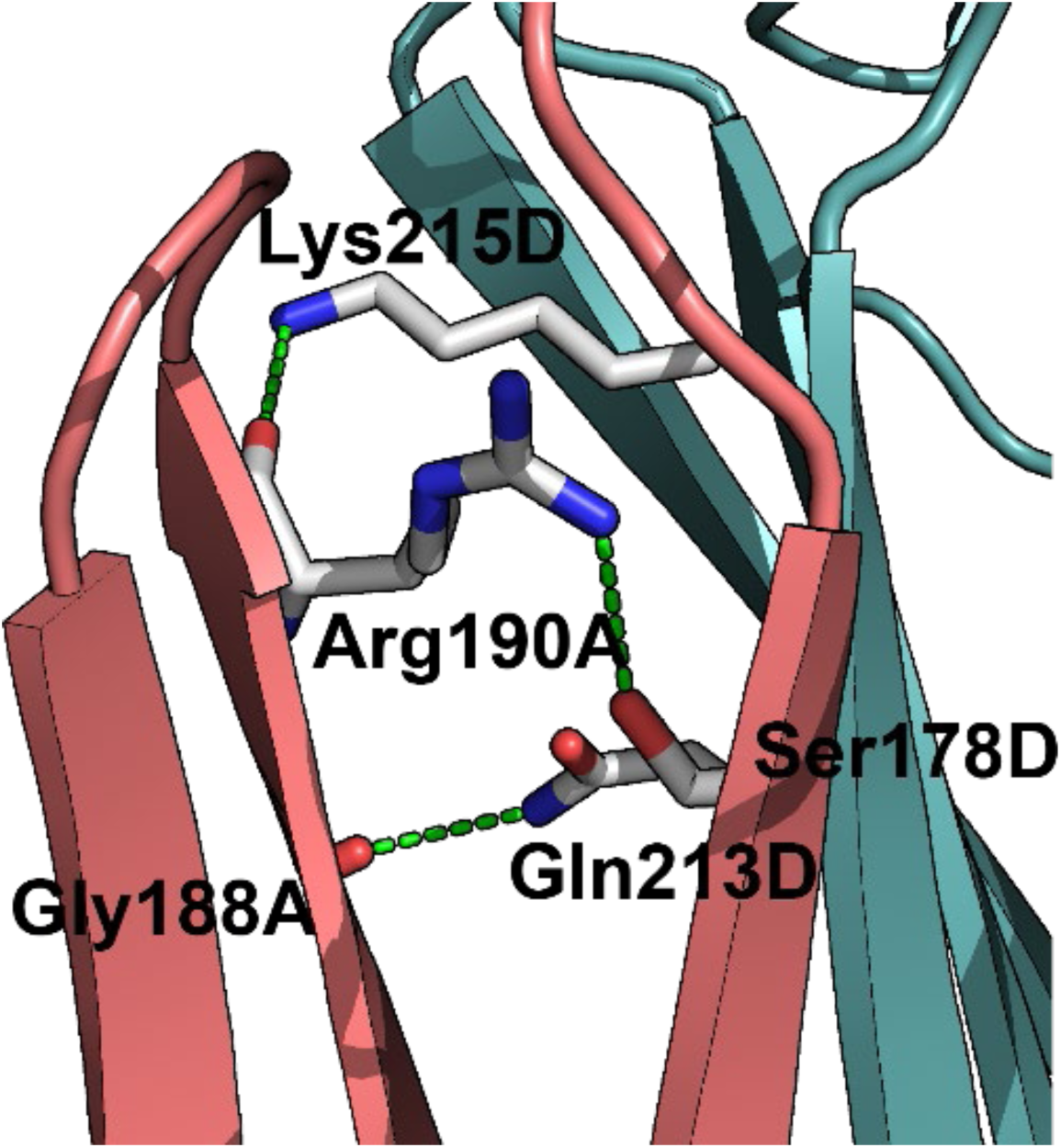
Detail of several hydrogen bonding interactions at the dimeric interface. Interacting residues are shown as sticks.

**SUPPLEMENTARY DATA FIGURE 2A.**
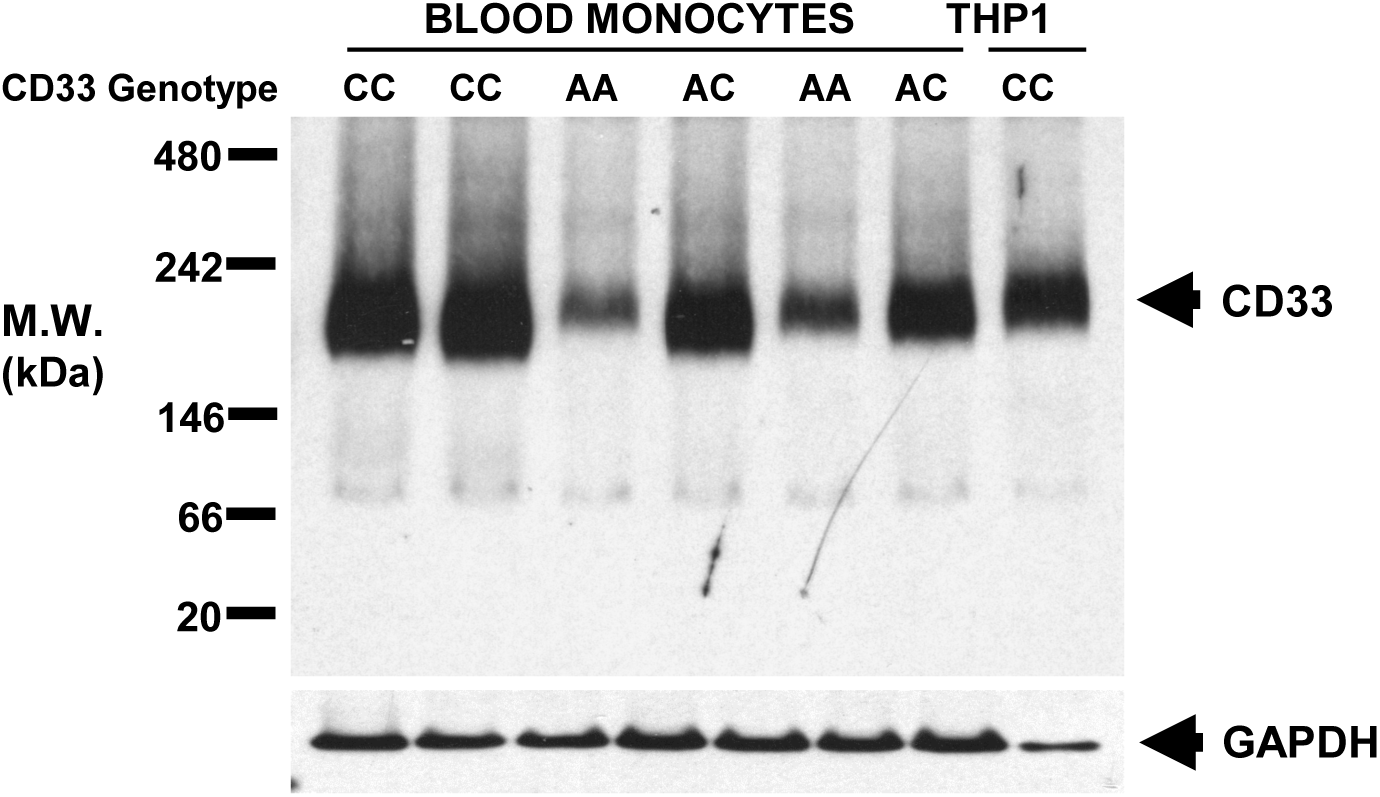
Biological confirmation of dimerization. Blue-Native PAGE analysis of digitonin-lysed native CD33 from fresh human peripheral blood monocytes and THP-1 cells. Western blotting with an anti-CD33 antibody demonstrates a species migrating in excess of the predicted monomeric size of 67- 75 kDa.

**SUPPLEMENTARY DATA FIGURE 2B.**
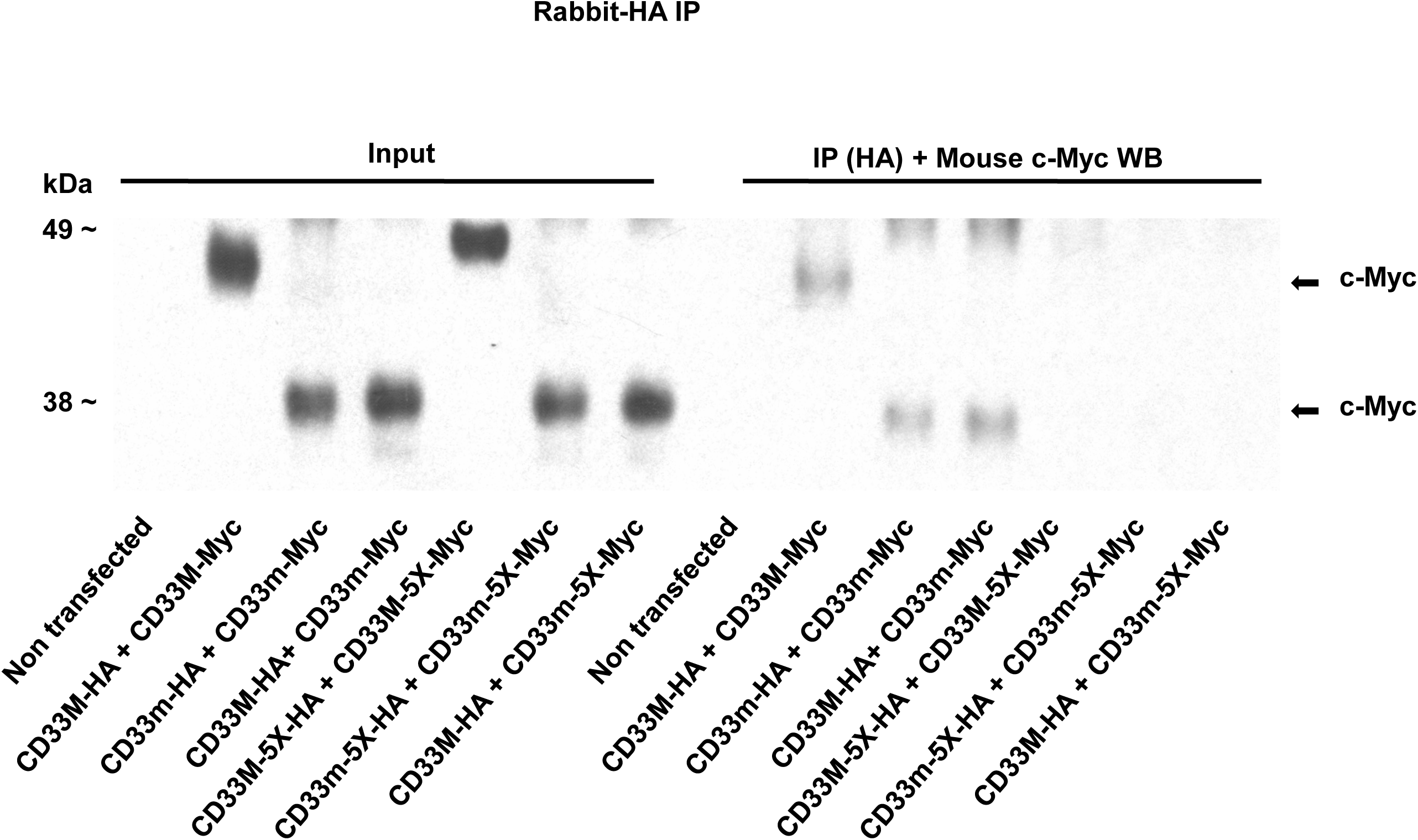
Mutagenesis of Phe177Ala + Trp179Ala + Leu180Ala + Arg190Ala + Gln213Ala residues within the dimer site was performed to disrupt the hydrophobic and polar interactions in the dimer site. Mutation of any combination of 1-4 of these 5 residues had no effect on dimer formation. However. mutation of all 5 residues resulted in formation of stable, fully glycosylated CD33^M^ and CD33^m^ monomers that do not co-precipitate other CD33 peptides

**SUPPLEMENTARY DATA FIGURE 2C.**
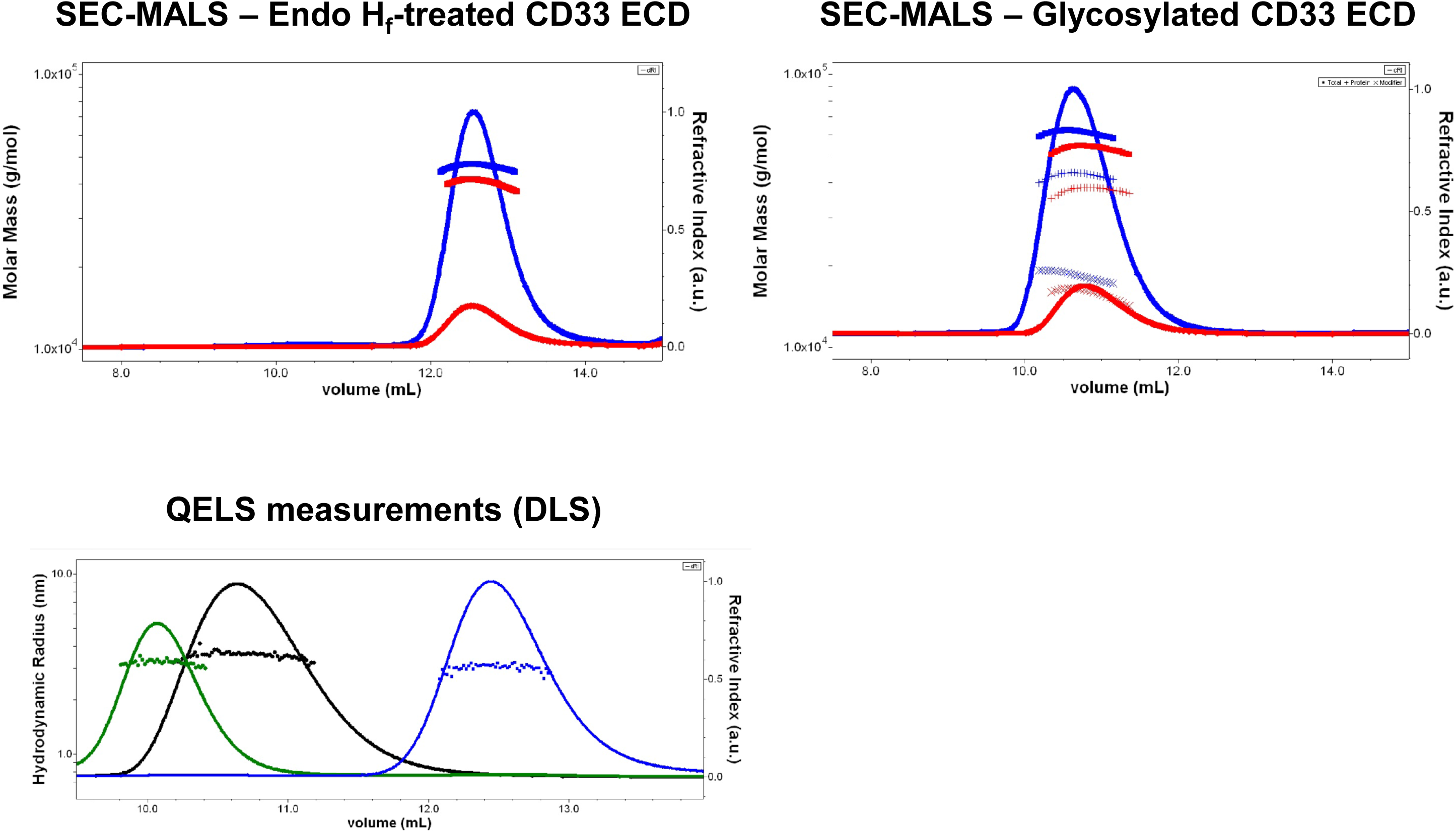
**SEC-MALS demonstrates CD33 ECD dimers in solution** ***Left Panel* - SEC-MALS analysis of Endo H_f_-trimmed CD33 ECD.** Shown are plots of refractive index (RI, right axis) and evaluated molar mass (left axis) against elution volume for samples loaded at 2.5 mg/ml (blue) and 0.5 mg/ml (red). The glycan-trimmed sample were considered as all protein but still analysed using either UV and RI as concentration signals. At the higher concentration (∼ 10 µM on column), RI indicates a mass of 47 kDa over the central 50 % of the peak while analysis using UV absorbance as the concentration source gives a mass of 50 kDa. These results confirm that CD33 ECD is present predominantly as a dimer in solution at the concentrations achieved over the central 50% of the SEC peak (∼ 5-10 µM). At the lower concentration loaded (∼ 2 µM on column) the apparent mass is slightly lower at 41 kDa and 44kDa (RI, UV). Consequently we can estimate that the Kd for this dimerisation is much below µM and probably on the order of 100s of nM. ***Right Panel* - SEC-MALS of fully glycosylated CD33 ECD.** As for the trimmed sample, protein was loaded at 2.5 mg/ml (blue) and 0.5 mg/ml (red). The glycosylated samples eluted overall earlier than the Endo H-trimmed samples consistent with a higher expected mass and larger hydrodynamic properties contributed by the additional carbohydrate. Due to the significant contribution of glycans to the mass of the system, a dual concentration conjugate analysis was applied (*dn/dc* protein 0.1885 ml g^-1^, *dn/dc* carbohydrate 0.145 ml g^-1^, UV protein 1252 ml g^-1^ cm^-1^, UV carbohydrate no absorbance). This analysis evaluates a protein mass of about 43 kDa (symbol +) and a carbohydrate mass of about 18 kDa (symbol X) consistent with dimer formation and with the predicted mass of the carbohydrate modifications on CD33 giving a total mass around 60 kDa (symbol filled square). ***Bottom Panel* - QELS measurements (DLS)** made for the SEC-MALS samples injected at 2.5 mg/ml. The hydrodynamic radius calculated for glycosylated CD33 ECD (black) of 3.5 nm is larger than expected for a 50kDa dimeric protein (for example, larger than that of the BSA standard (green) which is a 66 kDa protein) consistent with the presence of the extensive carbohydrate modifications GlcNAc_2_Man_9_ at 5 positions. Endo H_f_-trimmed CD33 ECD (blue) has an Rh of 3.0 nm, which is more consistent with a simple globular protein of 50 kDa mass as expected following the almost complete removal of its carbohydrate adducts.

**SUPPLEMENTARY DATA FIGURE 2D.**
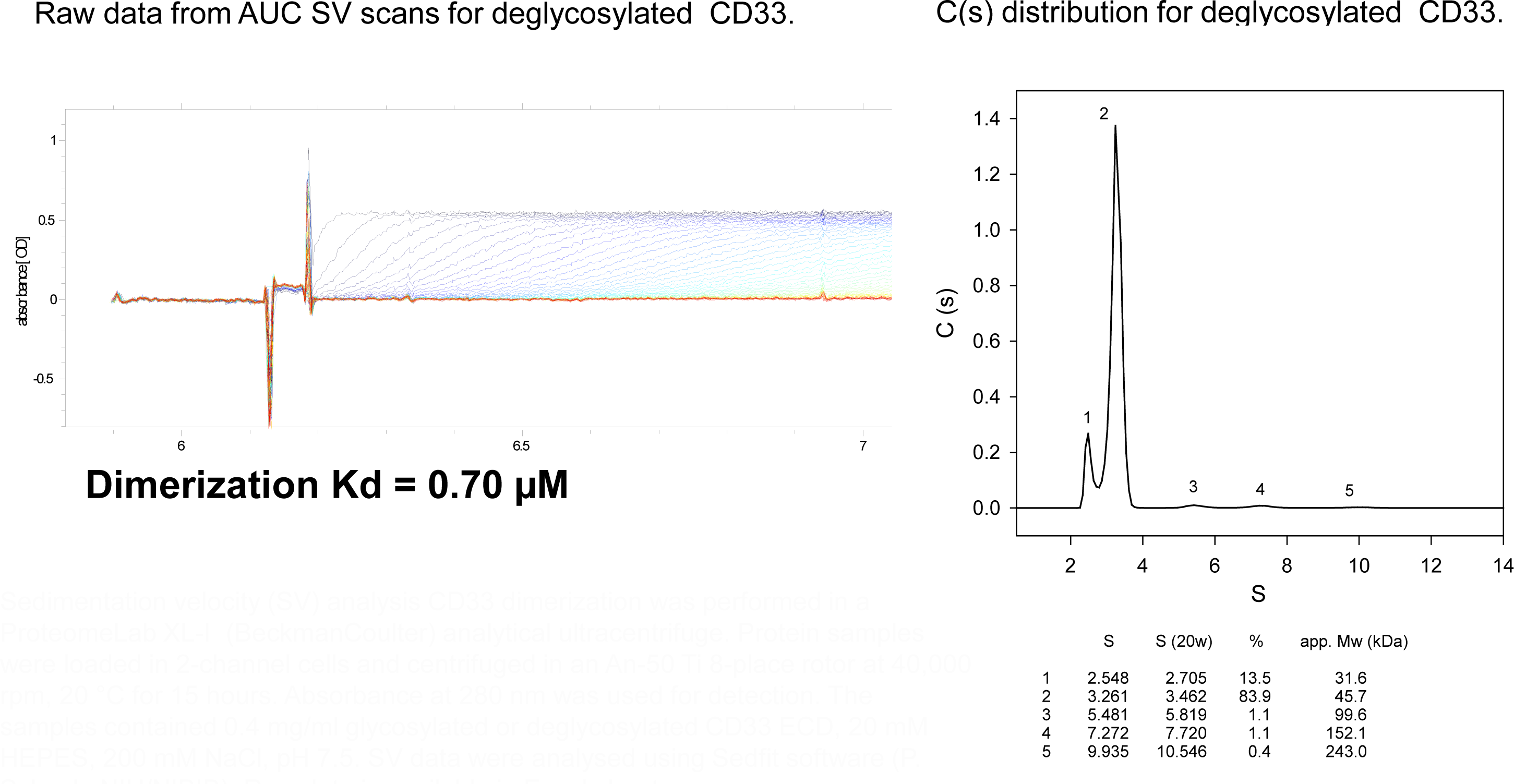
Analytical Ultracentrifugation (AUC) Sedimentation Velocity (SV) Experiment. Sedimentation velocity (SV) analysis CD33 dimerization was performed in a ProteomeLab XL-I (BeckmanCoulter) analytical ultracentrifuge. Protein samples were loaded in 2-channel cells and centrifuged in an An-50 Ti 8-place rotor at 40,000 rpm, 20 °C for 15 hours. Absorbance at 280 nm was used for detection. The samples contained 0.4 mg/ml glycosylated or deglycosylated CD33 ECD, 20 mM HEPES, 200 mM NaCl, pH 7.5. SV data were analysed using Sedfit software (P. Schuck, NIH/NIBIB). Raw data is available in Excel sheets

**SUPPLEMENTARY DATA FIGURE 2E.**
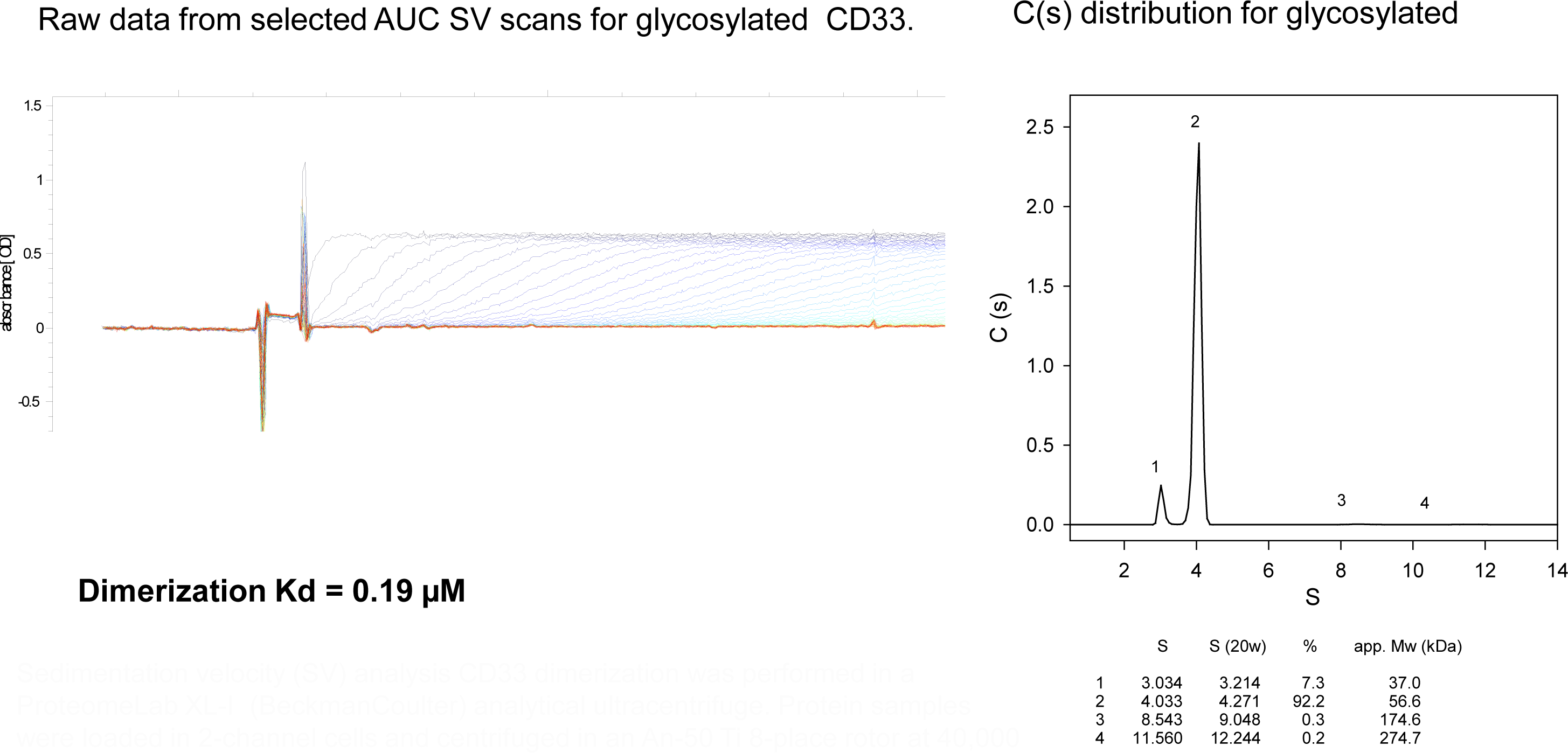
Analytical Ultracentrifugation (AUC) Sedimentation Velocity (SV) Experiment. Sedimentation velocity (SV) analysis CD33 dimerization was performed in a ProteomeLab XL-I (BeckmanCoulter) analytical ultracentrifuge. Protein samples were loaded in 2-channel cells and centrifuged in an An-50 Ti 8-place rotor at 40,000 rpm, 20 °C for 15 hours. Absorbance at 280 nm was used for detection. The samples contained 0.4 mg/ml glycosylated or deglycosylated CD33 ECD, 20 mM HEPES, 200 mM NaCl, pH 7.5. SV data were analysed using Sedfit software (P. Schuck, NIH/NIBIB). Raw data is available in Excel sheets

**SUPPLEMENTARY DATA FIGURE 2F.**
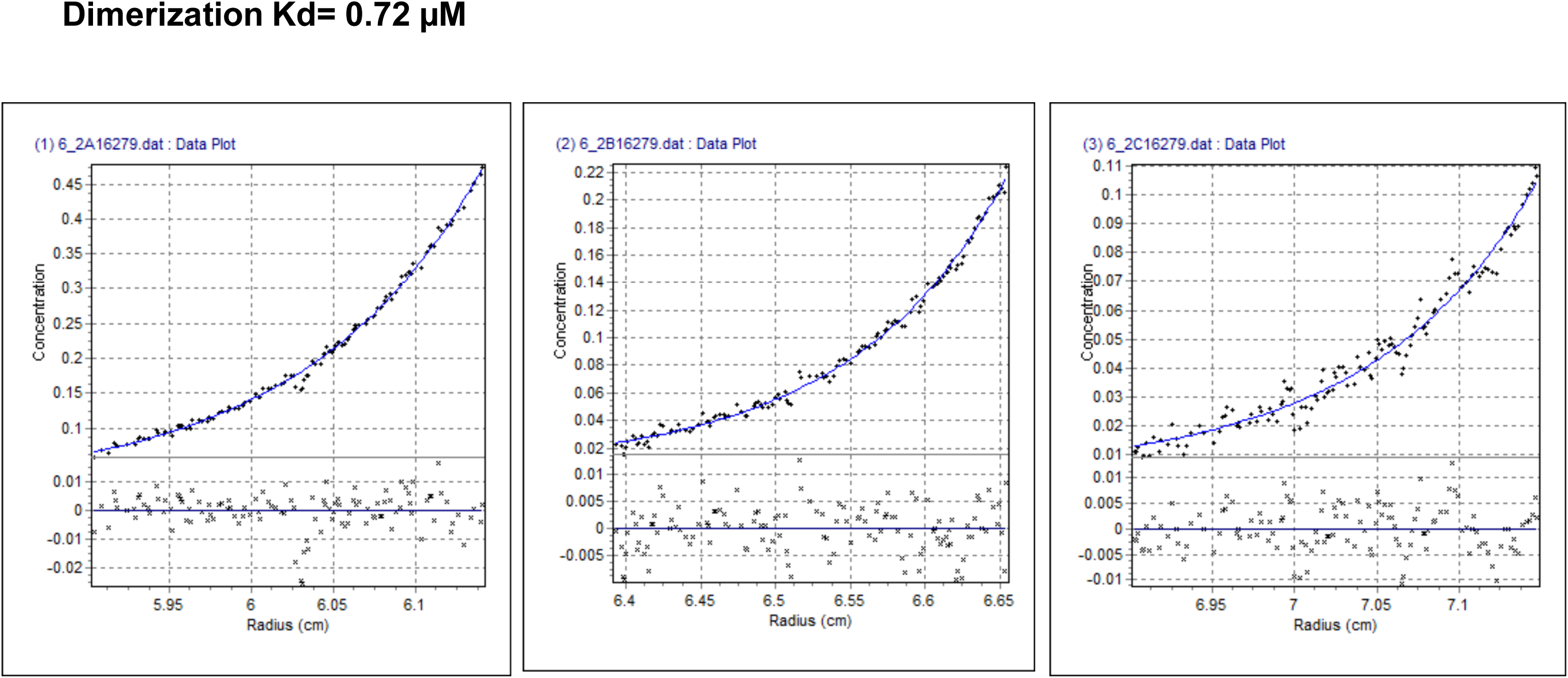
Sedimentation equilibrium (SE) experiments to analyse CD33 dimerization were performed in a ProteomeLab XL-I (Beckman Coulter) analytical ultracentrifuge. Samples of glycosylated and deglycosylated CD33 ECD at concentrations 0.2, 0.1 and 0.05 mg/ml were loaded in 6-channel equilibrium cells and centrifuged in an An-50 Ti 8-place rotor at 16,000 or 20,000 rpm and 20 °C until equilibrium was reached for each speed. Sample buffer was 20 mM HEPES, 200 mM NaCl, pH 7.5. SE data were analysed using HeteroAnalysis software (by J.L. Cole and J.W. Lary, University of Connecticut). **Analytical Ultracentrifugation (AUC) Sedimentation Equilibrium (SE) Experiment** AUC SE raw data (black) from 16,000 rpm run are shown for 0.2, 0.1 and 0.05 mg/ml deglycosylated CD33 (from left to right). Global fit (blue) was used to determine dimerization Kd assuming Mw=25.0 kDa for monomer.

**SUPPLEMENTARY DATA FIGURE 2G.**
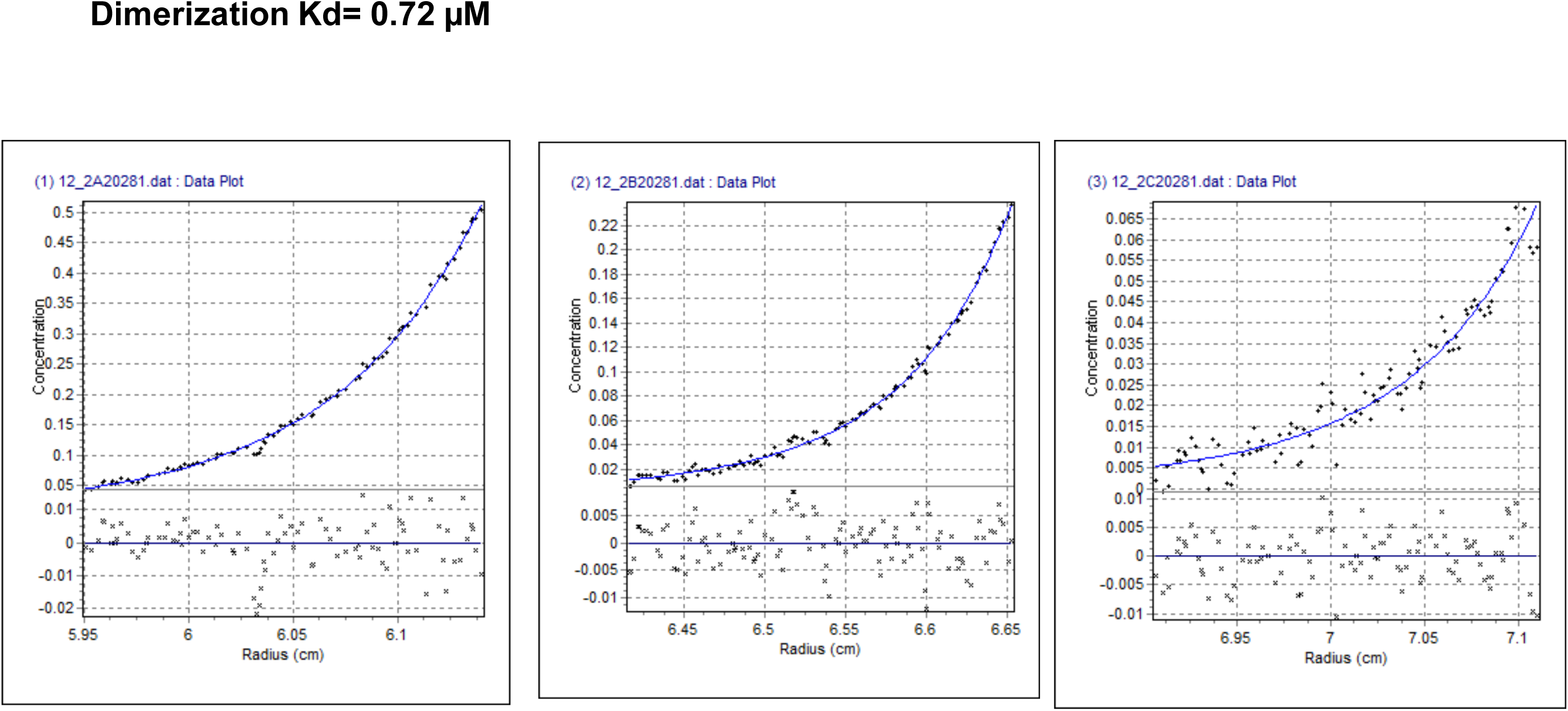
Sedimentation equilibrium (SE) experiments to analyse CD33 dimerization were performed in a ProteomeLab XL-I (Beckman Coulter) analytical ultracentrifuge. Samples of glycosylated and deglycosylated CD33 ECD at concentrations 0.2, 0.1 and 0.05 mg/ml were loaded in 6-channel equilibrium cells and centrifuged in an An-50 Ti 8-place rotor at 16,000 or 20,000 rpm and 20 °C until equilibrium was reached for each speed. Sample buffer was 20 mM HEPES, 200 mM NaCl, pH 7.5. SE data were analysed using HeteroAnalysis software (by J.L. Cole and J.W. Lary, University of Connecticut). **Analytical Ultracentrifugation (AUC) Sedimentation Equilibrium (SE) Experiment** AUC SE raw data (black) from 20,000 rpm run are shown for 0.2, 0.1 and 0.05 mg/ml deglycosylated CD33 (from left to right). Global fit (blue) was used to determine dimerization Kd assuming Mw=25.0 kDa for monomer.

**SUPPLEMENTARY DATA FIGURE 2H.**
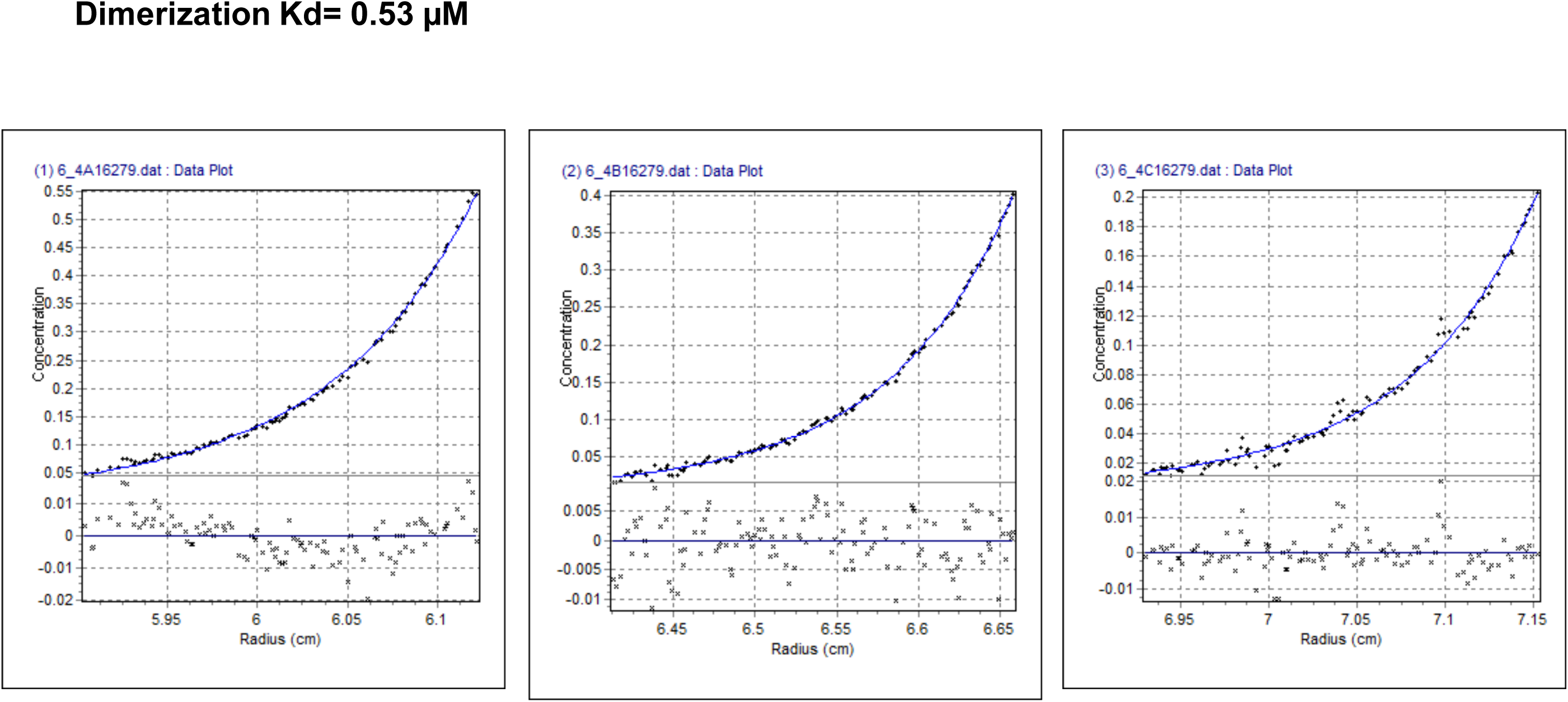
Sedimentation equilibrium (SE) experiments to analyse CD33 dimerization were performed in a ProteomeLab XL-I (Beckman Coulter) analytical ultracentrifuge. Samples of glycosylated and deglycosylated CD33 ECD at concentrations 0.2, 0.1 and 0.05 mg/ml were loaded in 6-channel equilibrium cells and centrifuged in an An-50 Ti 8-place rotor at 16,000 or 20,000 rpm and 20 °C until equilibrium was reached for each speed. Sample buffer was 20 mM HEPES, 200 mM NaCl, pH 7.5. SE data were analysed using HeteroAnalysis software (by J.L. Cole and J.W. Lary, University of Connecticut). **Analytical Ultracentrifugation (AUC) Sedimentation Equilibrium (SE) Experiment** AUC SE raw data (black) from 16,000 rpm run are shown for 0.2, 0.1 and 0.05 mg/ml glycosylated CD33 (from left to right). Global fit (blue) was used to determine dimerization Kd assuming Mw=31.3 kDa for monomer.

**SUPPLEMENTARY DATA FIGURE 2I.**
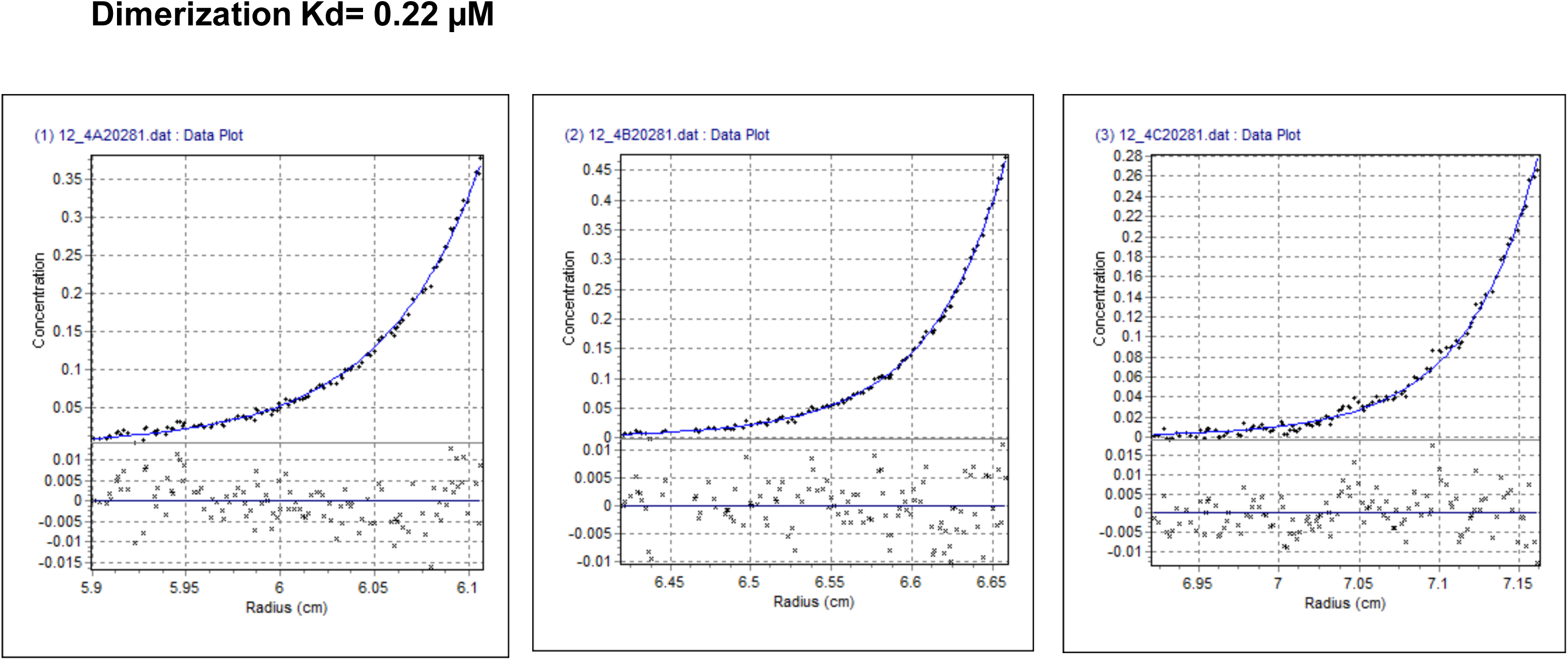
Sedimentation equilibrium (SE) experiments to analyse CD33 dimerization were performed in a ProteomeLab XL-I (Beckman Coulter) analytical ultracentrifuge. Samples of glycosylated and deglycosylated CD33 ECD at concentrations 0.2, 0.1 and 0.05 mg/ml were loaded in 6-channel equilibrium cells and centrifuged in an An-50 Ti 8-place rotor at 16,000 or 20,000 rpm and 20 °C until equilibrium was reached for each speed. Sample buffer was 20 mM HEPES, 200 mM NaCl, pH 7.5. SE data were analysed using HeteroAnalysis software (by J.L. Cole and J.W. Lary, University of Connecticut). **Analytical Ultracentrifugation (AUC) Sedimentation Equilibrium (SE) Experiment** AUC SE raw data (black) from 20,000 rpm run are shown for 0.2, 0.1 and 0.05 mg/ml glycosylated CD33 (from left to right). Global fit (blue) was used to determine dimerization Kd assuming Mw=31.3 kDa for monomer.

**SUPPLEMENTARY DATA FIGURE 2J.**
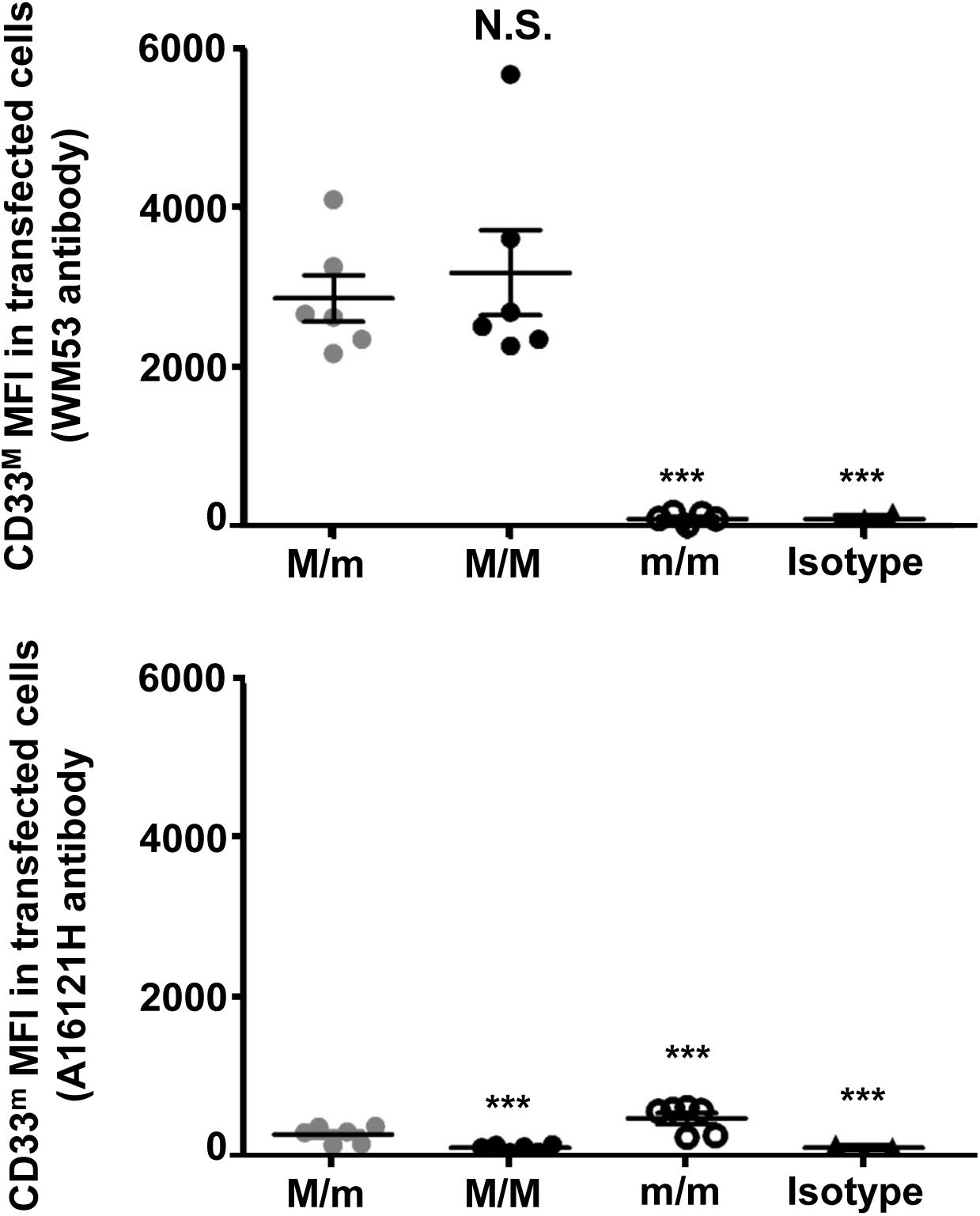
Flow cytometry analysis by double staining on CD33^M^/CD33^M^, CD33^m^/CD33^m^ or CD33^M^/CD33^m^ transfected cells using antibodies specific to CD33^M^ (WM53) and CD33^m^ (A16121H) respectively. Mean Fluorescence Intensities (MFIs) of cells expressing CD33^M^ (upper) and CD33^m^ (lower) were quantified. Error bars represent SEM. Statistical analysis was performed using one-way ANOVA followed by Tukey’s test (n = 6, in 3 independent experiments, \*\*\**p*.<0.001 *vs* M/m).

**SUPPLEMENTARY DATA FIGURE 2K.**
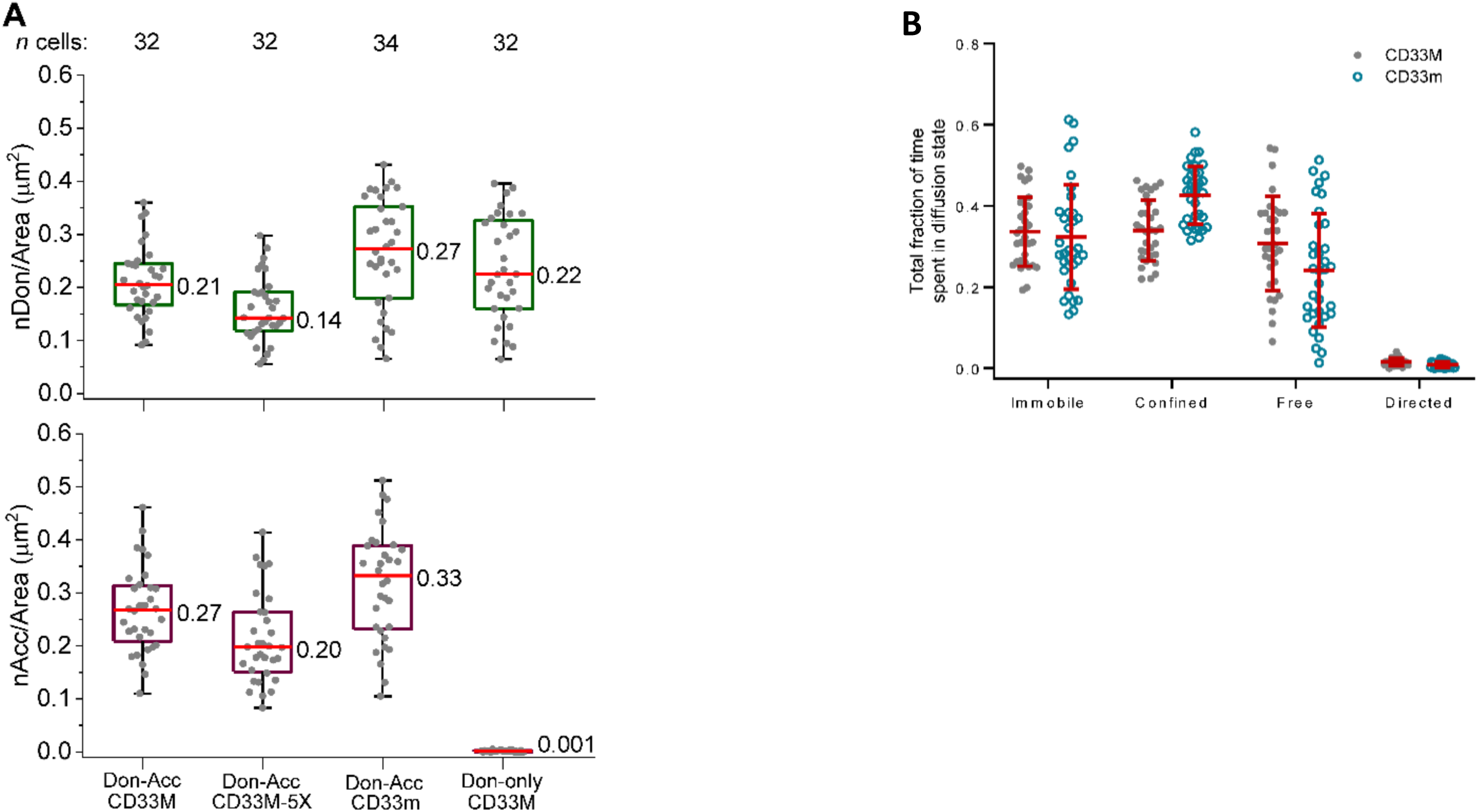
**Extended Data Figure 1K: Quantification of the surface density and diffusion states of labeled CD33 receptor samples.** **<u>A</u>,** Surface densities prior to smFRET imaging of donor and acceptor labeled samples for smFRET studies. Dots represent the number of acceptor (nAcc) or donor (nDon) particles per area for the indicated number of single cells (*n* cells) collected over 4 independent experiments for the CD33M and CD33m samples, and 3 independent experiments for the CD33M-5X and CD33M donor only samples. Box plot details are described in the legend of Fig. 2f-g. The median density of total labeled (acceptor + donor) receptors ranged from 0.34 – 0.60 molecules/μm^2^. The median density of the donor only labeled CD33M sample was 0.22 molecules/μm^2^ and near zero acceptors detected as expected.**<u>B</u>**, Total fraction of time spent in the immobile, confined, free, and directed diffusion states assigned by DC-MSS. Dots represent individual cell means and the middle and upper/lower lines depict the overall mean (values shown) and standard deviation, respectively, for 32 cells for the CD33M sample and 34 cells for the CD33m sample collected over 4 independent experiments.

**SUPPLEMENTARY DATA FIGURE 2L.**
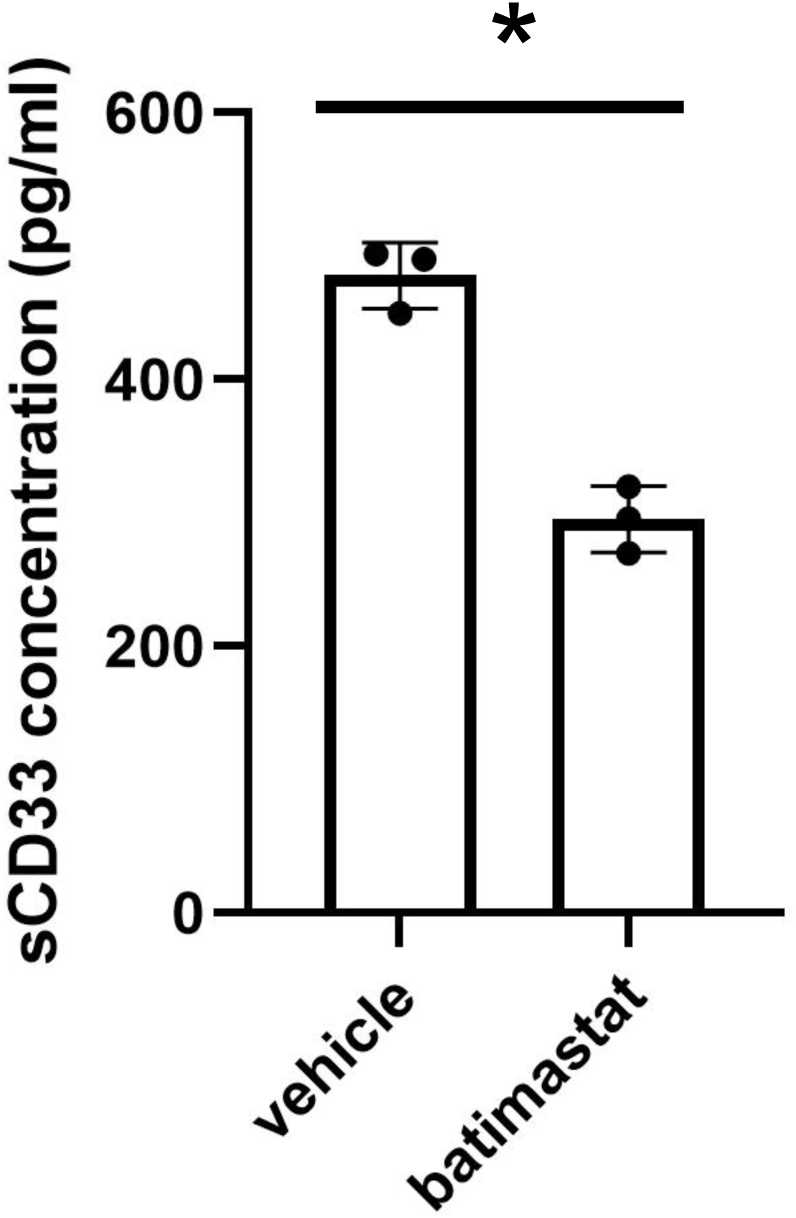
**A broad-spectrum sheddase inhibitor, batimastat could reduce soluble CD33 in iMG conditioned medium**. Differentiated iMGs were treated with 5 µM batimastat for 24 hours. Conditioned media was collected, centrifuged at 14,000 rpm for 10 minutes at 4°C. sCD33 were measured by ELISA (Abcam, ab283542) following the manufacturer’s protocol. Three independent wells of each treatments were collected, and all samples were assayed in duplicates. Statistical analysis was performed using paired t-test, p=0.0134.

**SUPPLEMENTARY DATA FIGURE 2M.**
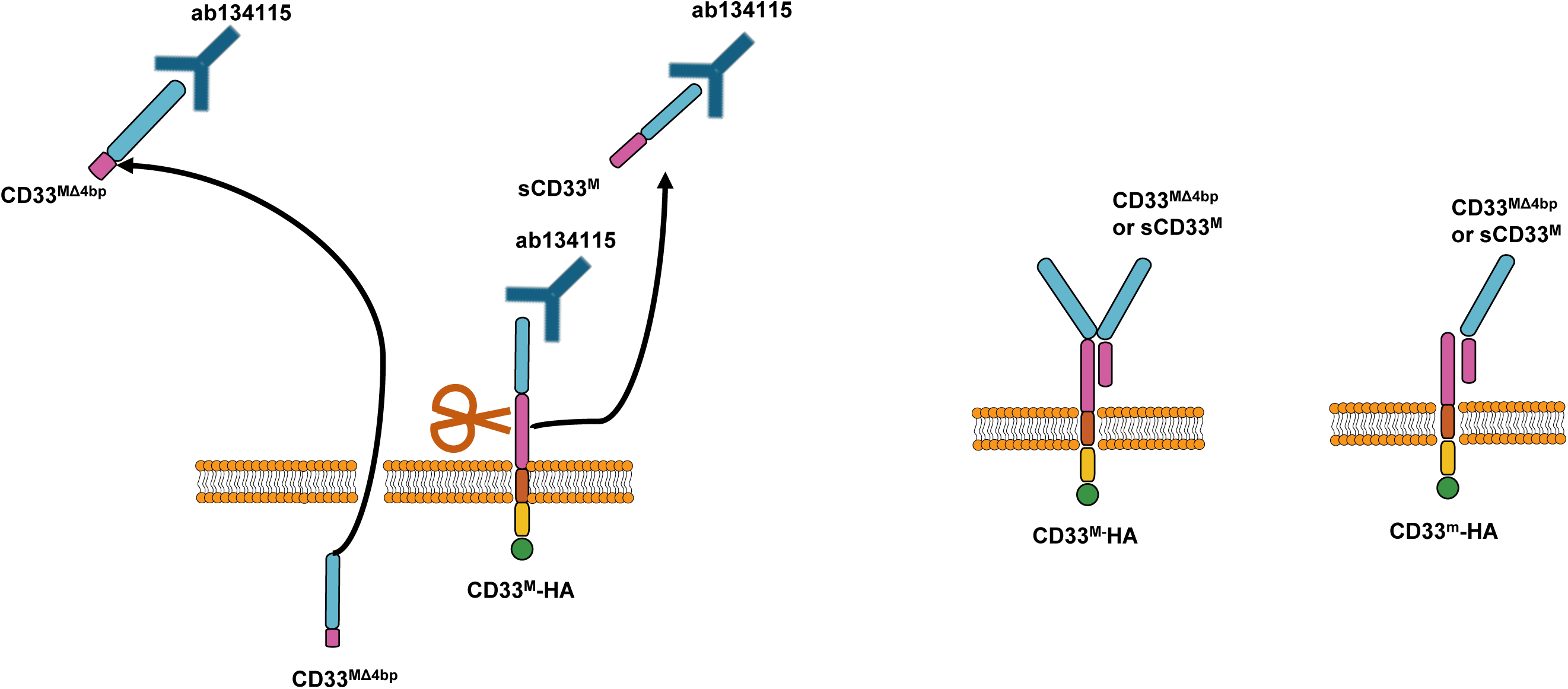
Cartoons depicting: **Left**: the putative secretion of CD33^MΔ4BP^ and putative endoproteolytic shedding of sCD33^M^, and the tagging and western blot strategy used here. The scissor cartoon depicts the site of a putative endoproteolytic cleavage or splice site that generates the endogenous soluble CD33^M^ ectodomain. **Right**: the putative binding of CD33^MΔ4BP^ (or speculatively sCD33^M^) to the extracellular domains of CD33^M^ or CD33^m^. The CD33^MΔ4BP^:CD33^M^ binding site is not defined., but could potentially recreate a dimeric ligand pocket with CD33^M^ monomers, and a monomeric ligand pocket with CD33^m^.

**SUPPLEMENTARY DATA FIGURE 3A.**
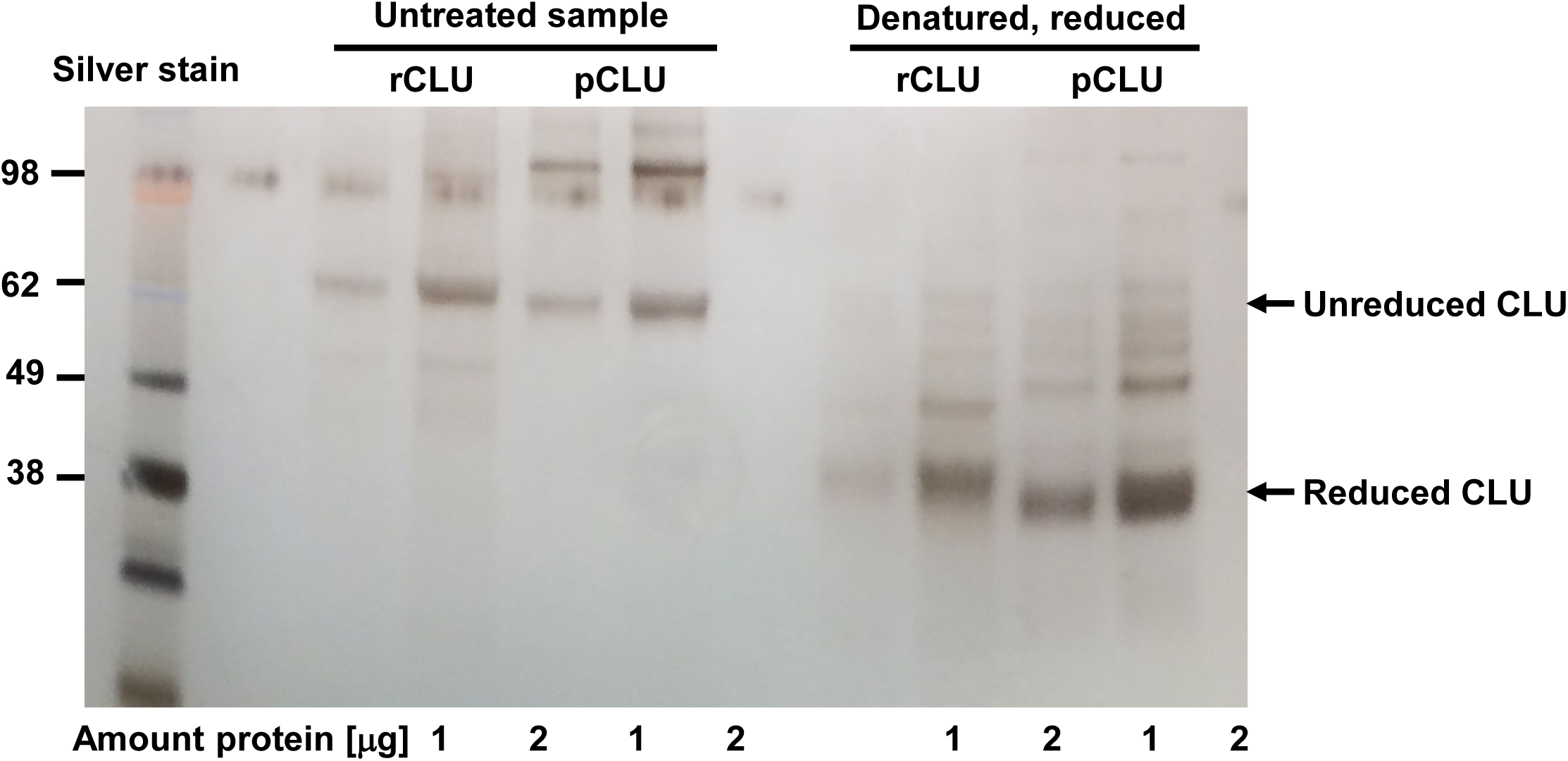
**Unprocessed Silver stained gel of recombinant clusterin (rCLU) or purified from human plasma (pCLU)** in the native state (left panel) and following denaturation and reduction (right panel).

**SUPPLEMENTARY DATA FIGURE 3B.**
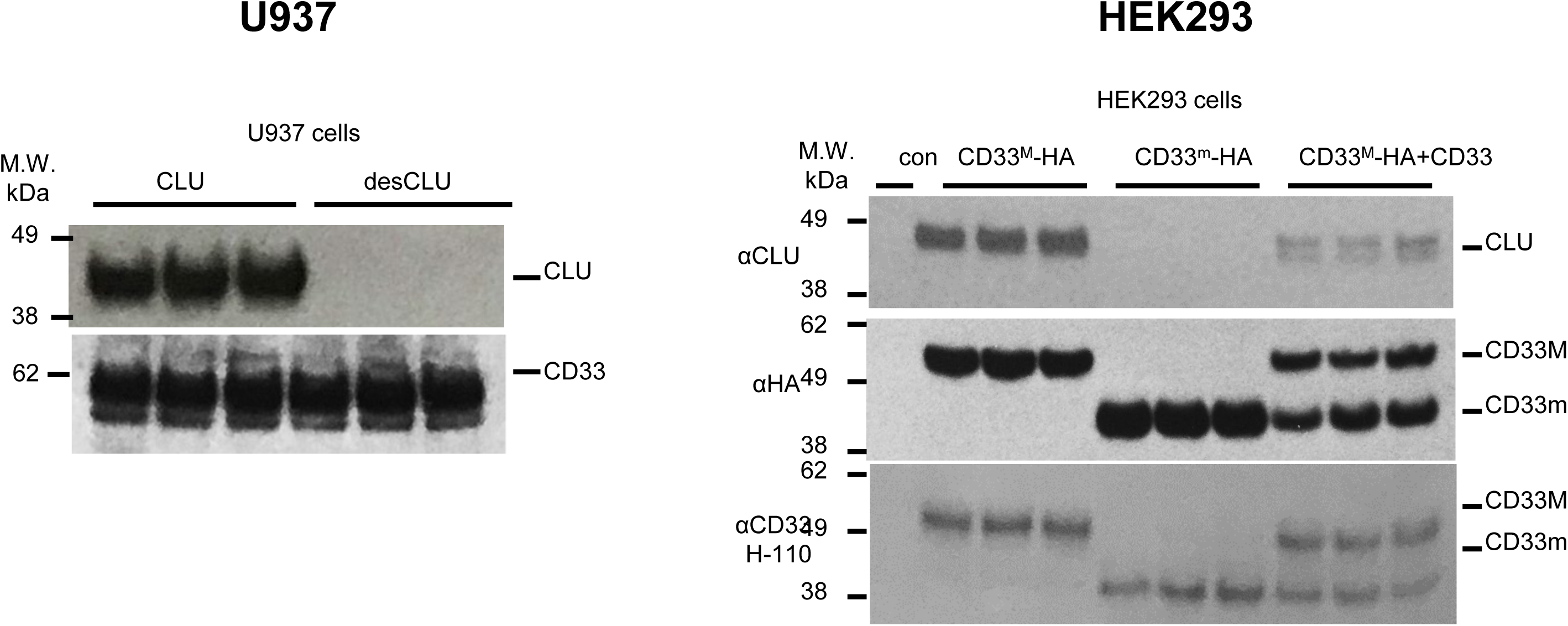
***Left*: endogenous CD33^M^ co-immunoprecipitates sialylated CLU, but not desialylated CLU in U937 cells.** Representative western blot for Figure 3a. *Top blot*: representative western blots of anti-CD33 co-immunoprecipitation products, blotted with anti-CLU antibodies. ***Lower blot*:** anti-CD33 co-immunoprecipitation products probed with anti-CD33 antibody, showing equivalent pulldown. This data is quantified in Figure 3a, left panel. Each lane shown represents one independent biological replication of the respective condition. ***Right*: exogenous CD33^M^ but not CD33^m^ co-immunoprecipitates with CLU in HEK293 cells.** Representative western blot for Figure 3b. *Top panel:* Representative western blots showing co-precipitation of CLU with CD33^M^ in HEK293 cells expressing CD33^M^ only (CD33^M^/CD33^M^ dimers), or in HEK293 cells expressing both CD33^M^ and CD33^m^ (CD33^M^/CD33^M^; CD33^M^*/C*D33^m^ or CD33^m^/CD33^m^ dimers) (CLU does not co-precipitate with CD33 in cells expressing only CD33^m^). *Top blot*: anti-HA pulldown probed with anti-CLU antibody. *Middle blot*: same IP products probed with anti-HA antibody. *Lower blot*: Protein input before precipitation probed with anti-CD33 antibody. This data is quantified in Figure 3b. In Figure 3B, the data for HEK293 cells expressing both CD33M and CD33m is displayed in the third column is the ratio of CLU immunoreactive band intensities versus just the CD33M band intensity. In the fourth column, the same CLU immunoreactive band intensity is expressed as the ratio of the cumulative band intensities as measured by HA immunoreactivity for both CD33m and CD33M isoforms.Each lane shown represents one independent biological replication of the respective condition.

**SUPPLEMENTARY DATA FIGURE 3C.**
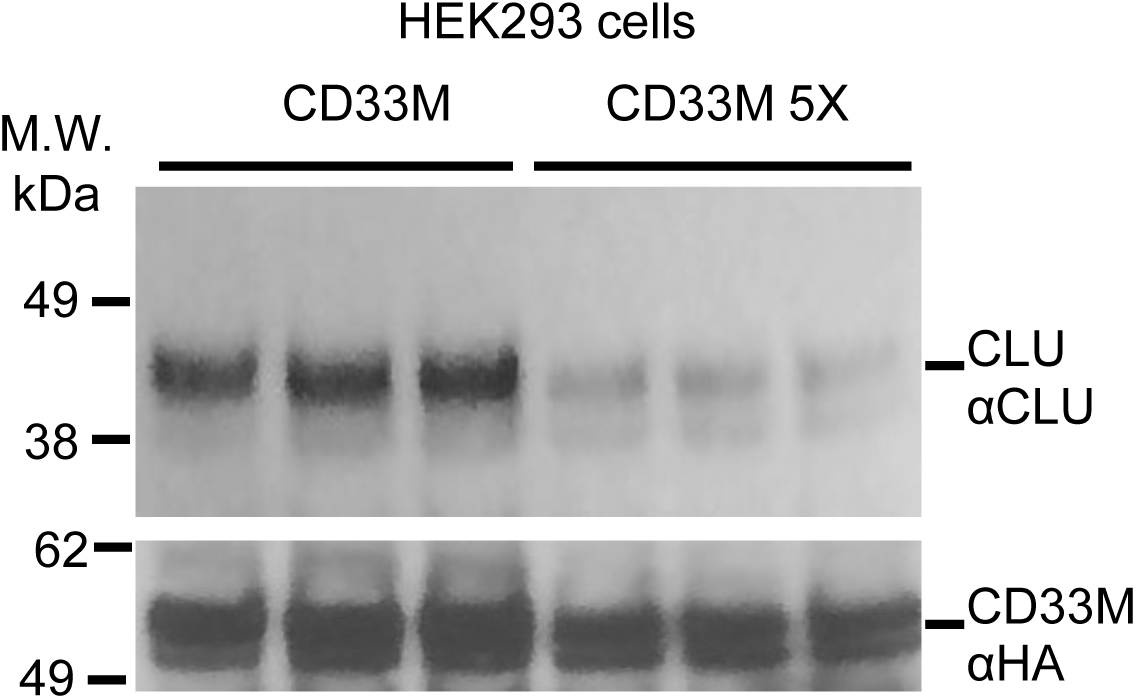
Dimerization of CD33 is required for CLU binding. Co-IP studies on lysates from HEK293 cells expressing similar quantities of CD33^M^ (*bottom panel*) reveal that much less CLU (*top panel*) is co-precipitated with the CD33^M^ 5X dimer interface mutant than with wild type CD33^M^. This is a representative blot for data for multiple biologica; replicates and is quantified in Figure 3c. Each lane shown represents one independent biological replication of the respective condition

**SUPPLEMENTARY DATA FIGURE 3D.**
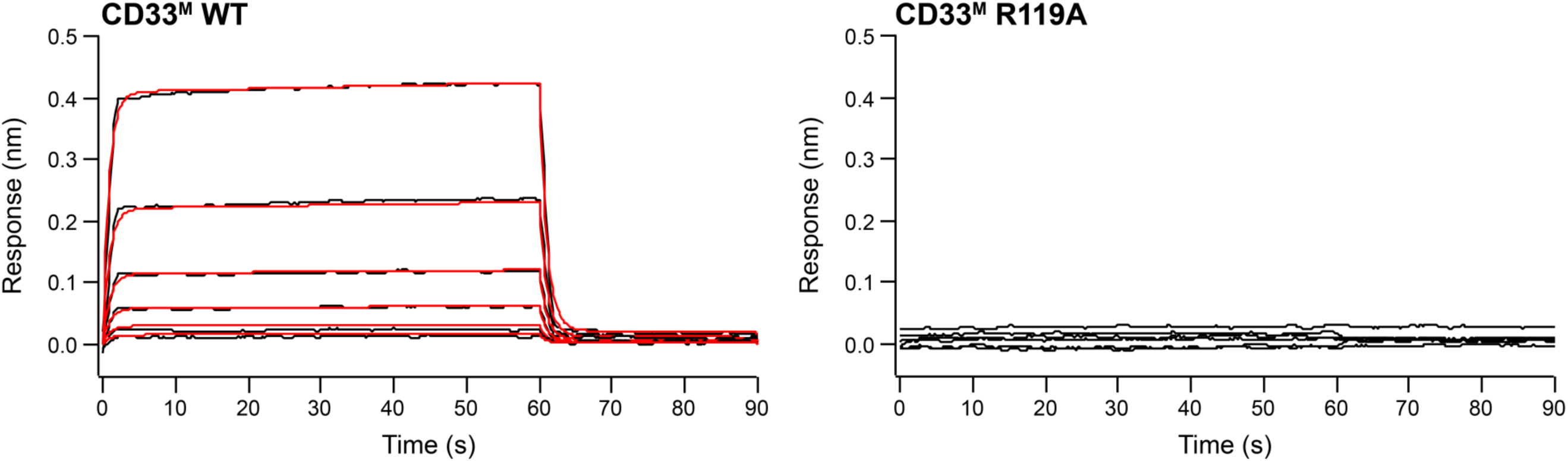
CD33^M^ ectodomain binds 3’-sialyllactose via the canonical R119. BLI response curves representing the association (0-60 s) and dissociation (60-90 s) of CD33^M^ ectodomain (ECD) WT (left panel) or R119A (right panel) in the concentration range of 15.6-250 µM with the 3’-sialyllactose-sp-biotin immobilized on the Super Streptavidin biosensors. Binding curves analysis using a 1:2 bivalent analyte model revealed that the dimer of CD33^M^ ECD WT binds to 3’-syalyllactose with dissociation constants 6.13 ± 0.25 mM (mean ± SEM). CD33^M^ ECD R119A showed no detectable binding in the same concentration range with WT.

**SUPPLEMENTARY DATA FIGURE 3E.**
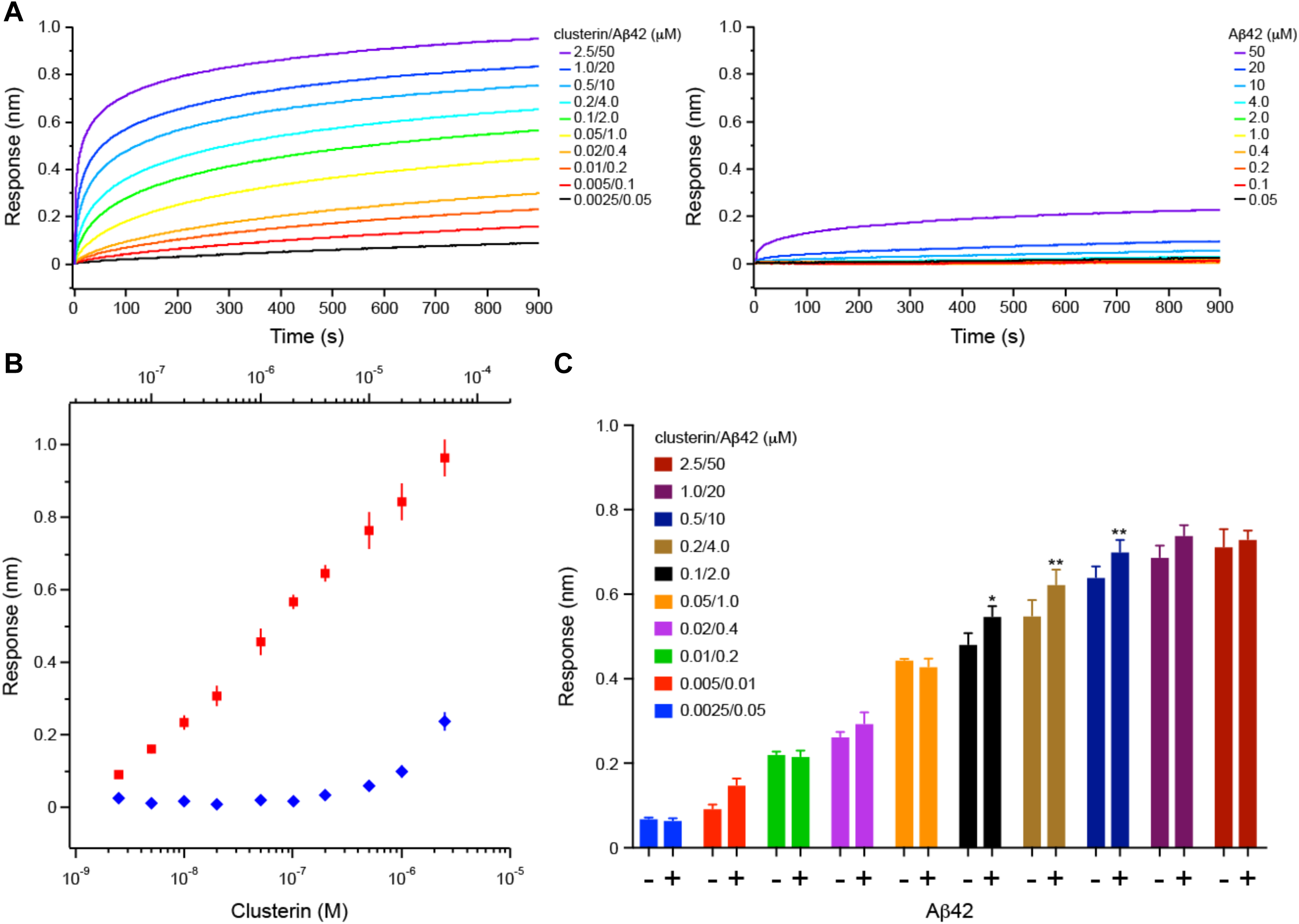
Biolayer Interferometry (BLI) studies on the clusterin/Aβ42 mixture binding to CD33^M^ ectodomain (ECD) **A.** Binding curves of clusterin/Aβ42 mixture (left panel) or Aβ42 (right panel). Each binding curve is the average of four independent measurements. Clusterin and Aβ42 concentrations are indicated. **B.** Steady-state BLI analysis of the clusterin/Aβ42 mixture (red square) or Aβ42 (blue diamond) binding to CD33^M^ ECD. Error bars represent SD. **C.** The bar chart of each data point presented in B. Error bars represent SD. Statistical analysis was performed using two-way ANOVA followed by Bonferroni’s multiple comparison test (n = 4, \**p*<0.05, \*\**p*<0.01 *vs* clusterin alone).

**SUPPLEMENTARY DATA FIGURE 3F.**
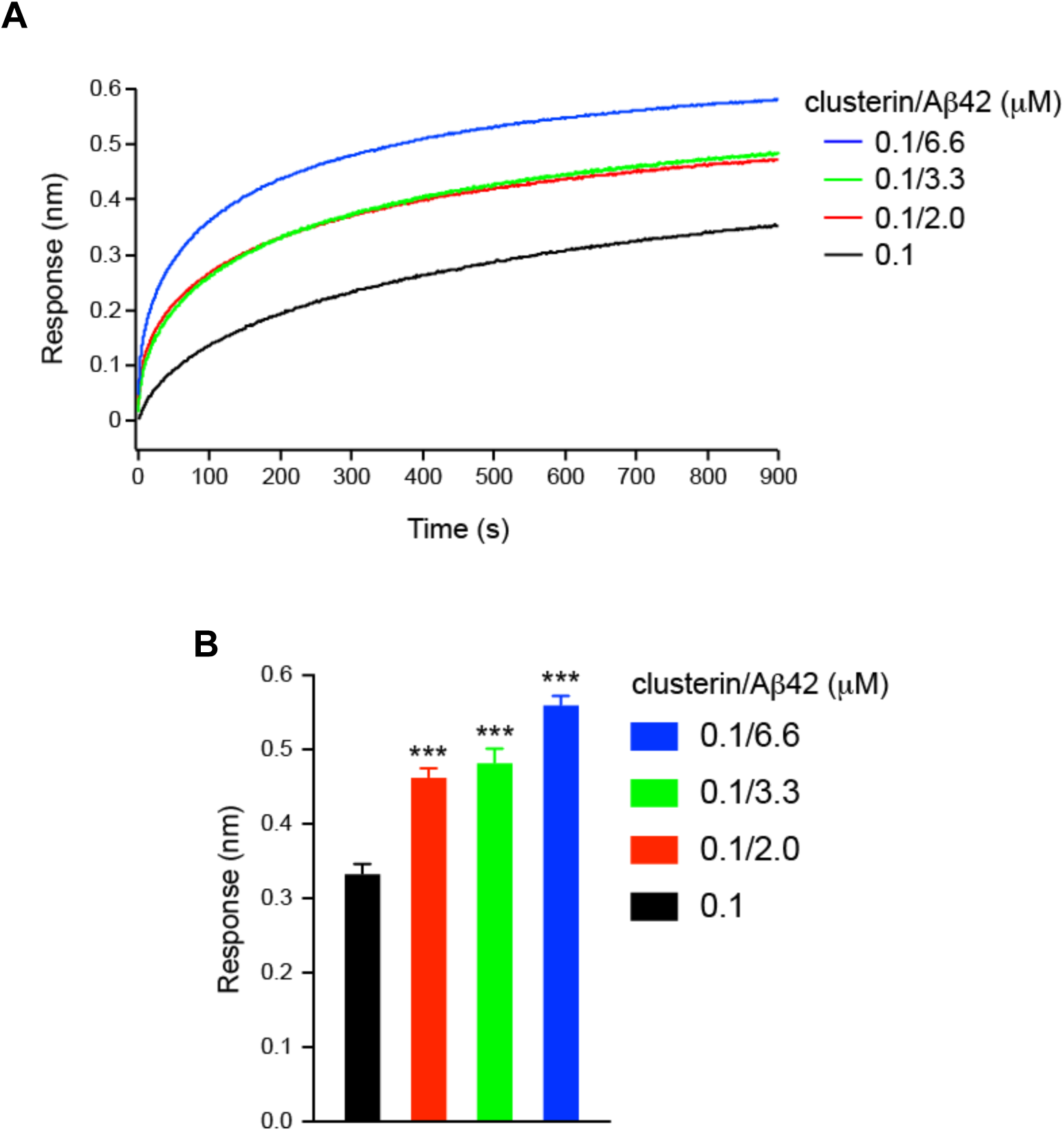
Clusterin/Aβ42 mixture bind to CD33^M^ ectodomain (ECD) in a clusterin/Aβ42 ratio dependent manner. **A.** Binding curves of clusterin (black) or clusterin/Aβ42 mixture (red, blue and green), respectively. Each binding curve is the average of four independent experiments. Clusterin and Aβ42 concentrations are indicated. **B.** Steady-state binding responses of clusterin or clusterin/Aβ42 mixture. Error bars represent SEM. Statistical analysis was performed using one-way ANOVA followed by Tukey’s test (n = 4, \*\*\**p*<0.001 *vs* clusterin alone).

**SUPPLEMENTARY DATA FIGURE 3G.**
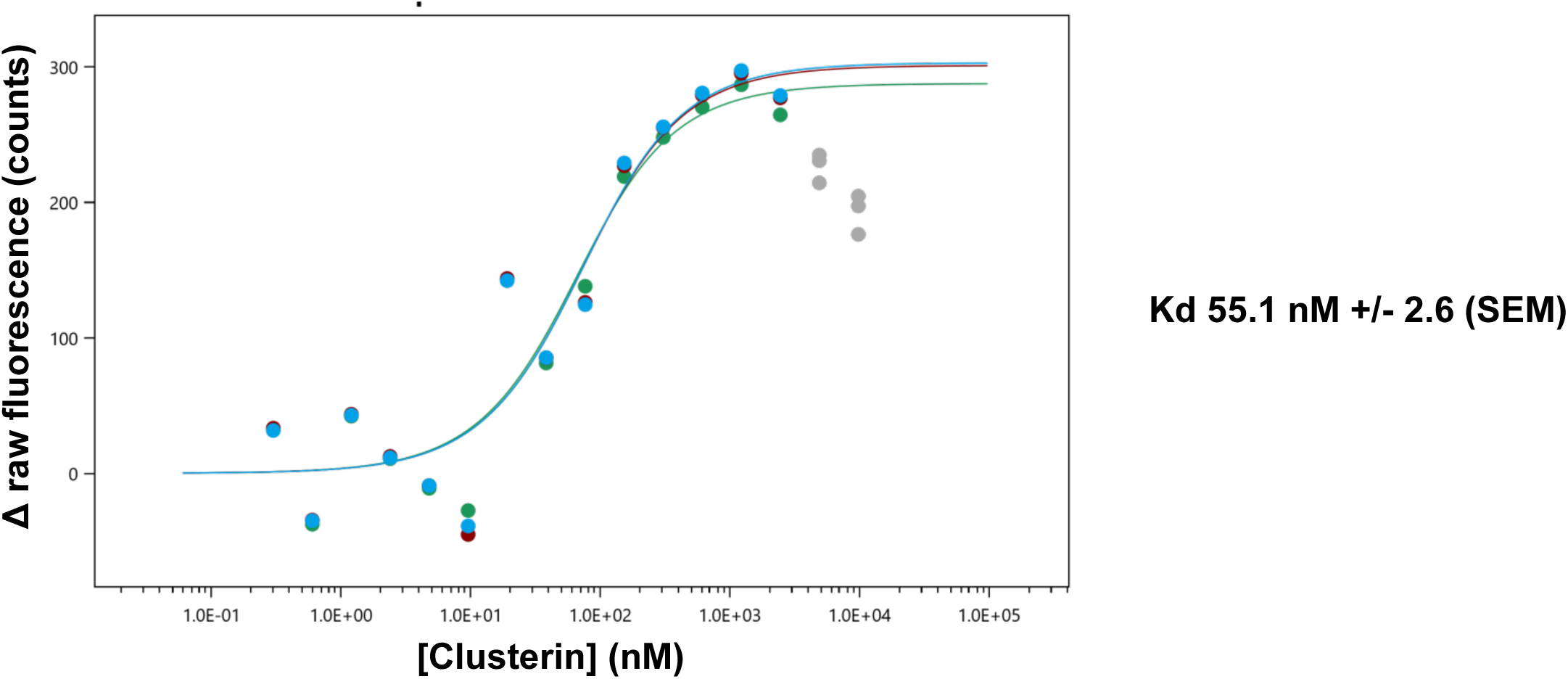
**Binding analysis by microscale thermophoresis measuring the change in the fluorescence intensity of fluorescently labeled deglycosylated CD33 ECD (initial fluorescence)**

**SUPPLEMENTARY DATA FIGURE 3H.**
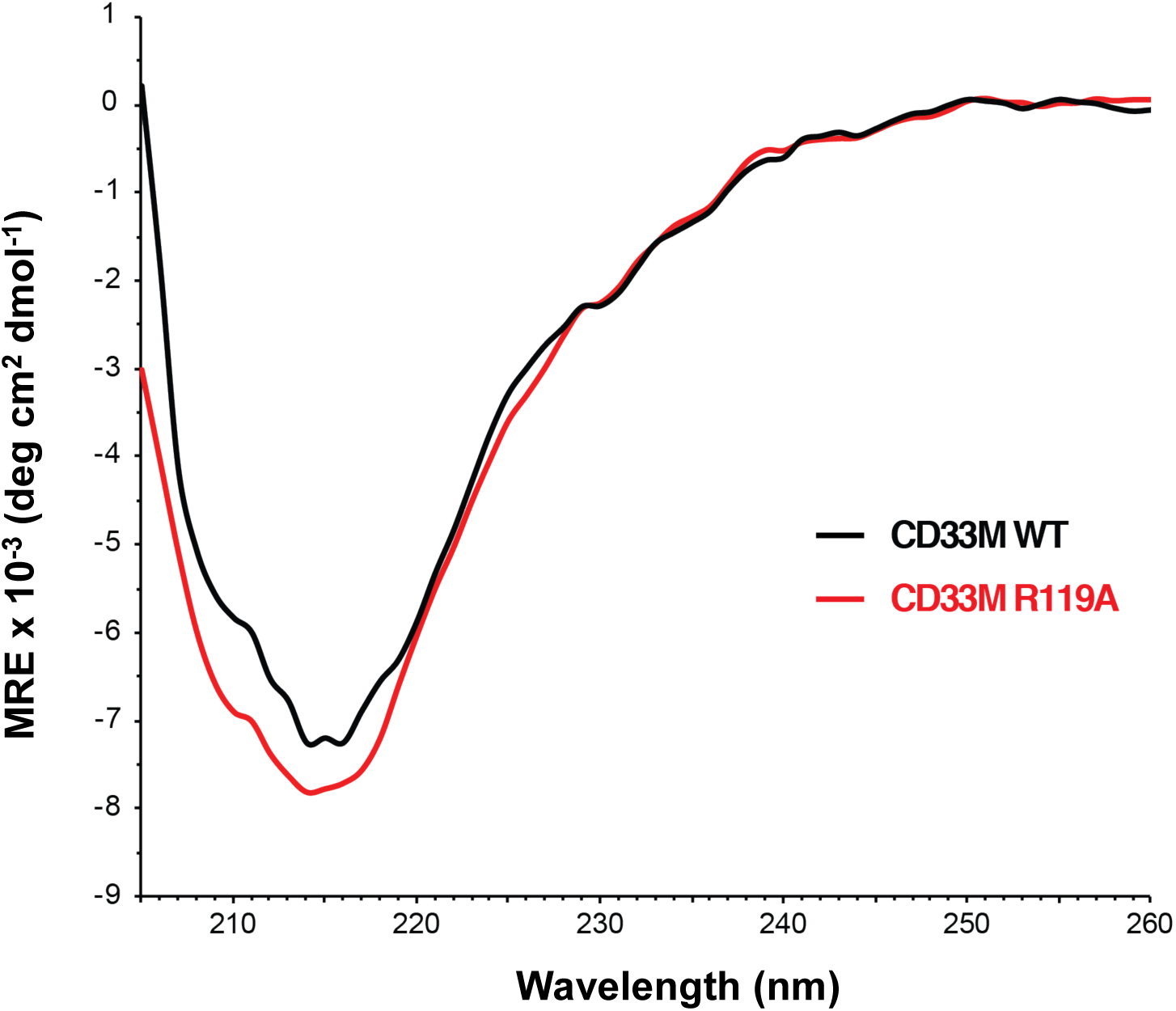
Far UV circular dichroism spectra for wild type CD33^M^ ectodomain (black) and R119A mutant CD33^M^ ectodomain (red) reveal similar folding states, indicating that the loss of sialic acid binding by the R119A protein is due to an authentic effect on ligand binding, rather than due to simple misfolding of the R119A mutant protein.. Spectra were recorded at 4°C with CD33 ECD at 0.2 mg/ml in 20 mM HEPES, 200 mM NaCl, pH7.5.

**SUPPLEMENTARY DATA FIGURE 4A.**
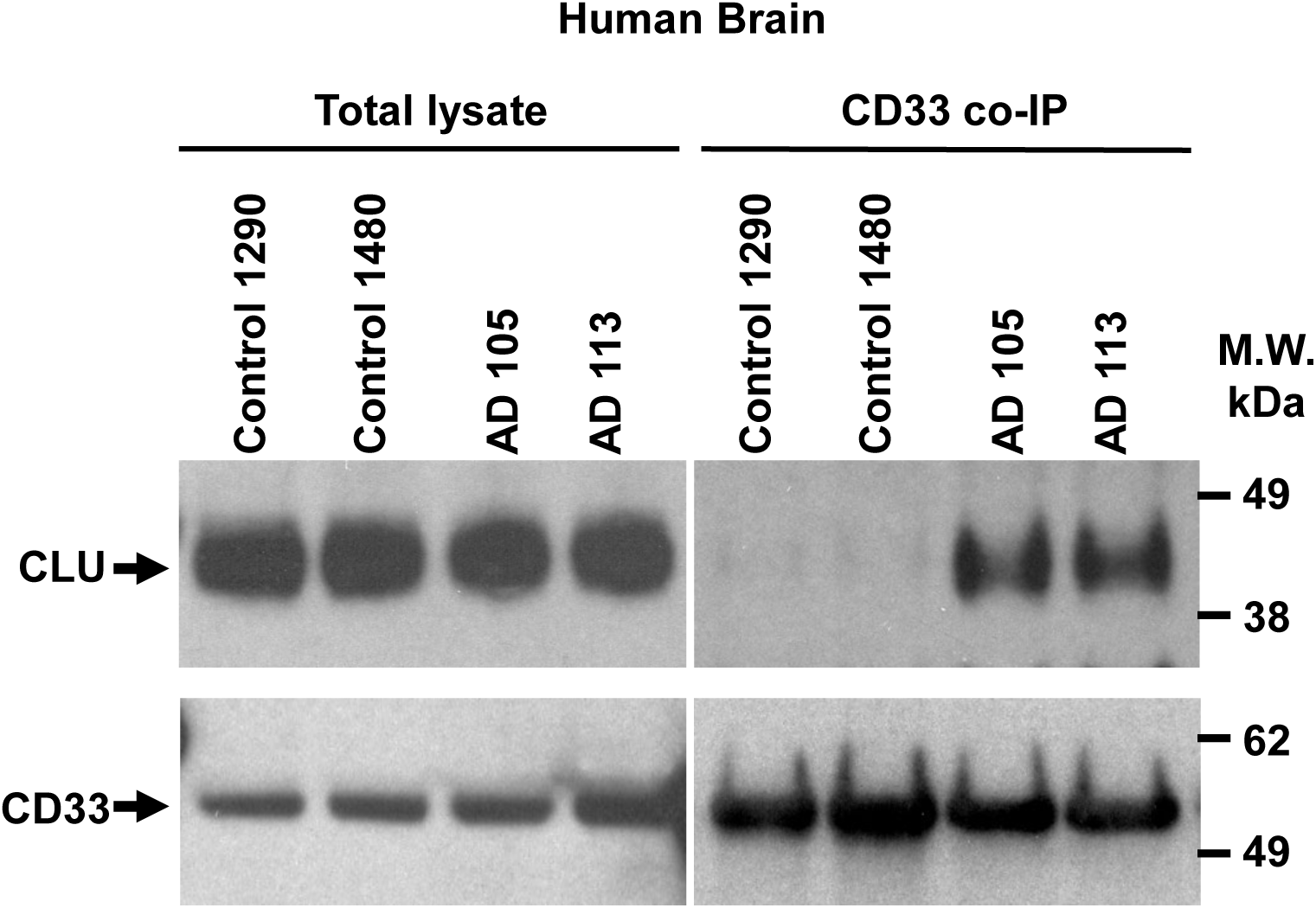
Representative Western blot for Figure 4c. Co-IP studies on lysates from HEK293 cells expressing similar quantities of CD33^M^ (*bottom panel*) reveal that much less CLU (*top panel*) is co-precipitated with the CD33^M^ 5X dimer interface mutant than with wild type CD33^M^. This data is quantified in Extended Data Figure 3D. *Right panel;* quantification. *** p<0.00001, error bars = mean ± SEM CD33^M^: 0.425±0.0073; CD33^M^ 5X: 0.012±0.006; n = 5 replications. Each lane represents one independent biological replication of the respective condition.

**SUPPLEMENTARY DATA FIGURE 4B.**
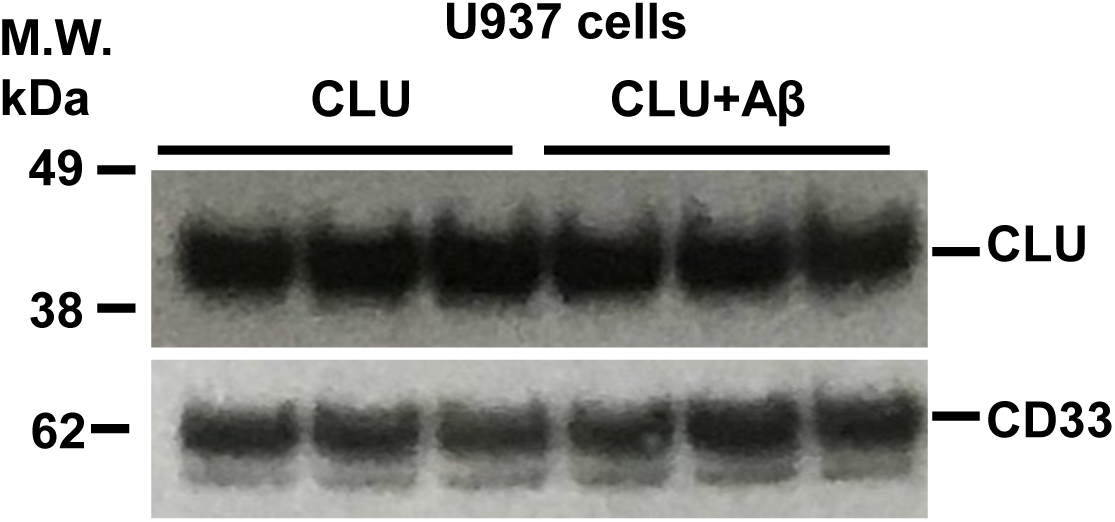
Representative Western blots for Figure 4d and 4e **showing that native human CLU alone, or complexed with Aβ oligomers also binds endogenous human CD33^M^ in U937 cells (right panel).** *Upper blot of each panel* is a representative western blots for anti-CD33 co-IP of CLU alone or CLU plus Aβ oligomers. *Lower blot of each panel* is a representative wester blot of CD33 co-IP products probed with anti-CD33 antibody to show similar CD33 IP recovery. Each lane represents one independent biological replication of the respective condition.

**SUPPLEMENTARY DATA FIGURE 5A.**
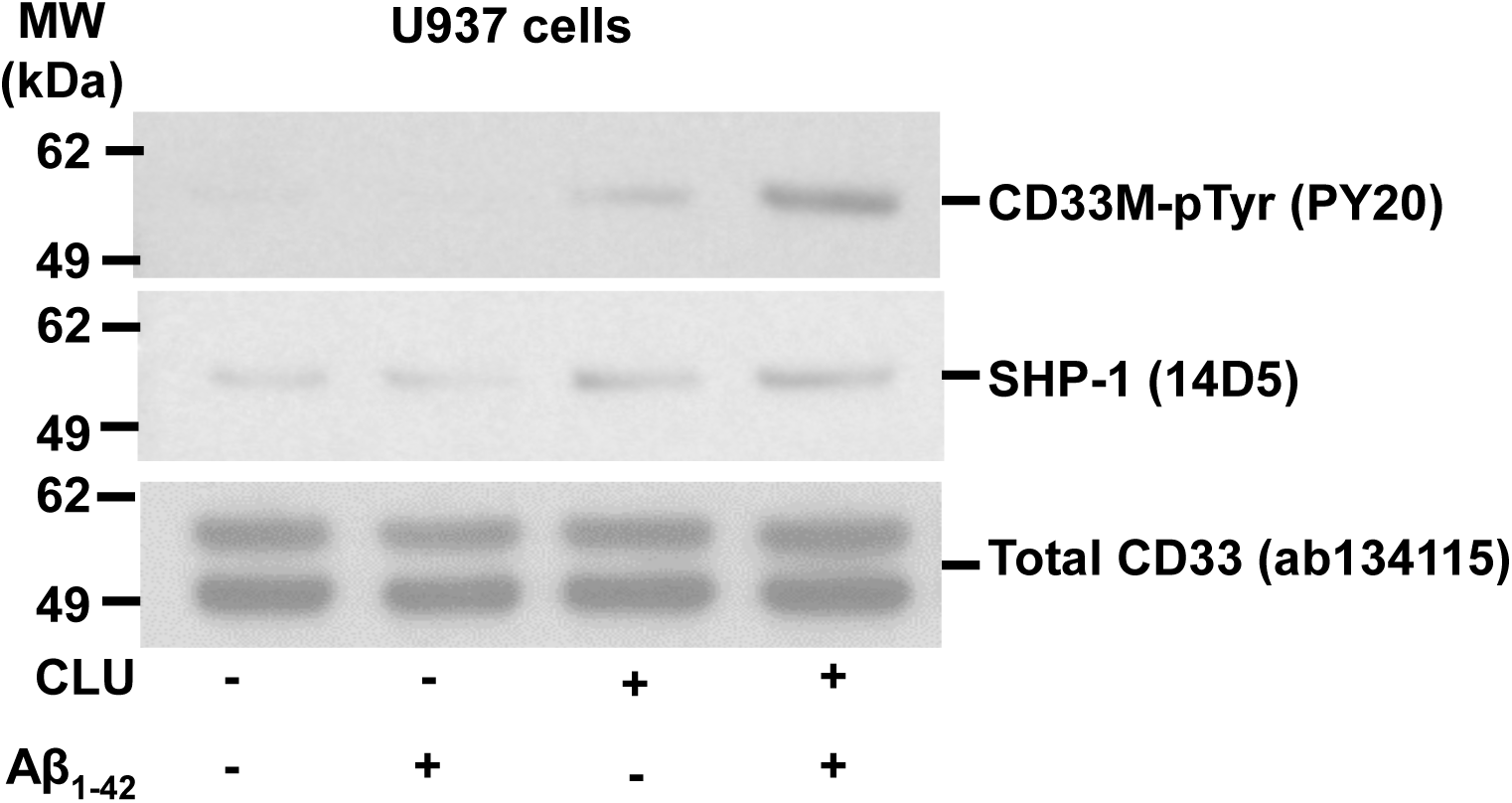
**Representative blot of n ≥ 3 independent biological replications for the graph in Figure 5a**, showing that human plasma CLU alone, and especially human CLU + Aβ oligomers induce ITIM tyrosine phosphorylation of endogenous CD33 *(top panel)* and recruitment of endogenous SHP-1 *(middle panel)* in human monocytic U937 cells.

**SUPPLEMENTARY DATA FIGURE 5B.**
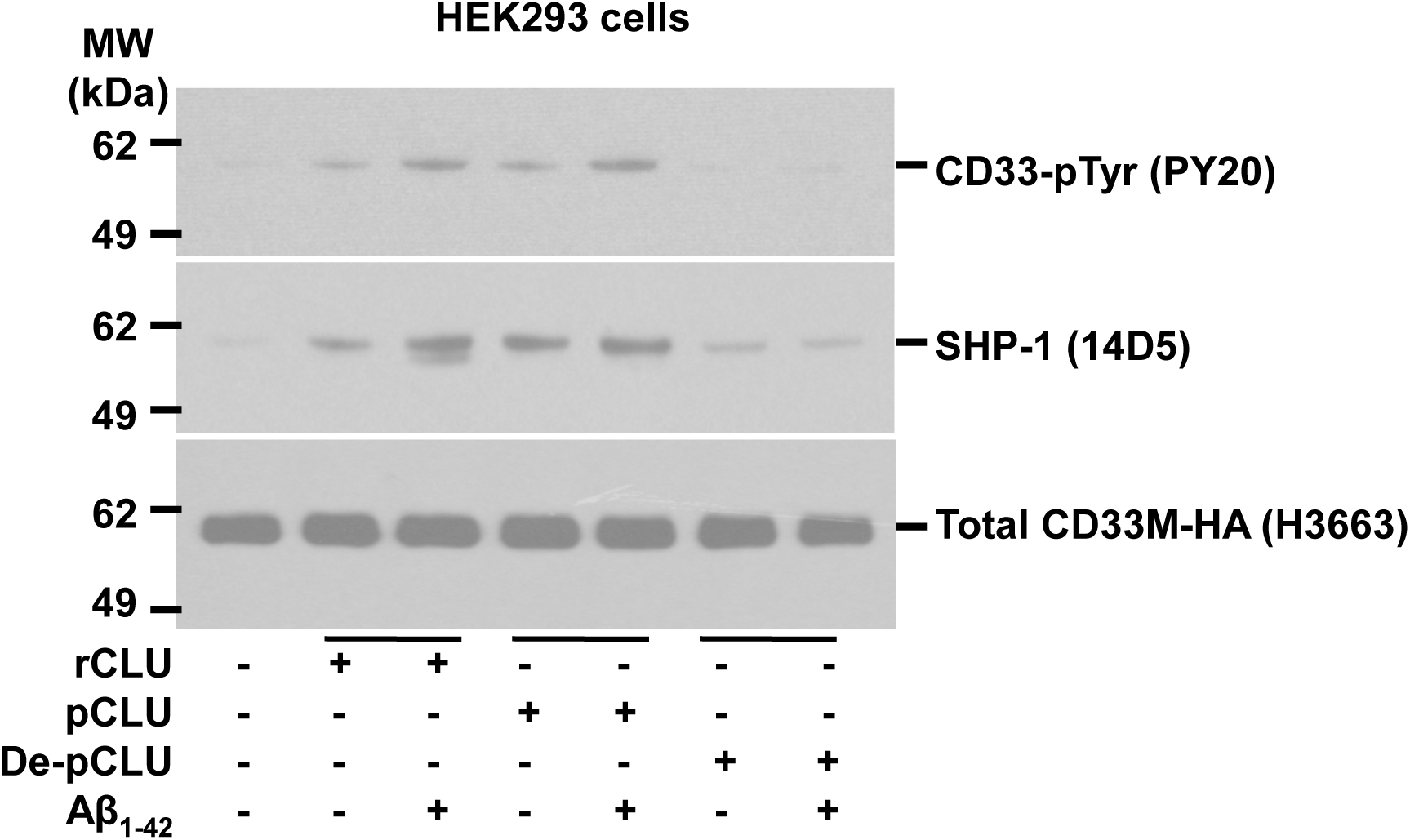
**Representative western blot of n = 3-4 independent biological replications with different cell lines for Figure 5b.** Recombinant human clusterin (rCLU) and native human plasma clusterin (pCLU), but not desialylated clusterin equivalently activate CD33M ITIM phosphorylation and SHP-1 recruitment in HEK293 cells. CD33M tyrosine phosphorylation (top panel) and SHP-1 recruitment (middle panel) were measured by Western blots band intensity ratios for pY20-phosphotyrosine immunoreactive CD33M versus total CD33M (bottom panel) by co-IP western blots..

**SUPPLEMENTARY DATA FIGURE 5C,D.**
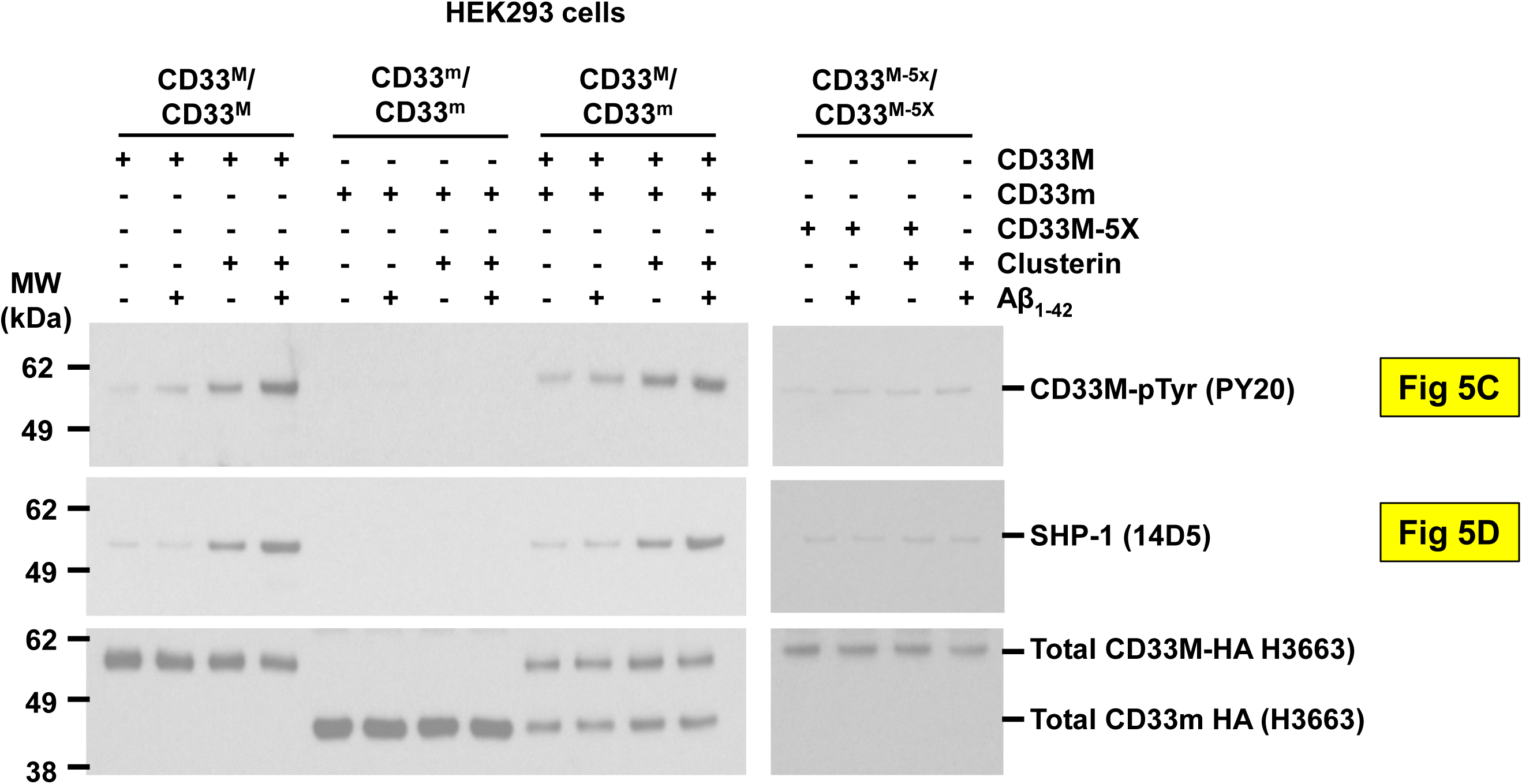
**Representative western blot from n = 3 independent biological replications for Figure 5c,d.** CLU + Aβ induces greater CD33 ITIM phosphorylation *(top panel)* and SHP-1 recruitment *(bottom panel)* than CLU alone, but only in CD33M expressing cells. Quantification of western blots of anti-CD33 immunoprecipitates probed with pY20 anti-phosphotyrosine, SHP-1 or anti-HA antibodies reveals that treatment with CLU alone, and especially treatment with CLU + Aβ oligomers, increases CD33 tyrosine phosphorylation and SHP-1 recruitment in cells expressing either CD33M only or CD33M plus CD33^m^. No CD33 phosphorylation or SHP-1 recruitment was observed in cells expressing only CD33m, and very little in CD33M 5X cells. n = 3 replications.

**SUPPLEMENTARY DATA FIGURE 5E.**
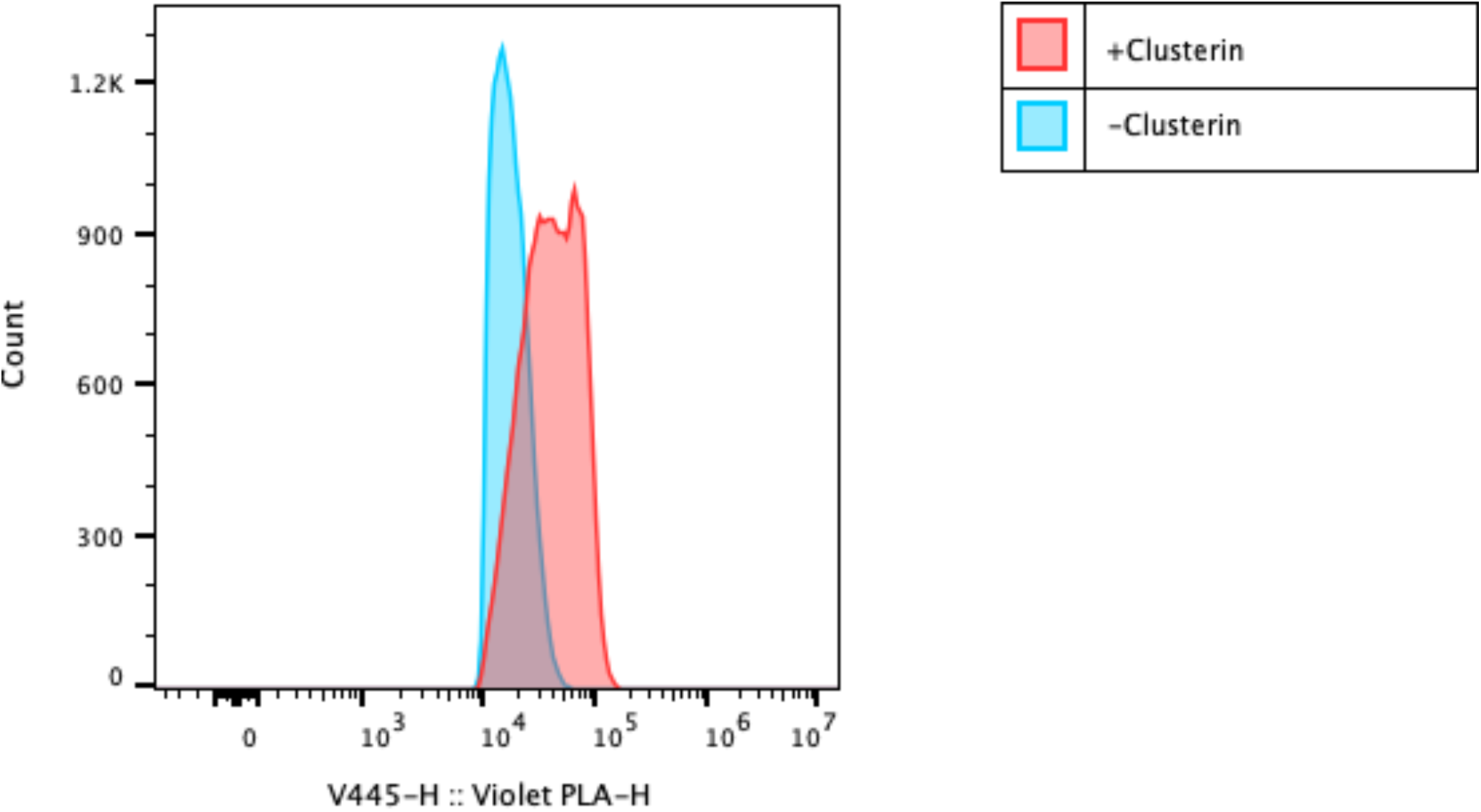
**Clusterin significantly increases SHP1 recruitment to CD33 from Figure 5e.** Example histogram showing the difference in CD33-SHP1 Flow PLA signal (violet 445) of human monocytes treated with CLU (red) and without CLU (blue) for 15 minutes

**SUPPLEMENTARY DATA FIGURE 5F.**
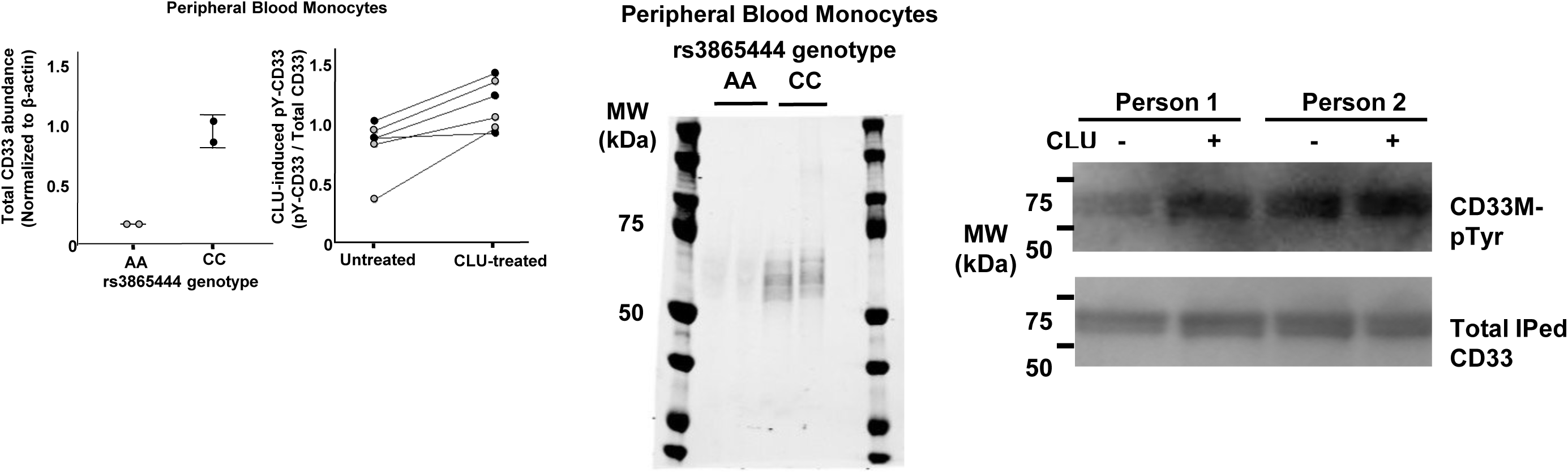
**Treatment of Human monocytes with 0.06 µM CLU for 15 min at 37°C increases tyrosine phosphorylation of the ITIM domain on endogenous CD33M.** Quantitative western blots of lysates for human blood monocytes reveal that monocytes from homozygous carriers of the rs3865444^AA^ reference /non-risk allele (gray dots) express less total CD33 and less CD33M than do homozygous carriers of the rs3865444^CC^ risk allele (black dots) (*Left panels*). Treatment of human monocytes with 0.06 µM CLU for 15 min at 37°C increases tyrosine phosphorylation of the ITIM domain on endogenous CD33M both in individuals homozygous for the rs3865444^CC^ risk allele (black dots) and in individuals homozygous for the rs3865444^AA^ reference /non-risk allele (gray dots) paired t-test, p=0.007 (2^nd^ *left panel).* NB: As demonstrated in the left panel, carriers of the rs3865444^CC^ risk allele express much more CD33M than do carriers of the rs3865444^AA^ reference allele. However, when equal quantities of CD33M protein are loaded onto these blots to facilitate comparison of basal and CLU-induced CD33 ITIM phosphorylation, the *specific activity* of CD33M signaling is the same. **Representative Western blots for Supplemental Figure 5F. Treatment of Human monocytes with 0.06 µM CLU for 15 min at 37°C increases tyrosine phosphorylation of the ITIM domain on endogenous CD33M.** *Middle panel*: Quantitative western blots of lysates for human blood monocytes reveal that monocytes from homozygous carriers of the rs3865444^AA^ reference express less total CD33 and less CD33M than do homozygous carriers of the rs3865444^CC^ risk allele. *Right panel*: Representative western blots showing that treatment of human monocytes with 0.06 µM CLU for 15 min at 37°C increases tyrosine phosphorylation of the ITIM domain on endogenous CD33M both in individuals homozygous for the rs3865444^CC^ risk allele (person 2) and in individuals homozygous for the rs3865444^AA^ reference /nonrisk allele (person 1). Representative western blots are available in Extended Data Figure 5C-G. NB: As demonstrated in the left panel, carriers of the rs3865444^CC^ risk allele express much more CD33M than do carriers of the rs3865444^AA^ reference allele. However, when equal quantities of CD33M protein are loaded (bottom right panel) onto these blots to facilitate comparison of basal and CLU-induced CD33 ITIM phosphorylation, the *specific activity* of CD33M signaling is the same.

**SUPPLEMENTARY DATA FIGURE 5G.**
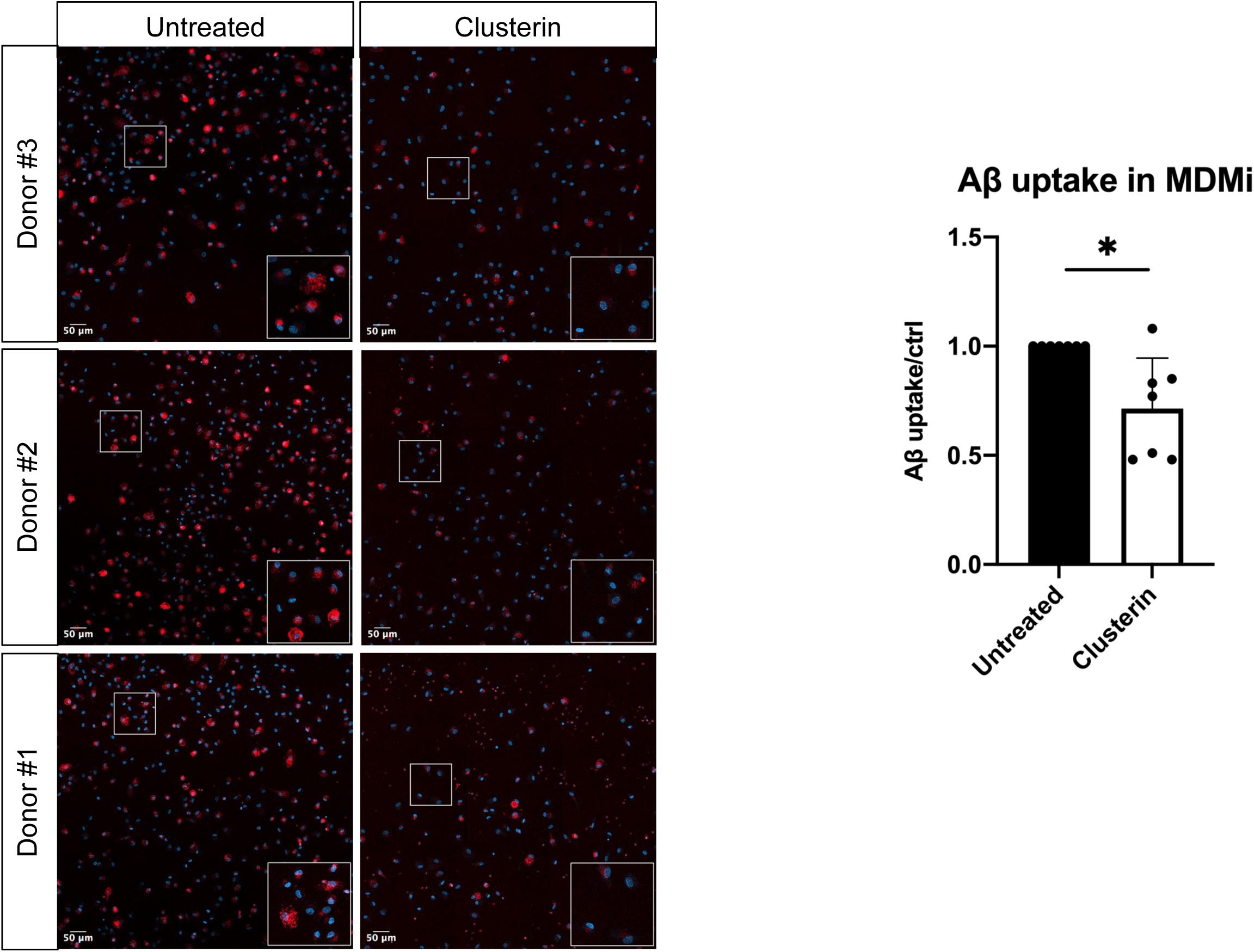
**Clusterin treatment reduces Aβ uptake in MDMi.** MDMi from 7 individuals were treated with 60nM clusterin for 15 minutes, then incubated with Aβ-647 and imaged for uptake. Representative images from 3 different donors are shown on the left. The bottom right corner of each image is a zoomed-in inlet of the white outlined region. Blue = DAPI. Red = Aβ. Scale bar is 50 µm. Quantification is shown in the bar graph on the right. *p<0.05. Wilcoxon test.

**SUPPLEMENTARY DATA FIGURE 6A.**
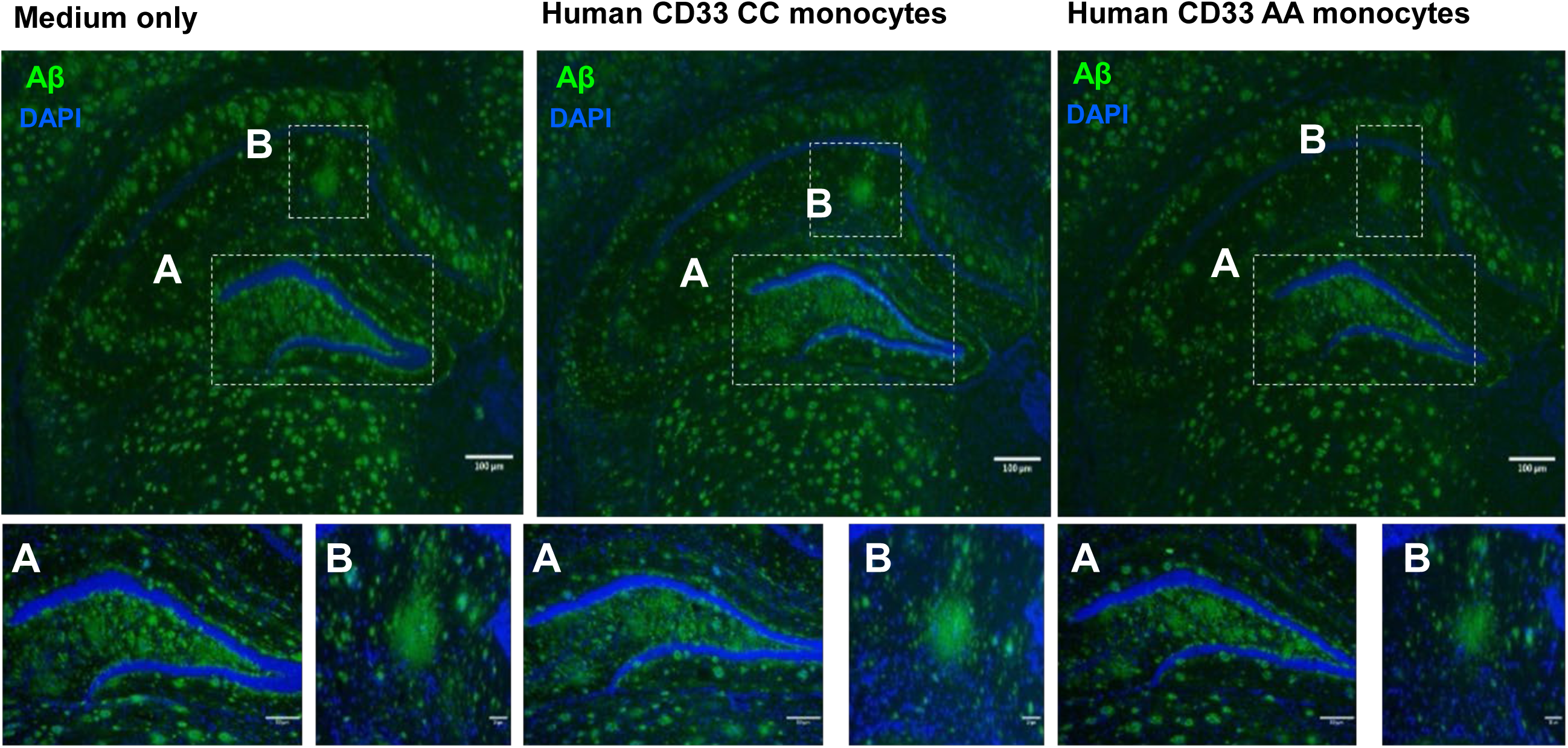
**Human monocytes from CD33 “CC” risk allele carriers** have reduced clearance of human amyloid plaques in *ex vivo* brain slices compared to monocytes from CD33 “AA” protective allele carriers. **Representative figures quantified in** Figure 6a of clearance of hippocampal Aβ plaques by human monocytes following incubation on serial sections of unfixed brain from 5XFAD mice. Immunofluorescent staining of hippocampal Aβ plaques (green) following 48h incubation with medium only with no cells (*left*) or with CC (risk allele) genotype monocytes (*middle*) or with AA (protective allele) monocytes (*right*). N = 10 independent replications). DAPI blue = nuclei.

**SUPPLEMENTARY DATA FIGURE 6B.**
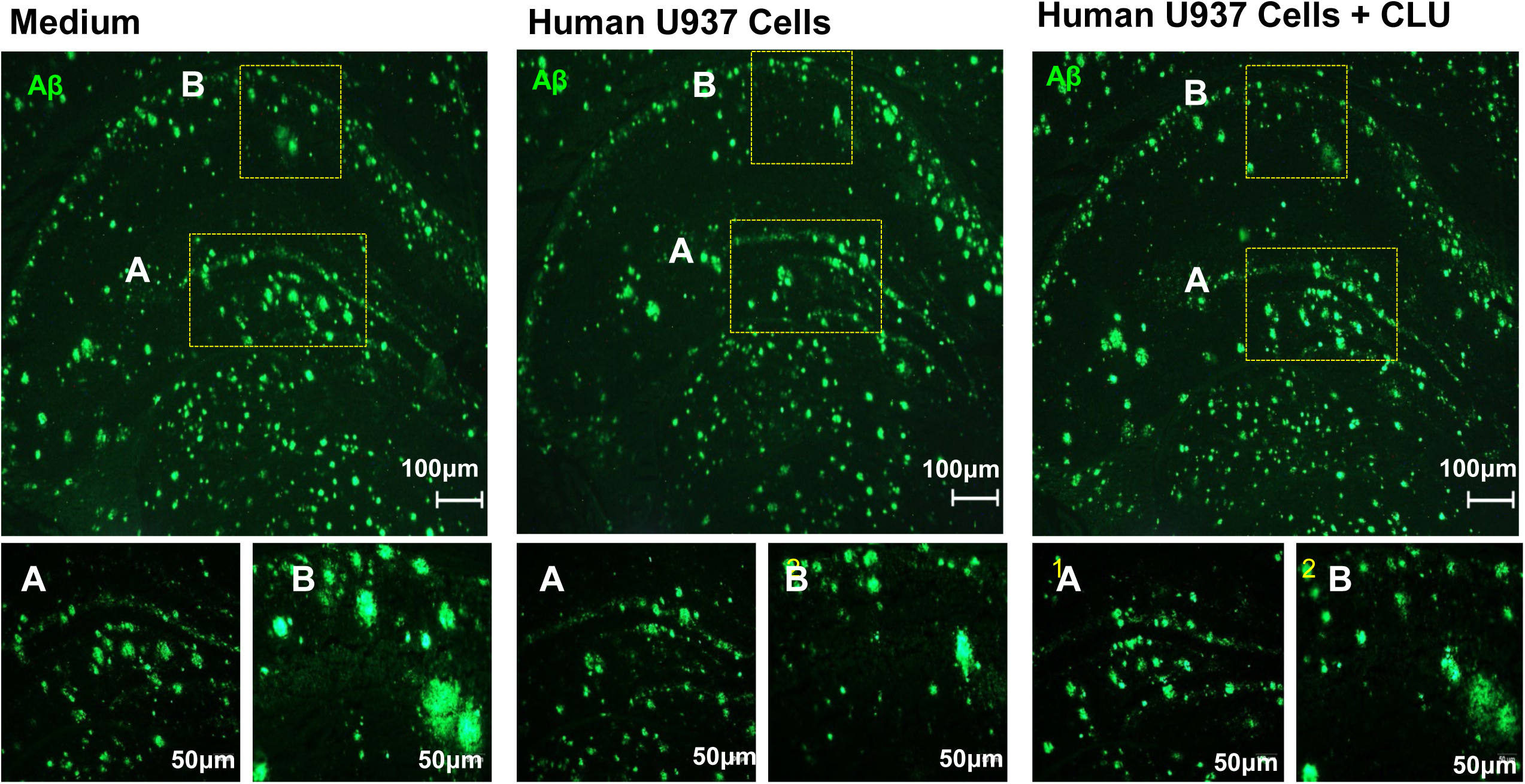
**Human U937 monocytic cells clear amyloid plaques from *ex vivo* brain slices, and this is impaired by CLU. Representative figures quantified in Figure 6b** of clearance of hippocampal Aβ plaques (green) by U937 cells following incubation on three serial sections of unfixed brain section of 5XFAD mice in the presence/ absence of CLU. Immunofluorescent staining of hippocampal Aβ plaques following 48h incubation with only medium (upper left figure) or with U937 cells alone (upper middle figure) or with U937 cells + CLU (upper right figure). Yellow boxes highlight two regions (A: 10X and B: 20X) that are shown at higher magnification in the corresponding images below the main figure). Scale bars: 100 μm in I and A; and 50 μm in B.

**SUPPLEMENTARY DATA FIGURE 6C.**
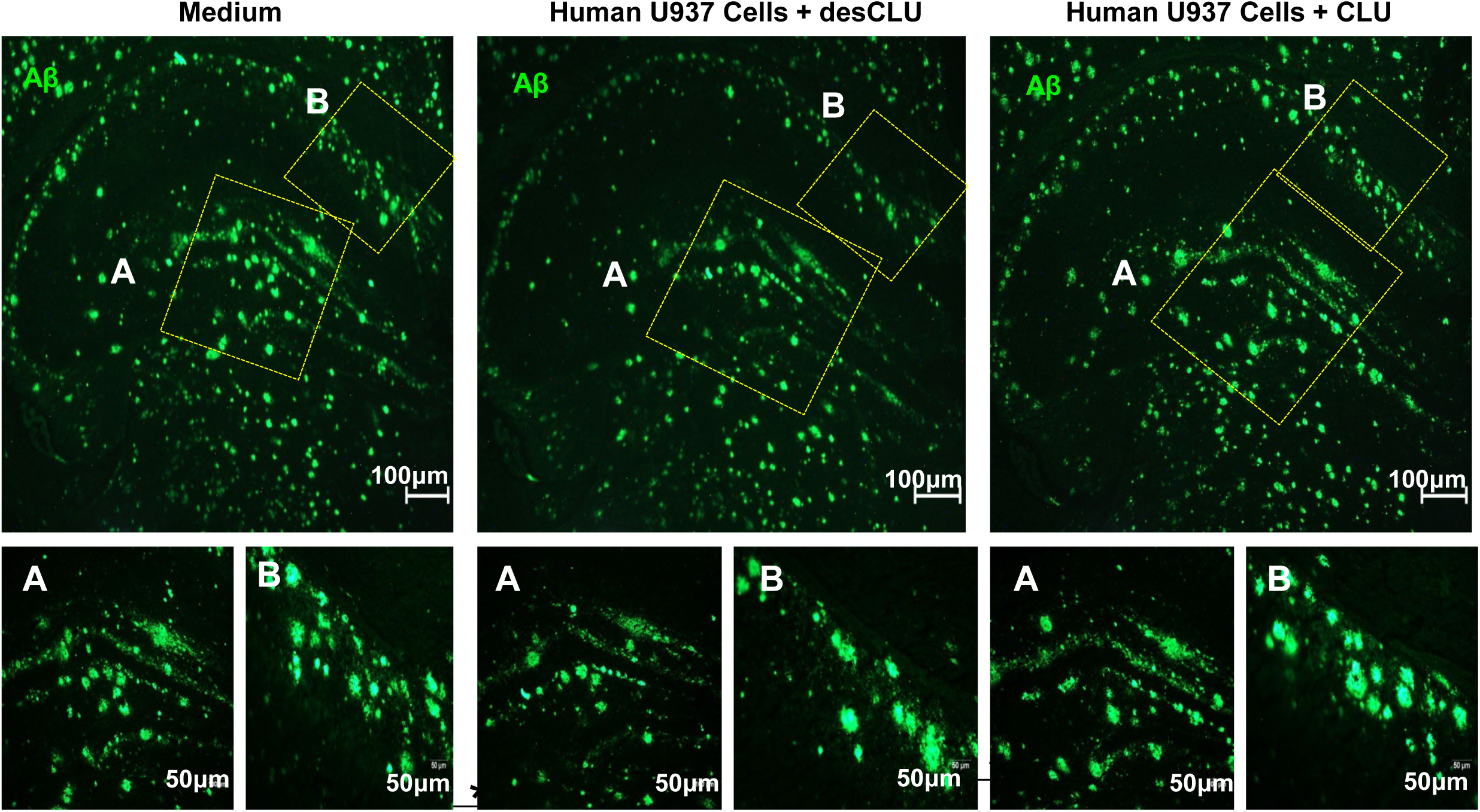
**Compared to CLU, desCLU does not downregulate clearance of Aβ plaques by human U937 monocytic cells. Representative figures quantified in Figure 6c** of clearance of hippocampal Aβ plaques (green) by U937 cells following incubation on serial sections of unfixed brain from 5XFAD mice in the presence of CLU or desCLU. Immunofluorescent staining of hippocampal Aβ plaques following 48h incubation with medium only but no cells (*left*) or with U937 cells + desCLU (*middle*) or with U937 cells + CLU (*right*). Small figure magnifications of regions of interest: A: x10; B: x20. Scale bars: 100 μm in I and A; and 50 μm in B. n = 10 replications

**SUPPLEMENTARY DATA FIGURE 6D.**
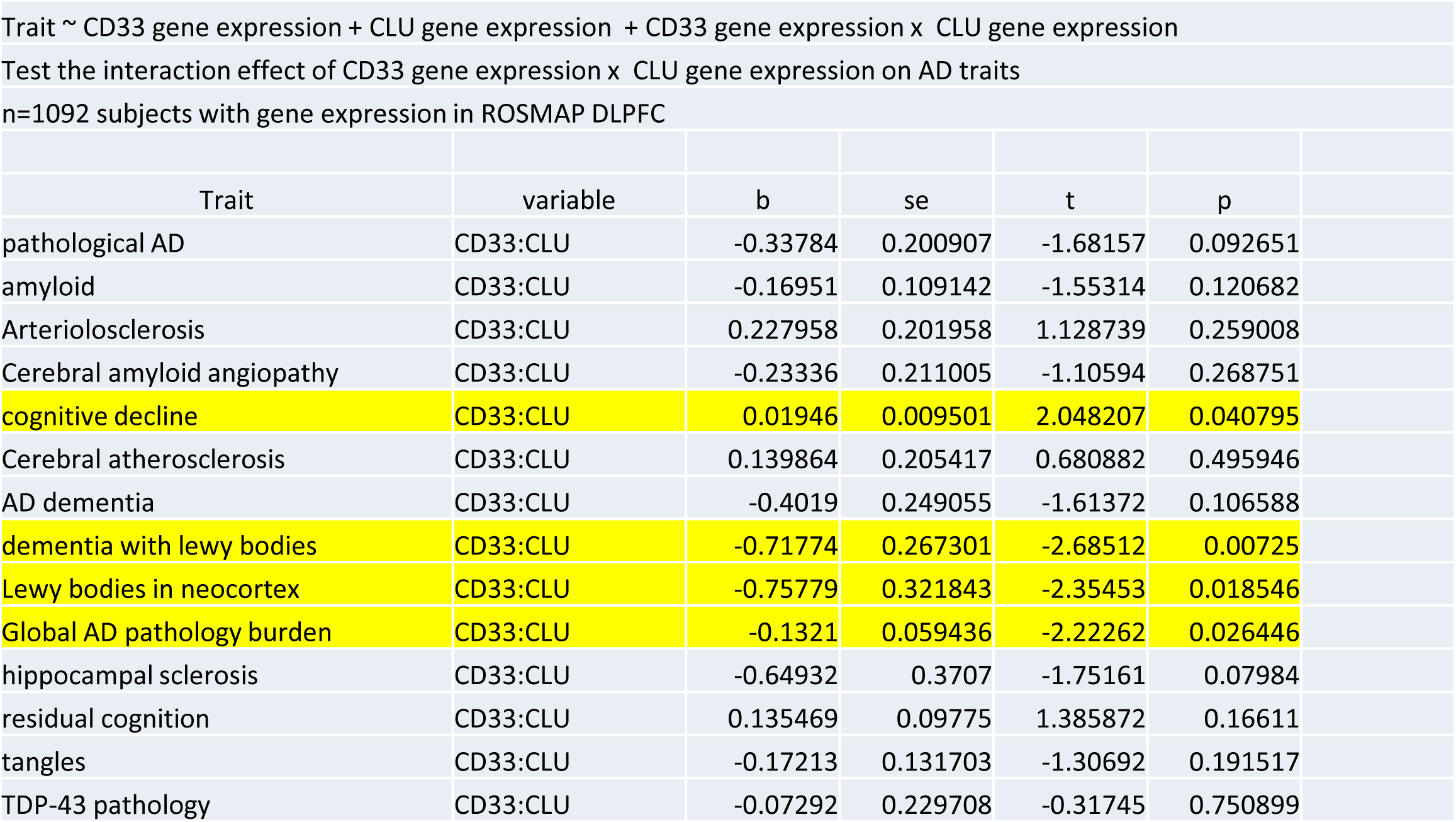

**SUPPLEMENTARY DATA FIGURE 6E.**
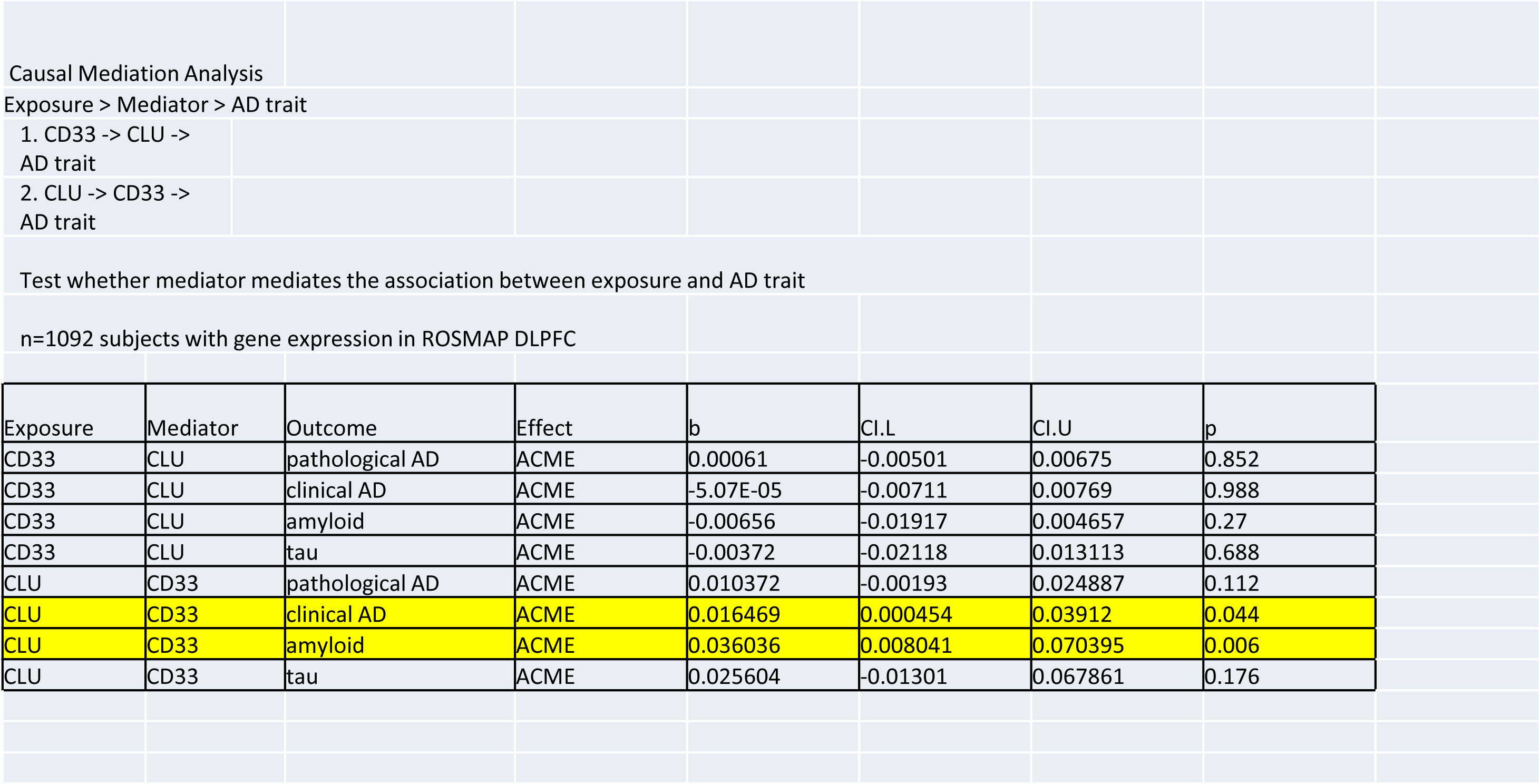

**SUPPLEMENTARY DATA FIGURE 6F.**
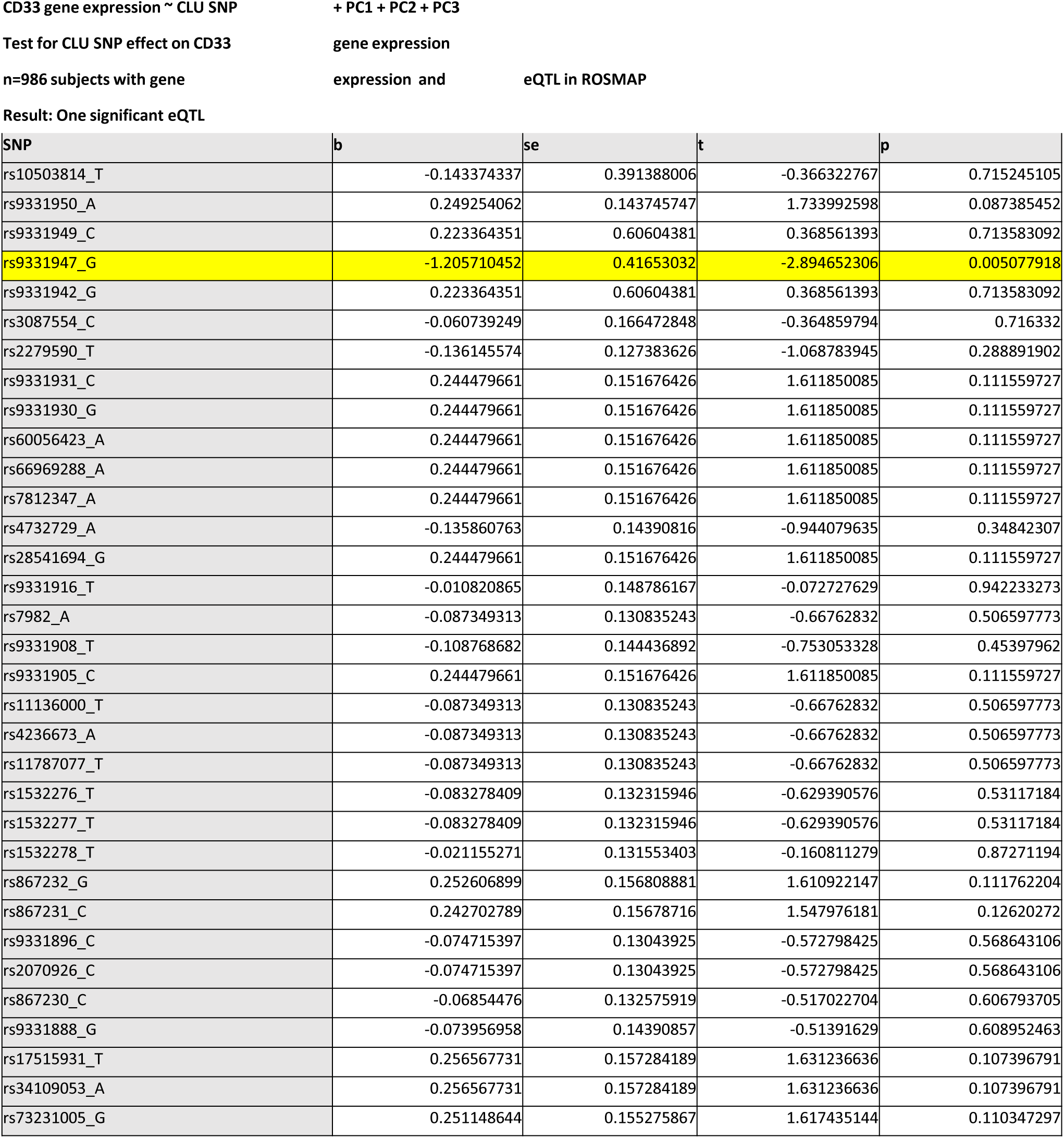

**SUPPLEMENTARY DATA FIGURE 6G.**
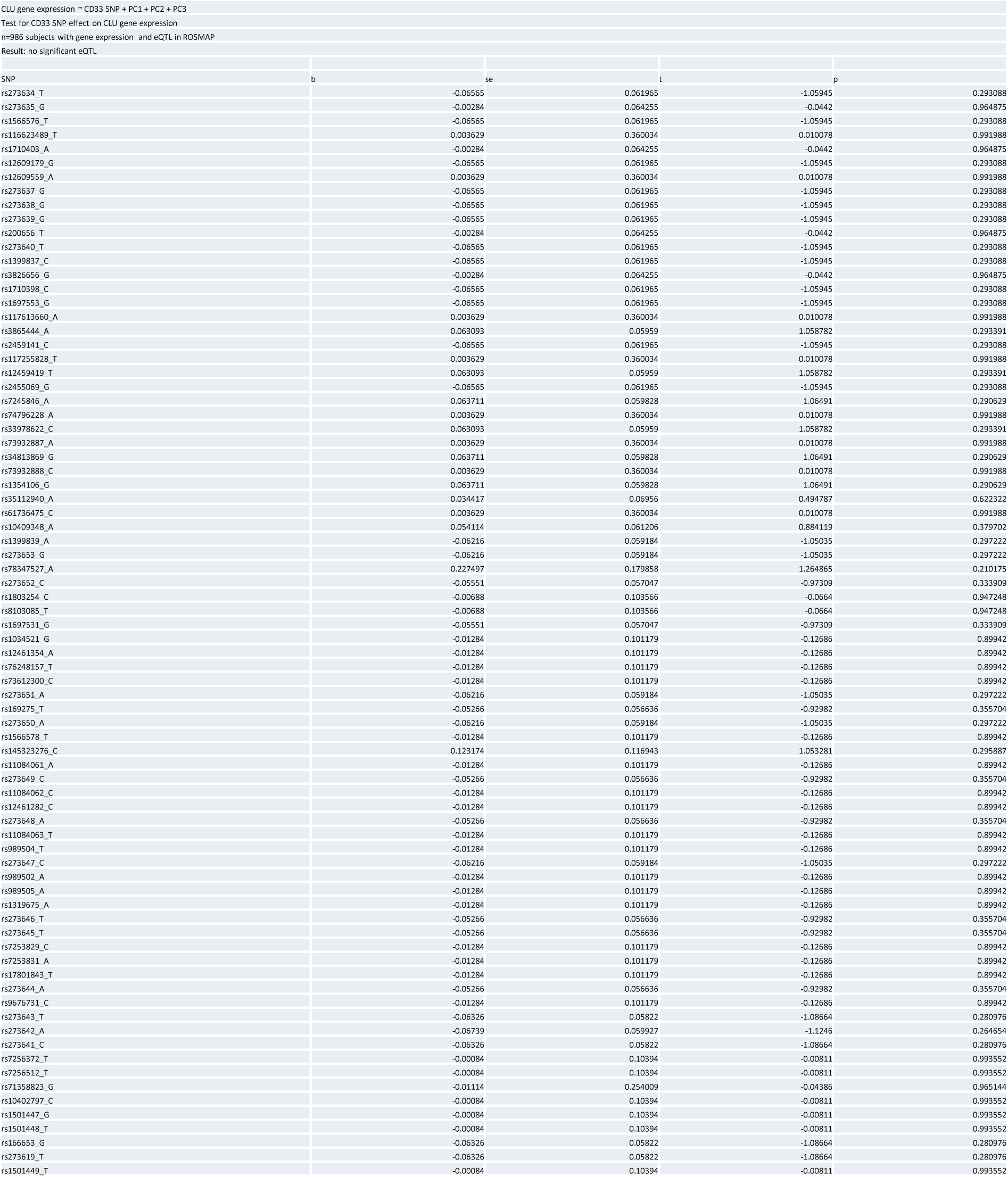

**SUPPLEMENTARY DATA FIGURE 6H.**
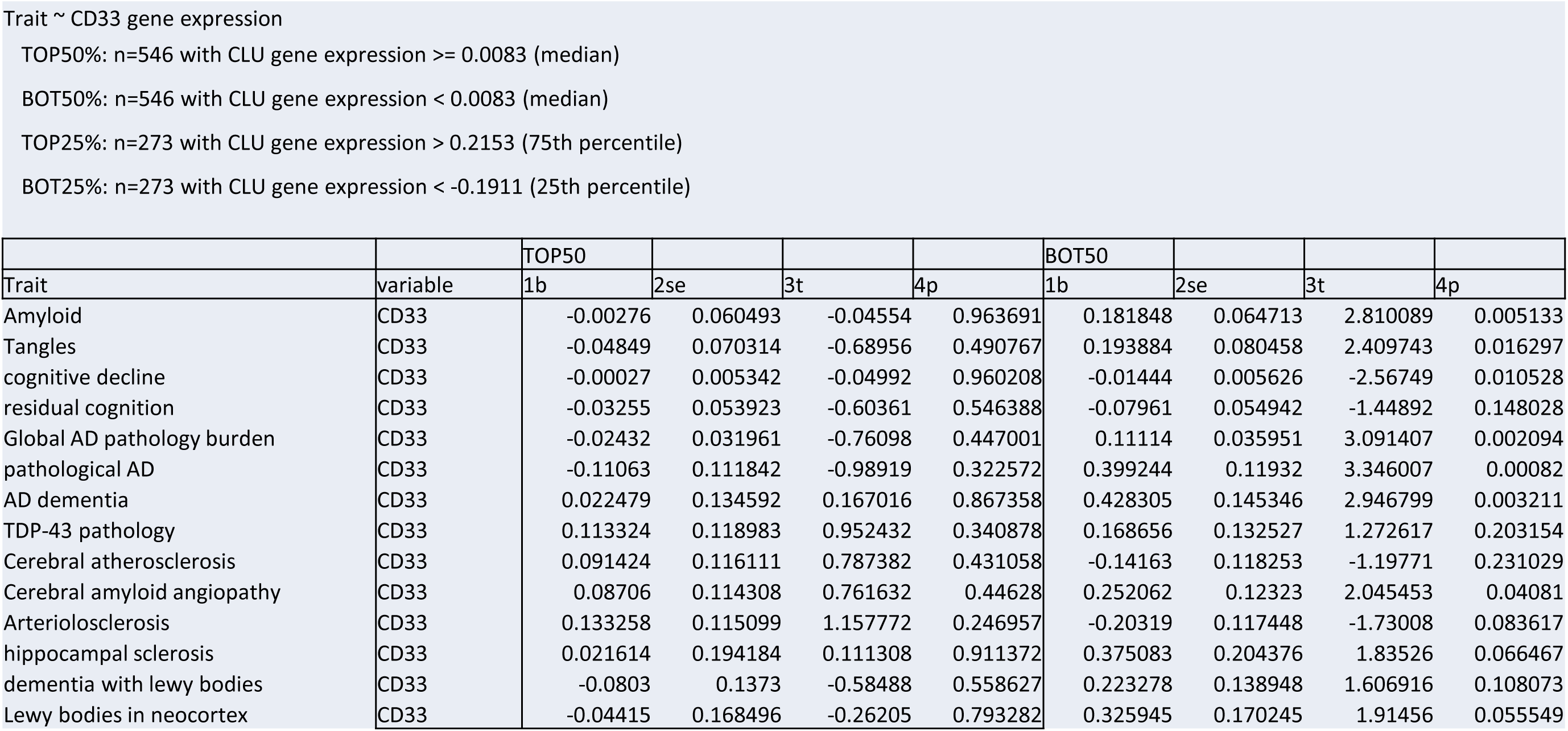

## Notes

### Competing Interest Statement

The authors have declared no competing interest.

